# Whole-genome characterization of *Plasmodium falciparum* population structure and drug resistance across Uganda and neighboring countries

**DOI:** 10.64898/2026.09.27.754784

**Authors:** Shahiid Kiyaga, Monica Mbabazi, Thomas Katairo, Victor Asua, Francis D. Semakuba, Bienvenu Nsengimaana, Stephen Tukwasibwe, Innocent Wiringilimaana, Jackie Nakasaanya, Jerry Mulondo, Eric Watyekele, Catherine Maiteki-Sebuguzi, David Patrick Kateete, Joyce Nakatumba Nabende, Daudi Jjingo, Gerald Mboowa, Charles Batte, Samuel L. Nsobya, Moses R. Kamya, Bosco B. Agaba, Isaac Ssewanyana, Andrés Aranda-Diaz, Fehintola Victoria Ajogbasile, Milya Davlieva, Grant Dorsey, Philip J. Rosenthal, Melissa D. Conrad, Bryan Greenhouse, Nicholas Hathaway, Jessica Briggs

## Abstract

Uganda has one of the world’s highest malaria burdens, but parasite population structure and connectivity have not been characterized nationally with whole-genome data. We analyzed 249 monoclonal *Plasmodium falciparum* genomes from 26 sentinel sites across Uganda in 2023 and compared them with publicly available genomes from five neighboring countries. Genetic diversity was high and differentiation low in Uganda, but identity-by-descent detected limited structure. This was most pronounced in the low-transmission southwest, where parasites shared ancestry and recent relatedness with isolates from western Tanzania. Selection scans identified signals at loci associated with resistance to artemisinins and their partner drugs, and to chloroquine and antifolates. The strongest signal was at PfPX1 where the PIN haplotype (L1222P, M1701I, D1705N) reached 70% prevalence nationwide, but selection was carried by PIN haplotypes that had additionally acquired D384A, largely confined to northern and eastern Uganda. A second substitution at the same codon, D384Y, formed a haplotype shared across the Tanzanian border. PfKelch13 variants were partitioned across multiple backgrounds, consistent with repeated independent emergence, and C469Y was enriched on the PfPX1 D384A-PIN background. These genomes provide a 2023 baseline for tracking potential mediators of drug resistance and support coordinated genomic surveillance across Uganda and its neighbors.

## Introduction

*Plasmodium falciparum* malaria is concentrated in sub-Saharan Africa, where over 95% of cases and deaths occur^1^. Uganda is among the highest burden countries globally, with spatially heterogeneous transmission ranging from hypo-endemic conditions in the southwestern highlands to intense, perennial transmission across northern lowlands^2–5^. Whether this ecological heterogeneity produces a correspondingly structured parasite population, and how transmission, human movement, and geography shape parasite connectivity in Uganda, remains poorly resolved. These questions are increasingly relevant given the rise of artemisinin partial resistance (ART-R) in Africa. *P. falciparum* kelch13 (*pfkelch13*) propeller domain mutations mediating ART-R, first documented in Southeast Asia, have since emerged independently in East Africa^6^. These mutations are now established across multiple regions in Uganda, with marked spatial heterogeneity in specific mutations and their frequencies^7–11^. Given that Uganda borders high-transmission countries with differing ART-R profiles, and because sub-national tailoring of interventions is increasingly being implemented, mapping parasite gene flow within Uganda and across its borders is a priority^10,12^.

Resolving parasite connectivity in Uganda requires genome-wide data. Central to this is characterization of identity-by-descent (IBD), genomic segments shared between parasites through inheritance from a recent common ancestor, which resolves ancestry, quantifies population connectivity, and detects recent selective sweeps^13–15^. Whole-genome sequencing (WGS) complements targeted sequencing by capturing genome-wide variation, enabling *de novo* discovery of selective sweeps anywhere in the genome, characterization of genome-wide linkage disequilibrium (LD) decay, and reconstruction of finer scale population structure and IBD^16^. Within Uganda, prior population genomic studies using microsatellites^17,18^ or targeted amplicons^3,19^ have found substantial parasite diversity and high multiplicity of infection, but low differentiation between sites, consistent with extensive gene flow. However, the extent of population structure and connectivity within Uganda and across its borders has not been assessed at genome-wide resolution, and no comprehensive country-wide WGS dataset exists.

The benefits of genome-wide resolution also extend to drug resistance surveillance. Targeted monitoring of resistance markers in *pfkelch13* (PF3D7_1343700; ART-R), *pfcrt* (PF3D7_0709000; 4-aminoquinoline resistance), *pfmdr1* (PF3D7_0523000; ACT partner-drug response), and *pfdhfr/pfdhps* (PF3D7_0417200/PF3D7_0810800; sulfadoxine–pyrimethamine resistance), can guide treatment policy, but cannot evaluate selection at unmonitored loci or reconstruct ancestry. Selection scans can discover emerging resistance *de novo*: for example, the recently identified PfPX1 (PF3D7_0720700; phosphoinositide-binding protein) PIN haplotype (L1222P, M1701I, D1705N), detected through genome-wide signals of directional selection, has spread widely across Uganda and is associated with reduced susceptibility to both dihydroartemisinin and lumefantrine^20,21^. IBD can additionally distinguish independent local emergence from dissemination of shared haplotypes^7–9,12,14,22^. This distinction bears on intervention strategies: regional dissemination of a lineage implicates parasite movement and importation, whereas repeated independent emergence implicates local drug pressure or other ecological factors^23–25^. Notably, these approaches have not been extended across borders to understand how Uganda’s resistance landscape relates to that of its neighbors at genome-wide resolution^8,10,12,19,26^.

Here we present the first country-wide whole genome analysis of Ugandan *P. falciparum*, to determine: (1) parasite structure and connectivity within Uganda and across its borders, and (2) which drug resistance alleles have evidence of selection, and whether they spread as shared lineages or emerged on distinct genetic backgrounds. We analyzed 249 high-confidence genomes, collected in 2023, from 26 sentinel sites across six regions of Uganda. We established a diversity and population structure baseline, mapped sub-national and cross-border connectivity by IBD, and contrasted selection and pairwise haplotype sharing at key drug resistance loci. Together, these analyses provide the first comprehensive, whole-genome view of *P. falciparum* population structure, connectivity, and drug resistance spread across Uganda and its borders, to inform malaria control strategies.

## Results

### Study cohort and dataset assembly

We analyzed a subset of *P. falciparum*-positive dried blood spot samples collected in 2023 from 26 sentinel health facilities across Uganda. Samples were selected for whole genome sequencing in two stages: an initial random selection of monoclonal samples with parasitemia >500 parasites per microliter, followed by targeted enrichment for *pfkelch13* mutant samples. Of these 340 samples, four were removed for low coverage, and four were excluded for low within-host fixation index (*F*_*ws*_ <0.95), consistent with mixed or complex infections that could bias haplotype-based and relatedness analyses. The remaining 332 samples were down-sampled to 8–10 per sentinel site to achieve geographic representation and to correct for oversampling of *pfkelch13* mutants. Down-sampling was informed by the empiric *pfkelch13* mutant frequency at each site^10^. The final nationally representative dataset comprised 249 whole-genome sequences from 26 sentinel sites distributed across six geographic regions: North (10 sites, n = 93), East (5 sites, n = 47), Southwest (6 sites, n = 60), Northeast (2 sites, n = 19), Northwest (2 sites, n = 20), and West (1 site, n = 10) (**Supplementary Figure 1**).

### Genome-wide genetic diversity was uniformly high across Ugandan regions

Analysis of 28,926 high-quality biallelic SNPs revealed high genetic diversity across all regions (**Supplementary Table 1**). The Western region is included in the table for completeness but is excluded from the comparisons below owing to its small sample size (n = 10), which limits the reliability of these estimates. Nucleotide diversity (*π*) was essentially invariant across regions, ranging from 2.30×10^−4^ in the Northeast and Northwest to 2.39×10^−4^ in the Southwest, a range of < 4%. Expected heterozygosity (*H*_*e*_) showed the same pattern, ranging from 0.143 in the Northwest to 0.154 in the Southwest, a range of < 8%, with regions broadly ordered as for *π*. LD showed similar homogeneity, rapidly decaying to background within ~5–10 kb genome-wide (**Supplementary Figures 2 and 3**), with only minor regional variation in decay rate.

### Identity-by-descent revealed geographically structured parasite relatedness across Uganda

Pairwise IBD was estimated using hmmibd-rs^27^ with default settings. IBD analysis revealed substantial fine-scale heterogeneity in parasite relatedness across Uganda (**Figure 1a)**. Across 30,876 pairwise comparisons, 0.376% were related at a pairwise IBD fraction ≥ 1/32, corresponding to an expected most recent common ancestor within approximately five generations, with a mean genome-wide IBD fraction of 0.00471 and a maximum of 1. The mean within-region IBD ranged from 0.004 (East) to 0.013 (Southwest). The Southwest region was the only region to have any isolates with complete IBD, with two clusters of two isolates each. The Southwest had the highest proportion of related pairs at 4.12%, followed by the Northeast (2.34%), West (2.22%), Northwest (0.53%), North (0.37%), and East (0.37%).

**Figure 1.**
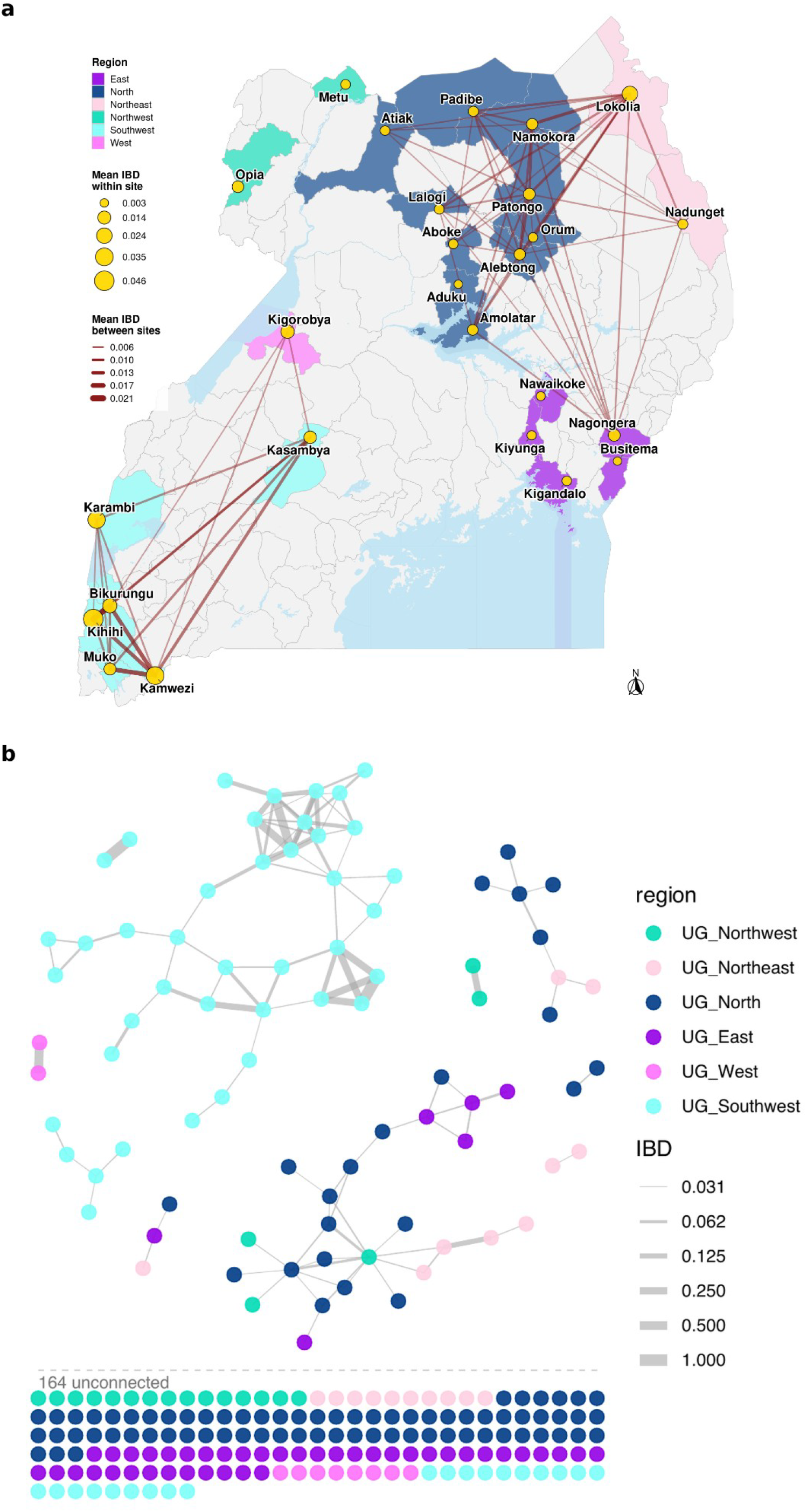
Spatial distribution of site-level IBD and between-site parasite relatedness across Uganda. **(a)** Map of *P. falciparum* sampling sites across Uganda, coloured by geographic region and overlaid on district boundaries. Yellow circles represent study sites, with circle size proportional to the mean IBD within a site. Brown connecting lines indicate between-site parasite relatedness, with line thickness proportional to the mean IBD between two sites. Lines are shown only for site pairs with mean IBD ≥ 0.006. **(b)** Sample-level IBD network coloured by region, with 116 edges and 76 nodes connecting pairs of isolates with IBD fraction ≥ 0.03125 (generations to most recent common ancestor (MRCA) ≤ 5).

Within sites, IBD-based connectivity varied geographically: mean within-site IBD sharing was 0.6% (0.2-2.2%, 19 sites) in the Northern and Eastern regions versus 2.3% (1.0-4.5%, 7 sites) in the lower transmission Southwest and West. Between-site IBD sharing within a sub-region averaged 0.4% (0.1-1.1%) in the Northern and Eastern regions versus 0.8% (0.4-2.0%) in the Southwest and West regions (**Figure 1a**). Sample-level IBD networks (**Figure 1b**) showed that related parasites (76/249) clustered geographically, forming localized sub-networks within the broader national population. While IBD networks annotated with drug resistance genotypes (**Supplementary Figures 4-6**) showed that related parasites frequently carried identical resistance mutations within the same region, most parasites carrying the same mutations, even in the same region, were typically unrelated at the genome-wide level.

### Broad-scale population structure was weak but detectable

Population structure was assessed using Uniform Manifold Approximation and Projection (UMAP) dimensionality reduction^28,29^, ancestry and admixture inference via sparse non-negative matrix factorization (sNMF), and pairwise Jost’s D differentiation. To place Ugandan population structure in a regional context, we incorporated publicly available WGS data for seven neighboring populations: Kinshasa - Democratic Republic of the Congo (DRC) (n = 59, 2022), Eastern Kenya (n = 32, 2019–2020), Western Kenya (n = 20 2015, n = 10 2018), Western Tanzania (n = 33, 2022), Eastern Tanzania (n = 30, 2019), Ethiopia (n = 16, 2013–2015), and Sudan (n = 6, 2015–2017). For each country, we prioritized the most recently collected samples; however, if recent sampling was restricted to sites distant from Uganda, we included older samples from more proximate locations (e.g., Western Kenya). We also compared our current samples with past Uganda samples (n=19 from 2006-2019, n=55 from 2020-2022). The accessions and metadata for these other samples can be found in **Supplementary Table 2**.

UMAP of LD-pruned genotypes using 30 principal component analysis (PCA) components (corresponding to 10% of the variation of the input data) showed that the majority of Ugandan *P. falciparum* isolates from different regions overlapped in low-dimensional space, forming a single, dense cluster with limited geographic stratification when analyzed in isolation (**Supplementary Figure 7**). Parasites from all six regions occupied a largely continuous UMAP distribution, indicating substantial genetic overlap across Uganda. Most Northern, Northwestern, Eastern, and Western parasites were extensively intermixed. Southwestern parasites exhibited the clearest regional differentiation, although some overlap with parasites from other regions remained. When the Uganda datasets were analyzed together with the neighboring country comparators, Southwestern Ugandan parasites clustered closest to Western Tanzanian parasites, and Northern/Northwestern isolates occupied intermediate positions between Ugandan and neighboring regional clusters (**Figure 2a**). Western Kenyan isolates clustered primarily alongside Eastern Ugandan isolates, and Eastern Kenyan isolates clustered primarily with Eastern Tanzanian isolates. Three of the Sudanese isolates clustered with Uganda, while the remaining three clustered with the Ethiopian isolates, as did a single Northern Ugandan isolate, consistent with admixture data (**Supplementary Figure 8**). Previous Ugandan WGS isolates (from 2006-2022) clustered with the current Ugandan isolates (**Supplementary Figure 9**).

**Figure 2.**
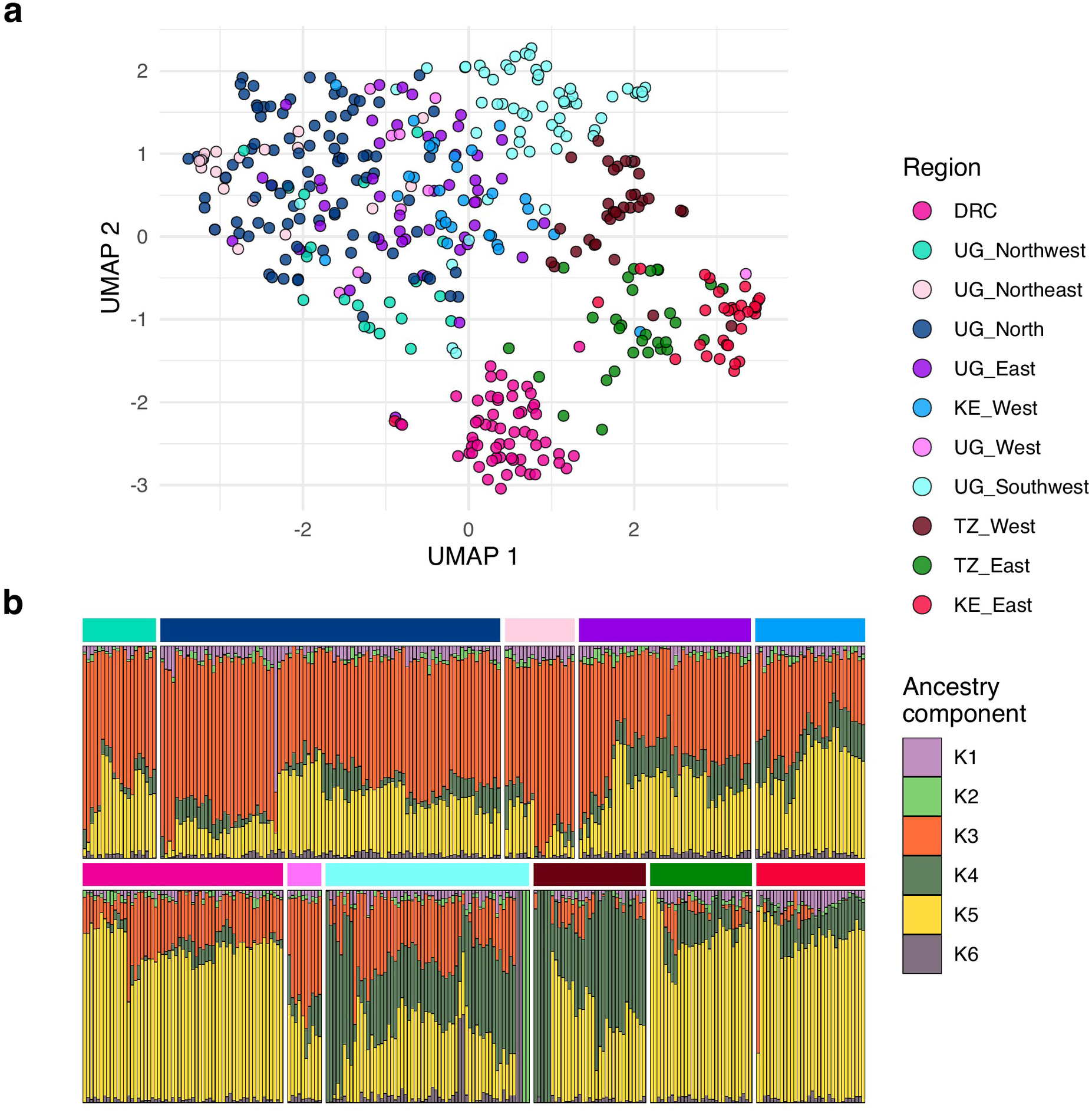
Population structure and admixture of Ugandan *P. falciparum* in regional context. **(a)** UMAP projections for isolates from Uganda and neighbouring countries (Democratic Republic of the Congo (DRC), Kenya, Tanzania), with isolates coloured by region in Uganda or by country; Ethiopian and Sudanese comparators are shown in **Supplementary Figure 8. (b)** Six population admixture (lowest cross-validation error population size) of Ugandan regions and neighboring countries. Horizontal bars on top of each set of admixture isolates correspond to regions and are colored the same as the UMAP projection. Legend title “Ancestry Component” has the admixture ancestry color key, which corresponds to the vertical bars in **b**.

The best estimate of the number of ancestral populations (*K)* for admixture (with the lowest cross-entropy validation score) was six. Admixture patterns were consistent with the UMAP results and showed that ancestry was shared across regions, although the proportion of diversity attributed to different ancestry components differed subtly (**Figure 2b**). Southwestern Ugandan parasites showed ancestry profiles more similar to Western Tanzania than to parasites from northern and eastern Uganda. In addition, these data suggest several distinct sub-populations were circulating in Southwestern Uganda. Two isolates from the Southwest (light green vertical bars, 100% K2 ancestry) were unrelated to any other population or isolate within the region, while two others (purple-grey vertical bars, 100% K6 ancestry) were similarly distinct but shared ~0.5 IBD with two additional isolates; these isolates appear in the admixture with approximately half their ancestry assigned to the K6 ancestry. Genetic differentiation between Ugandan regions, quantified using pairwise Jost’s D, was uniformly low but non-zero (**Supplementary Figure 10**).

### Geographic distribution of antimalarial resistance markers

Among the 249 monoclonal isolates analyzed, non-synonymous PfKelch13 mutations were present in 34.5%, with regional variation matching that reported previously by design **(Supplementary Figure 11)**^10^. C469Y and A675V predominated in the northern regions, whereas the Southwest carried a more diverse repertoire led by A675V, P441L, and R561H. At loci not used in matching (e.g., PfCRT, PfMDR1, PfDHFR, PfDHPS), the spectrum and geographic distribution also matched contemporaneous amplicon surveillance at these sites^10^; intermediate-frequency markers (e.g., PfMDR1 Y184F) were at lower prevalence in our isolates, as expected given selection for monoclonal infections. Chloroquine resistance mutations were uncommon (country-wide wild-type PfCRT CVMNK near-fixed at 95.2%), except in the Northwest region (K76T at 4.0%). For PfMDR1, wild-type N86 was near-universal, Y184F common (49.0%), and D1246Y infrequent (3.2%) **(Supplementary Figure 11)**. Antifolate resistance was widespread: the PfDHFR double mutant (N51I + S108N) was fixed (100%), co-occurred with near-universal PfDHFR C59R (96.0%), PfDHPS K540E (95.6%) and A437G (95.6%), with the full quintuple-mutant haplotype occurring at 92.4%. Mutations associated with higher-level resistance markers, PfDHPS A581G and PfDHFR I164L, were concentrated in the Southwest (46.7% and 60.0%, respectively) and absent from the North and Northeast.

We next examined loci implicated in antimalarial response that are not routinely included in targeted surveillance panels. For PfGCH1 (PF3D7_1224000), which encodes the first enzyme of the folate biosynthesis pathway and whose amplification is associated with antifolate resistance, only variant R230K was found at ~2% in the North, Northwest, and Southwest regions. For PfAAT1 (PF3D7_0629500), a digestive vacuole amino acid transporter implicated in chloroquine resistance, variants S258L, F313S, and F442L were detected. Variant S258L was found to be almost at fixation with a country-wide prevalence of 97%. Variant F313S was found in two Southwest isolates. Variant F442L was found at 1-5% prevalence in all regions and at 4% prevalence overall.

We performed a focused analysis of PfPX1 PIN haplotype (L1222P, M1701I, D1705N) and found it was present in every region, with an overall prevalence of 70%, ranging from 55% in the Northwest and Southwest to 90% in the Northeast. Substitutions at 16 other positions in PfPX1 were observed in at least two isolates each (**Supplementary Table 3**). Several substitutions were segregated by PIN background, occurring almost exclusively in PIN-containing isolates (D384A, D384Y, C1484Y) or in non-PIN isolates (D384G, D1711N, S1211R, C1484F), with eight additional variants at frequencies of 1-2%. Position 384 was the only multiallelic codon identified, carrying three distinct substitutions that differed in both PIN background and regional distribution. D384A was the most abundant substitution at this position and occurred only once without the full PIN haplotype, in an isolate that also encoded 1222P; it was detected only twice in the Southwest/West but reached 53.6% prevalence in the North/East, was highest in the North at 63.4%, and was present on 75% of all PIN haplotypes (**Supplementary Figure 12**). D384G was confined to the North and East, with 6 of 8 occurrences on non-PIN haplotypes. D384Y was found exclusively on PIN backgrounds. Among the non-PIN PfPX1 variants identified here, only C1484F has previously been associated with artemisinin resistance, in ART-R parasites from Southeast Asia^30^.

Two variants showed haplotype sharing across the Uganda–Tanzania border. S1211R was largely restricted to the Southwest and clustered closely with Western Tanzanian isolates carrying the same mutation (**Supplementary Figure 13**). Similarly, seven D384Y-PIN isolates, six from Western Tanzania and one from the Southwest, formed a long extended haplotype cluster, while a single additional D384Y–PIN isolate from the Northwest carried a distinct surrounding haplotype, consistent with independent occurrence (**Supplementary Figure 14**). D384Y has been detected only in Uganda and Tanzania and only from 2021 onward. By contrast, careful searching of Pf8 data, accounting for the multiallelic codon, found neither D384A nor D384G outside Uganda; the oldest available Uganda genome in which D384A has been seen was collected in 2017.

### Identity by descent sharing and signatures of selection at drug resistance loci

To identify genomic regions under recent positive selection in Uganda, we utilized both IBD and integrated haplotype Score (iHS). For IBD, using the output from hmmibd-rs, we calculated the proportion of isolate pairs sharing IBD at each SNP position and applied the normalised selection statistic (*X*_*iR,s*_)^14,27^, whereas for iHS we plotted the fraction of SNPs with |iHS| > 2 in 100-kb windows stepped every 10kb^31^.

IBD-based and iHS selection scans within the five Ugandan sub-regions (Northwest, North, Northeast, East, Southwest) identified significant selection at several antimalarial-resistance loci located in chromosomes 4-8, 12, and 13 (**Figure 3**). The strongest and most consistent signal was at *pfpx1* in both iHS and IBD, which was significant in all but one (Southwest) of the five regions; the peak was most significant in the North (-log10p ~ 179 in IBD, max |iHS| 3.78 at SNP Pf3D7_07_v3:893359 encoding PfPX1 AA 384) and was accompanied by extensive pairwise IBD sharing there and in the Northeast (up to ~55-74% of pairs), while remaining significant but weaker in the East and Southwest. For *pfkelch13*, only the North showed a significant selection peak in both iHS and IBD. Significant selection at *pfdhps* was detected in the North, Northeast, East, and Southwest but not in the Northwest. *pfgch1* showed significant peaks confined to the Northwest and Southwest. At *pfcrt*, the Northwest carried a significant selection peak, accompanied by the highest pairwise IBD sharing within any region (~20% of pairs). More localized significant signals were seen at *pfmdr1* in the Northeast only and *pfaat1* in the East only. No significant selection or appreciable IBD sharing was observed around *pfdhfr*.

**Figure 3.**
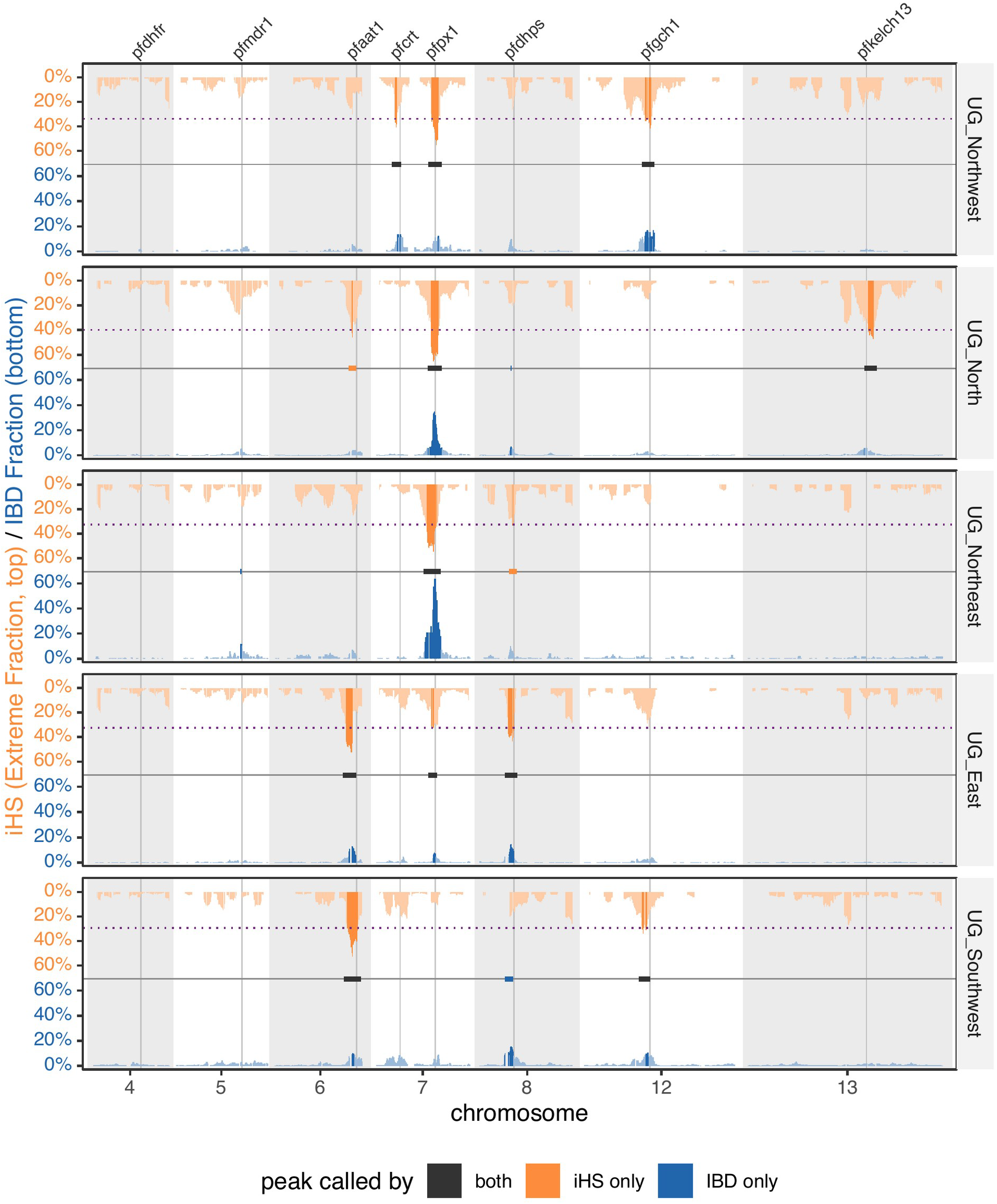
Genome-wide iHS and IBD-based selection/pairwise sharing scans IBD by region. Above the central horizontal axis (orange bands), the fraction of SNPs with |iHS| > 2 in 100-kb windows stepped every 10 kb. The purple dotted line indicates the threshold for the top 1% of iHS peaks and peaks above it are shown in dark orange, while peaks below are light orange. Below the central horizontal axis (blue bands), the proportion of isolate pairs sharing IBD at each SNP position within each region. SNPs with significant evidence of selection by the normalized IBD selection statistic (*X*_*iR,s*_, Bonferroni-corrected) are shaded dark blue while non-significant SNPs are shaded light blue. Horizontal bars on the central axis indicate significant peaks: black for both methods, orange for iHS only, dark blue for IBD only. Because iHS detects extended haplotypes without requiring that they are identical by descent, a locus selected on several independent backgrounds gives a strong iHS peak with little IBD sharing (*pfkelch13* in the North), whereas a sweep from a single origin gives both (*pfpx1* in the northern regions and the East). Known drug resistance genes with nearby or overlapping selection peaks are labeled and thin black vertical lines are drawn down the midpoint of the gene. Only chromosomes carrying significant peaks or genes of interest (4–8, 12, and 13) are shown here; **Supplementary Figures 15-16** show all chromosomes.

To further characterize the populations carrying *pfpx1* mutations showing evidence of selection and regional structure, we analyzed both *pfpx1* and *pfkelch13* genes with extended haplotype homozygosity (EHH), variant’s iHS, and gene specific IBD networks. The two loci showed strikingly different patterns of IBD sharing. At *pfpx1*, there were multiple significant |iHS| signals; the amino acid change with the highest country-wide |iHS| was D384A (3.92), followed by D1705N (3.35), L1222P (3.25), and M1701I (3.24). D384G did not show significant selection (**Figure 4**). Publicly available genomes from Ugandan isolates collected in 2020-2021 showed similar patterns of significantly high iHS and long EHH for all 4 variants (**Supplementary Figure 17**). The selection signal at the PIN variant sites appears to be carried by the D384A allele. Excluding D384A-PIN isolates substantially reduced EHH (**Supplementary Figure 18**) and largely removed the selection signals in iHS and IBD (**Supplementary Figure 19**), despite a PIN prevalence of 40-65% PIN in the population. The difference between the extended haplotypes can also be seen clearly when plotting out the genotypes 40kb up and downstream from *pfpx1* **(Supplementary Figure 14)**.

**Figure 4.**
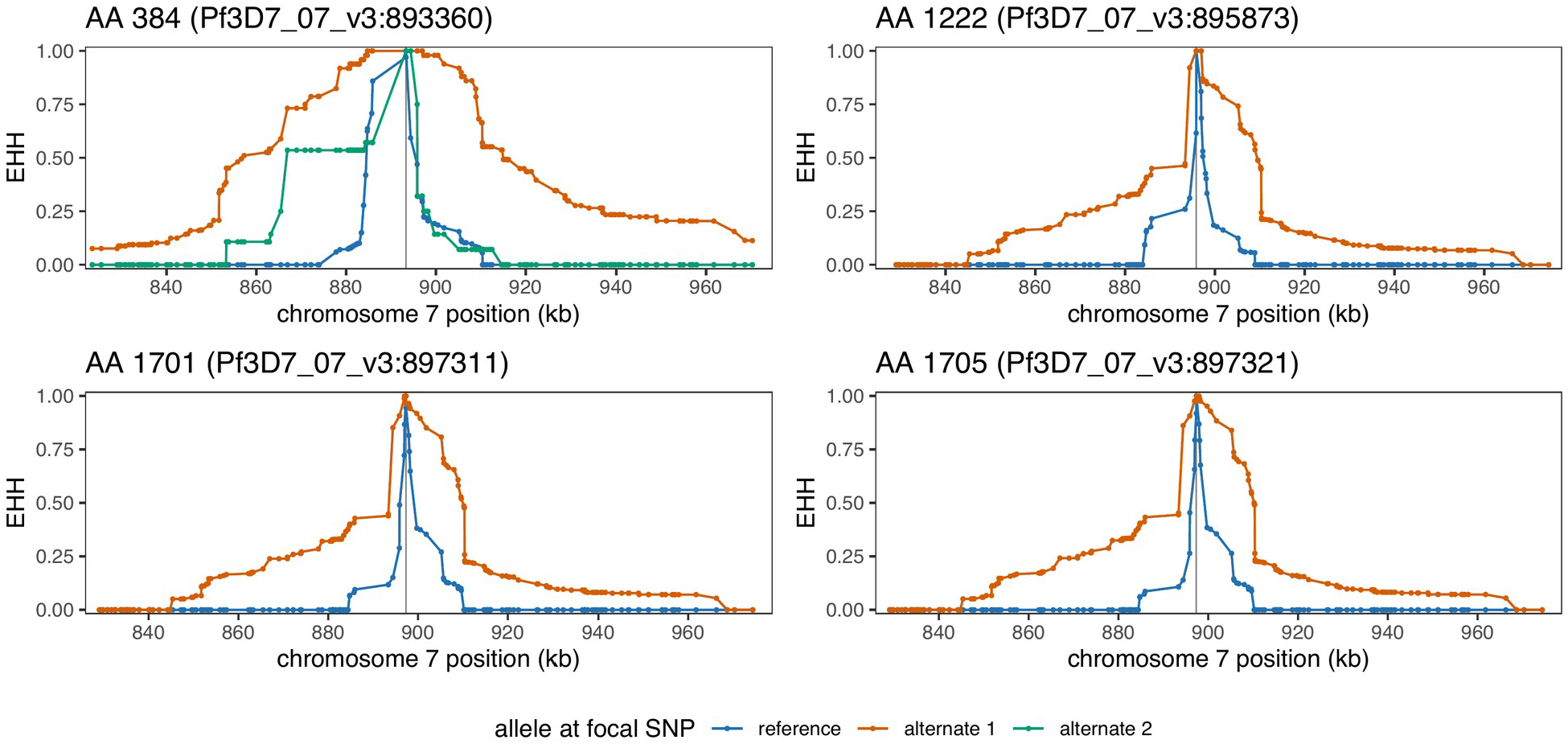
Extended haplotype homozygosity of PfPX1 amino acid positions 384, 1222, 1701, and 1705. EHH plots for Uganda 2023 isolates at positions 384 (top left), 1222 (top right), 1701 (bottom left), and 1705 (bottom right), were generated with the R package *rehh* via a custom wrapper that handles multiallelic sites by treating each alternative allele separately against the reference^32^. Blue lines denote the reference allele, and orange the first alternative allele. Positions 1222, 1701, and 1705 are biallelic; position 384 is multiallelic, and a second alternative allele is shown in green. At 384, the orange line corresponds to D384A and the green line to D384G, the non-PIN-associated allele. iHS scores for each alternative allele relative to the reference were: D384A, |iHS| 3.92 (p = 3.49 × 10^−4^); D1705N, |iHS| 3.35 (p = 3.19 × 10^−3^); L1222P, |iHS| 3.25 (p = 4.69 × 10^−3^); M1701I, |iHS| 3.24 (p = 4.77 × 10^−3^); D384G, |iHS| 0.23 (p = 1).

IBD sharing over the interval encoding PfPX1 amino acids 384–1705 was high both within and between northern Ugandan regions (**Figure 5c**). In IBD networks over this interval, where isolates are connected only when a shared IBD block completely covers the region, tight clusters are seen for the D384 mutations that co-occur with PIN (D384A-PIN and D384Y-PIN), while D384-PIN forms a much looser cluster (**Figure 5d**). Including the 2020-2021 Uganda data, trending the length of IBD blocks over D384A, the median block size has gone from 53.9kb in 2020, to 99.5kb in 2021, to 60.1kb in 2023, while the blocks around D384-PIN have remained a median size of ~25kb. This is consistent with D384A-PIN sweeping more recently than PIN alone. In addition, small clusters of isolates without the PIN variant were seen, primarily consisting of samples from Southwestern Uganda and Western Tanzania; among these is a cluster with isolates containing S1211R (a variant of unknown relevance), which was not observed with the PIN haplotype (**Supplementary Figure 13**).

**Figure 5:**
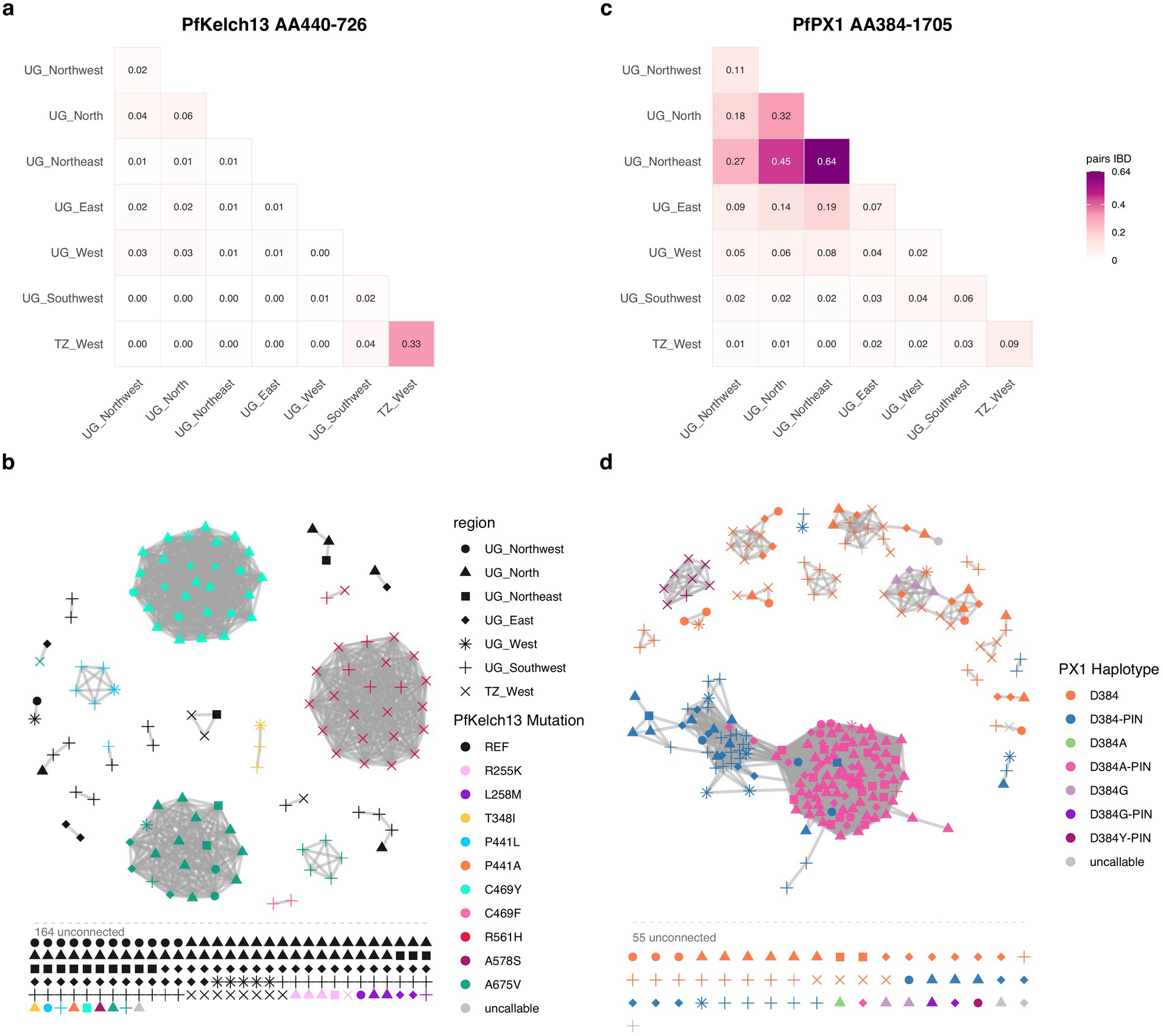
Between-region and between-sample IBD sharing at the PfKelch13 propeller domain and the PfPX1 384-1705 interval. **(a)** Proportion of isolate pairs whose IBD segments completely cover the genomic interval encoding PfKelch13 amino acids 440–726 (the propeller domain), within and between Ugandan regions and western Tanzania. **(b)** Network over the genomic interval encoding PfKelch13 amino acids 440–726, joining a pair of isolates when a shared IBD segment covers the interval in full. Node color indicates the PfKelch13 mutation carried by the isolate. **(c)** Proportion of isolate pairs whose IBD segments completely cover the genomic for the interval encoding PfPX1 amino acids 384-1705, which spans the 384–PIN haplotype. **(d)** Network over the the genomic interval encoding PfPX1 amino acids 384-1705 with node colour indicating the *pfpx1* haplotype, defined by the amino acid at position 384 together with the presence or absence of PIN. Node shape indicates region of origin in both networks. Isolates with no qualifying edge are shown in the grid beneath each network. Both heatmaps share a single color scale, so sharing is directly comparable between the two loci. Extended genotype patterns for (d) are shown in **Supplementary Figure 13**. Other neighboring populations were excluded because of differing sampling periods or an absence of informative IBD connections. Extended haplotype and genotype patterns are shown in **Supplementary Figures 14-26**.

In marked contrast, IBD sharing at *pfkelch13* was substantially lower across Ugandan regions and was partitioned regionally rather than forming a single connected block **(Figure 5a and 5b)**. This regional structure broadly reflected geographic differences in the mutational spectrum, though the distributions were not strictly partitioned. C469Y was concentrated in northern and northwestern Uganda, where pairwise IBD sharing formed a single cluster and extended haplotype analysis was consistent with a single origin (**Figure 5b, Supplementary Figure 20**). A675V was common in both the North and Southwest, forming distinct clusters and extended haplotypes in each, consistent with separate origins (**Figure 5b, Supplementary Figure 21, Supplementary Figure 22**). P441L and R561H were more prominent in the Southwest, and Ugandan R561H isolates clustered closely with Tanzanian isolates carrying the same mutation. At least two distinct R561H haplotypes were evident, consistent with the two independent R561H origins reported previously in Tanzania^33^ (**Supplementary Figures 21 and 23**). The regional peak at *pfkelch13* in the North (**Figure 3**) is therefore best explained by the local expansion of C469Y and A675V, and the iHS signal at this peak was considerably stronger than the IBD signal, which only marginally reached significance. This pattern would be expected when multiple haplotypes are under selection at the same locus, since iHS detects extended haplotypes without requiring that they be identical by descent.

We assessed the association between the PIN haplotype and PfKelch13 markers of ART-R countrywide using Fisher’s exact test. Within the northern and eastern regions (North/Northwest/Northeast/East), where the D384A-PIN haplotype is essentially confined (81/83 isolates), C469Y was enriched on the D384A-PIN background relative to all other callable PfPX1 haplotypes (28.4% vs 6.3%; OR 5.8, 95% CI 1.8–24.4; BH-adjusted p = 0.004) and showed the same trend relative to D384-PIN only (28.4% vs 7.7%; OR 4.7, 95% CI 1.0–44.3; BH-adjusted p = 0.067). A675V was unassociated with either 384-PIN combination. Although not statistically significant, 7 of the 8 P441L isolates identified countrywide carried the PIN haplotype and the small number of P441L observations limits the power of this comparison. (**Supplementary Figures 24-25**).

## Discussion

We report the first country-wide, whole-genome characterization of *P. falciparum* in Uganda, comprising 249 genomes from 26 sentinel sites across Uganda and, for regional context, publicly available genomes from seven neighboring populations. Genome-wide diversity was consistently high, and differentiation among regions was weak, but IBD analysis detected modest population structure. This was most pronounced in southwestern Uganda, where parasites were more closely related to one another than in any other region and shared ancestry and recent relatedness with western Tanzanian parasites, consistent with gene flow across the southern border. In addition, the PfPX1 and PfKelch13 loci associated with decreased ACT drug susceptibility differed in their patterns of haplotype sharing and selection. The PfPX1 PIN haplotype was predominant nationally by 2023, but the recent selection signal at this locus was carried by PIN haplotypes that had additionally acquired D384A, a substitution at a multiallelic codon concentrated in northern and eastern Uganda. PfKelch13 variants were regionally partitioned across multiple backgrounds, consistent with the independent emergence of different mutations. The two loci were not independent: C469Y was enriched on the D384A-PIN background in the regions where that haplotype occurs, raising the possibility that the *pfpx1* background shapes which *pfkelch13* mutations establish.

Nucleotide diversity and expected heterozygosity varied little among regions in Uganda, and genome-wide linkage disequilibrium declined rapidly with increasing physical distance, indicating a large, highly recombining parasite population in which frequent outcrossing between genetically distinct parasite clones breaks down haplotype structure^16^, consistent with results from earlier microsatellite and amplicon-sequencing results in Ugandan parasites^3,17–19^. UMAP on LD-pruned genotypes likewise collapsed most Ugandan parasites into a single cloud, and pairwise Jost’s D was consistently low, as expected where high within-population heterozygosity constrains allele-frequency based measures^34,35^. However, IBD, which tracks segments co-inherited from a recent common ancestor^15,36^, recovered geographically coherent patterns of relatedness that these metrics missed: a weakly connected network across Northern sites, and higher connectivity in the Southwest^14,16^.

Parasites from the Southwest region, which has lower transmission intensity than other regions of Uganda, were far more closely related than in any other region. The southwestern parasites shared little relatedness with parasites from elsewhere in Uganda, and their ancestry resembled a mixture of Ugandan, western Tanzanian, and two additional populations. Higher relatedness is expected where low transmission intensity reduces superinfection and outcrossing, favouring clonal propagation^3,15,37^. The region may also act as a low-transmission sink, receiving parasites imported from higher transmission settings, including those imported from the nearby countries of Rwanda, Tanzania and DRC^38–41^. A recent study of transmission-network inference at two very low transmission sites in southwestern Uganda estimated that >85% of infections were imported, with nearly all inferred source infections occurring in individuals reporting recent travel to other regions of Uganda^19^. However, domestic importation alone would not readily account for PfKelch13 R561H in our Southwest isolates, a variant first described in Rwanda and rare elsewhere in Uganda^9,11,42^; these Southwestern R561H isolates also share IBD over the PfKelch13 propeller domain with R561H isolates from Tanzania. Higher-level antifolate markers PfDHFR I164L and PfDHPS A581G were likewise concentrated in the Southwest (60.0% and 46.7%) in our analysis. This is consistent with prior surveillance data from Uganda which indicated that these markers are at highest prevalence in the southwest, with I164L more recently expanding into central and eastern Uganda^10,43^. Both mutations have also been reported in neighboring populations, including high prevalence of PfDHPS A581G in eastern DRC^44^ and PfDHFR I164L in eastern Rwanda^45^, with both seen in Western Tanzania^46,47^. Together with the ancestry shared between southwestern Ugandan and western Tanzanian parasites, this suggests gene flow across the southern and western borders, though confirming this hypothesis would require parasite genomes from Rwanda and eastern DRC.

Other cross-border comparisons also showed that Ugandan parasites were more similar to western than eastern Kenyan parasites. Western Kenyan isolates collected in 2015 and 2018 overlapped extensively with Ugandan parasites in the UMAP, particularly those from eastern Uganda, and shared similar ancestry components in the admixture analysis. This is consistent with previous genomic analyses of East Africa, in which parasites from western Kenya clustered with Ugandan parasites and shared a predominant ancestry component, while eastern Kenyan parasites were genetically more differentiated^12,48^. Contemporaneous DRC parasites from Kinshasa, far from Uganda, formed a distinct cluster, consistent with genomic evidence of isolation by distance within the DRC and across African *P. falciparum* populations^26,49^.

The northern vs southwestern differences seen in genome-wide relatedness were also evident at the loci linked to reduced ACT susceptibility. The PIN variant was predominant nationally in 2023 at 70% overall, ranging from 90% in the Northeast to 55% in the Northwest and Southwest. Niare et al. showed that PIN prevalence exceeded 50% in northern Uganda by 2016 and in eastern Uganda by 2023^21^, consistent with our finding of PIN in over half of isolates in every region. However, the extended PfPX1 haplotypes under selection in the northern regions were largely absent from the Southwest. Parasites with PX1 PIN haplotypes and wild type D384, which were most common in the southwest, had short extended haplotypes, reduced extended haplotype homozygosity compared to parasites with the D384A-PIN haplotype, and formed a loose cluster with a median shared block length of approximately 25kb. This is consistent with an earlier national expansion of D384-PIN, followed by recombination decay of its flanking haplotype^20,21^.

In contrast, in northern Uganda, the PfPX1 selection signal appeared to be driven by the accumulation of D384A on a PIN background. In the 2023 samples, this mutation carried the highest country-wide |iHS| of any PfPX1 amino acid change (3.92, p = 3.5 x 10-4); above D1705N (3.35), L1222P (3.25) and M1701I (3.24), indicative of strong recent selection, while D384G at the same codon showed none. D384A co-occurred with PIN in 97 of 98 isolates and was present on 75% of PIN haplotypes. Its distribution matched the geography of the selection signals seen in Figure 3: 63.4% in the North and 53.6% across the northern and eastern regions, versus two (of 70) isolates in the Southwest and West, where the PfPX1 signal was weakest. Isolates carrying D384A-PIN formed tight IBD clusters over the 384 to 1705 interval, with median shared blocks of 54 to 100 kb across sampling years compared with approximately 25 kb for D384-PIN. D384A was detected as early as 2017 but not elsewhere in Pf8 isolates once the multiallelic codon was accounted for^50^. These observations are consistent with D384A arising on an already widespread PIN background and expanding recently. Its near-absence from the Southwest and West may reflect its recent emergence combined with conditions there: lower transmission, greater parasite exchange with western Tanzania than with northern Uganda, and the presence of other PfPX1 variants on the PIN background. No phenotype has been reported for any substitution at position 384, and the reduced dihydroartemisinin and lumefantrine susceptibility described for PIN was measured on backgrounds that were not resolved at this codon^20,21^. Ex vivo drug susceptibility testing of isolates differing only at position 384 will thus be of great interest.

Southwestern and western Tanzanian D384Y isolates carried a different substitution at the same codon but at different genomic locations, with D384A occurring at second codon base (Pf3D7_07_v3:893359) and D384Y at first codon base (Pf3D7_07_v3:893358). D384Y was found only on PIN backgrounds, only in Uganda and Tanzania, and only from 2021 onward. Seven D384Y-PIN isolates, six from western Tanzania and one from the Southwest, shared a long extended haplotype, whereas a single northwestern D384Y-PIN isolate carried a distinct surrounding haplotype consistent with independent occurrence. Two different substitutions at position 384 therefore arose on PIN backgrounds, each geographically restricted, each first detected within the past decade, and each carried on a tightly clustered haplotype, while a third substitution at the same codon (D384G), occurring mostly off PIN, showed no evidence of selection. One explanation for these patterns is convergent selection at position 384 on a PIN background, although we did not run selection scans in the Tanzanian comparator isolates, and the D384Y cluster comprises seven genomes from a population sampled in 2022 in which PIN was less common. Considering the amino acid substitutions, alanine and tyrosine, a methyl group and a bulky aromatic ring, share little chemical similarity beyond the loss of the aspartate, so selection at this codon may act principally on loss of the carboxylate. Notably, position 384 lies outside the PX domain predicted to bind phosphoinositide, in a region of PfPX1 with no assigned function. Other PfPX1 variation persisted in the south independent of D384A, including a southwestern PIN cluster defined by C1484Y and a cross-border non-PIN cluster defined by S1211R, a variant uncommon among African isolates that reached prevalences of 33% in Indonesia and 12% in Papua New Guinea^50^.

No PfKelch13 mutation approached the frequency or the haplotype dominance of D384A-PIN, and the only significant selection signal at PfKelch13 was in the North. Extended haplotype analysis resolved a single origin for C469Y but separate northern and southwestern origins for A675V. Within the northern and eastern regions, where D384A-PIN was essentially confined (81 of 83 isolates), C469Y was enriched on the D384A-PIN background relative to all other PIN haplotypes, and in the same direction relative to D384-PIN. A675V was not associated with either 384-PIN haplotype, and 7 of the 8 P441L isolates identified countrywide carried D384-PIN and P441L was mostly confined to the southwest. Any advantage of the D384 mutations therefore may be specific to particular PfKelch13 mutations. In our data, an advantage is established only for C469Y, which longitudinal surveillance indicates has been increasing in northern Uganda relative to A675V^10^. PIN was already common in northern Uganda before these PfKelch13 mutations expanded^8,21^, so C469Y is more likely to have arisen on a D384A-PIN background than the reverse. D384A-PIN may be permissive for C469Y rather than causal for ART-R, adding to the resistance phenotype or offsetting a fitness cost of the mutation. Artemisinin resistance in Southeast Asia also arose preferentially on a defined genetic background^30^, and the reverse ordering has been described there for partner-drug resistance, where a piperaquine-resistance determinant was acquired on an existing *pfkelch13* background^23^.

Loci associated with decreased susceptibility to ACT partner drugs and resistance to chemoprevention agents showed regional structure, despite uniform antimalarial and chemopreventative policy throughout Uganda. Regional differences at these markers may reflect where resistance arose or was imported, and how strongly local conditions favored their selection. At the antifolate loci, the PfDHFR/PfDHPS quintuple mutant is close to fixed, with country wide prevalence 92.4%; the *pfdhps* selection signal therefore likely reflects historical selection on now largely fixed mutations, with more recent activity layered onto that background. The higher-level resistance markers PfDHPS A581G and PfDHFR I164L were concentrated in the Southwest, a distribution compatible with more efficient selection where transmission, host immunity, and within-host competition are lower, and with possible importation across the southern and western borders as discussed above. Notably, comparable high-level antifolate resistance markers have been reported in neighbouring Rwanda and DRC, and in newly arrived refugees settling in western Uganda from the DRC^10,26,40,41,51–53^. A *pfgch1* signal confined to the Northwest and Southwest may likewise reflect linked antifolate resistance, since resistance-associated variation at this locus has been reported to be driven by copy-number amplification that can co-select with *pfdhfr* and *pfdhps* alleles^54,55^.

At *pfcrt*, chloroquine resistance markers were generally uncommon, consistent with long-standing AL use, which selects for wild-type PfCRT K76 and PfMDR1 N86. The detection of K76T in the Northwest, accompanied by a significant *pfcrt* selection signal and the highest pairwise IBD sharing at this locus of any region, was therefore notable and may reflect parasite movement from South Sudan, where artesunate–amodiaquine, which selects for K76T and PfMDR1 N86Y, is the first line antimalarial^56^. Consistent with this interpretation, both mutations were substantially more frequent among recently arrived South Sudanese refugees than among patients attending nearby northern Uganda health facilities^41^. Considering other potential drug resistance markers, the localised significant signals at *pfmdr1* (Northeast) and *pfaat1* (East), together with the near-fixed PfAAT1 S258L and the previously undescribed F442L variant, are of unknown significance^10,14,22^.

Overall, these data establish a 2023 country-wide genomic baseline for *P. falciparum* in Uganda. The strongest selection signal in the genome lay at a locus in PfPX1 recoverable because local haplotype assembly resolved a multiallelic codon that routine biallelic SNP filtering discards. Two loci contributing to reduced ACT susceptibility are diverging in their dynamics: D384A-PIN is expanding as a single background in northern Uganda, while PfKelch13 mutations are emerging repeatedly across the country. Establishing whether the D384-PIN background alters drug susceptibility, and whether it shapes which PfKelch13 mutations become established, will require ex vivo phenotyping and repeated sampling coordinated across national borders.

Several features of the study limit our interpretations. The Uganda data represent a single cross-section from 2023, so any inference about the age or direction of selection is necessarily indirect. Publicly available WGS data from Uganda’s neighbours were also incomplete and uneven in both year and location, limiting interpretation of cross-border sharing at *pfpx1* and *pfkelch13*. For example, South Sudanese and Ethiopian genomes were too scarce to include in the analysis, and no recent Rwandan genomes were publicly available, leaving unrepresented the borders most relevant to the *pfcrt* signal in the Northwest and the antifolate markers in the Southwest. Where comparator data were available, geographic and temporal mismatches limited their informativeness: the DRC genomes were from Kinshasa rather than the eastern provinces bordering Uganda, and the western Kenyan genomes predated the regional expansion of both the PfPX1 PIN and PfKelch13 haplotypes. Furthermore, the primary analysis sample set was restricted to monoclonal infections and assembled by amplicon-matched downsampling to correct for the initial oversampling of PfKelch13 mutants, rather than drawn as a random sample. However, at loci not used in matching, both the drug resistance mutation spectrum and its geographic distribution agreed with contemporaneous amplicon surveillance at the same sites^10^, indicating that the final cohort recovered the resistance landscape of the underlying population. Sampling was also uneven across regions; the West (n = 10) was covered too sparsely to enter the inferential comparisons; denser sampling would improve power to resolve fine structure and to detect weaker selection. Despite these limitations, our data represent the most geographically comprehensive whole-genome dataset assembled for Ugandan *P. falciparum* to date.

## Methods

### Study design, sample selection, and quality control

This study was conducted as part of the Implementing Malaria Molecular Surveillance in Uganda (IMMRSE-U) project, a nationwide genomic surveillance platform designed to generate high-resolution genomic evidence for parasite population structure analysis, transmission connectivity mapping, and antimalarial drug resistance monitoring. The analysis used *Plasmodium falciparum*-positive dried blood spot (DBS) samples collected from microscopy positive patients in 2023 from sentinel health facilities across Uganda. The initial sequencing dataset comprised 340 samples. Four were removed because of low WGS coverage, and a further four were excluded because of low within-host fixation index (*F*_ws_ < 0.95), consistent with mixed or complex infections that could bias haplotype-based and relatedness analyses, reducing the dataset to 332 monoclonal genomes. Samples were further downsampled to 8–10 samples per sentinel site to achieve geographic representation and reduce potential bias from overrepresentation of *pfkelch13* mutations from the Multiplex Amplicons for Drug, Diagnostic, Diversity, and Differentiation Haplotypes using Targeted Resequencing (MAD4HatTeR)^57^ amplicon sequencing data from the same sites in the same year^10^ to correct for bias in over sequencing of samples with *pfkelch13* mutations. This yielded the final analysis cohort of 249 high-confidence genomes from 26 sentinel sites distributed across six geographic regions: North (10 sites, n = 93: Aboke, Aduku, Alebtong, Amolatar, Atiak, Lalogi, Namokora, Orum, Padibe, Patongo), East (5 sites, n = 47: Busitema, Kigandalo, Kiyunga, Nagongera, Nawaikoke), Southwest (6 sites, n = 60: Bikurungu, Kamwezi, Karambi, Kasambya, Kihihi, Muko), Northwest (2 sites, n = 20: Metu, Opia), Northeast (2 sites, n = 19: Lokolia, Nadunget), and West (1 site, n = 10: Kigorobya) (**Supplementary Figure 1**). Sentinel sites were selected to provide broad spatial coverage of Uganda across different epidemiological and ecological settings. Isolates were grouped by collection site and region; regional groupings were used for primary comparative analyses, while site-level groupings were used for fine-scale analyses of parasite relatedness and transmission connectivity. West is retained only for descriptive counts and excluded from inferential comparisons due to n = 10.

### Ethical considerations

The study obtained ethical approval from the University of California, San Francisco Institutional Review Board (Reference 341609), the Makerere University School of Biomedical Sciences Research Ethics Committee (Protocols SBS-2021-167 and SBS-2024-566), and the Uganda National Council for Science and Technology (Protocols HS2309ES and HS5584ES). Written informed consent was obtained from all adult participants. Parental or guardian consent was obtained for all participants under 18 with additional assent from children aged 8–17 years.

### DNA extraction, selective whole-genome amplification, and sequencing

Genomic DNA was extracted from DBS using Tween-20/Chelex 100 as previously described^58,59^. After DNA extraction, *P. falciparum* parasite density was quantified by varATS-based quantitative PCR as previously described^60^. Because DBS frequently contains substantially more human DNA than parasite DNA, selective whole-genome amplification (sWGA) was used to enrich *Plasmodium falciparum* DNA before sequencing. sWGA was performed according to the previously described protocol^61^, with two modifications: phi29-XT DNA polymerase was used in place of the original phi29 polymerase, and the stock concentration of the 6A and 10A *P. falciparum*-specific primers was increased from 100 µM to 500 µM. All other steps were performed as previously described. Successful amplification was assessed by fluorometric DNA quantification, and negative extraction and amplification controls were included to monitor potential contamination.

Sequencing libraries were prepared from sWGA-amplified DNA using the Illumina DNA Prep workflow according to the manufacturer’s instructions. The workflow was modified by quantifying individual libraries before pooling; libraries were normalized to the same concentration and pooled to promote even representation across samples. The pooled library was subsequently size-selected on a 2% agarose gel, and fragments of 500–800 bp were excised and purified using the Zymoclean Gel DNA Recovery Kit according to the manufacturer’s instructions before final quantification and fragment-size assessment. Libraries were sequenced using paired-end reads on an Illumina NovaSeqX platform, targeting approximately 40–50 million paired-end reads per sample. The complete modified sWGA and whole-genome sequencing library preparation protocol is available at protocols.io: https://www.protocols.io/view/plasmodium-falciparum-whole-genome-sequencing-swga-q26g7qmj9lwz/v1.

### Read processing, alignment, variant calling, curation, and SNP quality control

Raw FASTQ files were assessed for read quality, adapter contamination, read length distribution, and per-base quality using FastQC v0.12.1^62^. Adapter sequences and low-quality bases were removed using Trimmomatic v0.40^63^, and reads failing length thresholds after trimming were discarded. Trimmed reads were aligned to the *P. falciparum* 3D7 reference genome (version 3) using BWA-MEM v0.7.19^64^. Alignment files were sorted and indexed using SAMtools, and duplicate reads were marked and removed using Picard MarkDuplicates v3.4.0^65^. Per-sample sequencing and alignment metrics including mean genome-wide coverage, breadth of coverage across the core genome, mapping rate, duplicate proportion, and genotype missingness were calculated using mosdepth v0.3.14^66^.

Variant discovery was performed using an optimised multi-sample GATK4 workflow^67^. Briefly, variants were initially called separately for each sample using GATK HaplotypeCaller v4.6.1.0^68^ in genomic variant call format (GVCF) mode. Sample-level GVCFs were consolidated using GenomicsDBImport and jointly genotyped with GenotypeGVCFs to generate a multi-sample variant call format (VCF) file. A sequential filtering procedure was applied to obtain a high-confidence set of SNPs. At the locus level, variants were excluded when they failed any of the following hard-filtering criteria: quality by depth (QD) <5, mapping quality (MQ) <55, strand odds ratio (SOR) >3, mapping-quality rank-sum test (MQRankSum) <−5, and read-position rank-sum test (ReadPosRankSum) <−5. SNPs located within tandem repeats were removed using BEDTools v2.31.1^69^, and analyses were restricted to the *Plasmodium falciparum* core genome. Hypervariable, subtelomeric, centromeric, repetitive, and poorly mappable genomic regions were excluded to minimise mapping and variant-calling artefacts. Highly polymorphic multigene families, including var, rifin, and stevor, were also excluded.

Genotype-level filtering required a minimum read depth of two reads and a minimum within-sample allele frequency of 0.01. Sample completeness was subsequently evaluated using allele-depth support, and samples were excluded when fewer than 85% of evaluated loci had adequate coverage, defined as an allele-depth sum of at least 10 reads. At the locus level, SNPs were retained only when genotype missingness was below 5% and at least 95% of samples had adequate sequence coverage. The final analysis dataset was further filtered using BCFtools v1.23.1^70^ to retain variants with a minor allele frequency greater than 0.02. Within-host infection complexity was assessed by estimating the within-host fixation index (*F*_*ws*_) using the moimix R package v0.0.2.9001^71^. Samples with *F*_*ws*_ >0.95 were classified as predominantly monoclonal and retained for population-genomic analyses. Following variant-, genotype-, locus-, and sample-level quality control, the final analysis-ready dataset comprised 28,926 high-confidence biallelic SNPs across 249 samples, with high overall genotyping completeness.

### Genetic diversity analyses, linkage disequilibrium, and antimalarial drug-resistance genotyping

Genetic diversity was estimated across the full dataset and stratified by site and region. Diversity metrics were calculated using scikit-allel v1.3.13^72^ and VCFtools v0.1.16^73^. Nucleotide diversity (*π*) was calculated as the average number of pairwise nucleotide differences per site, computed genome-wide and in 10-kb sliding windows with 5-kb steps, retaining windows with sufficient callable sites. Expected heterozygosity (*H*_e_) was calculated from allele frequencies as *H*_e_ = 1 − Σ *p*_i_^2^, where *p*_i_ is the frequency of allele *i* at a locus. Haplotype diversity was estimated to quantify the number and frequency distribution of distinct haplotypes within regions. Linkage disequilibrium was calculated as the squared allele-frequency correlation coefficient (*r*^2^) between SNPs using PLINK v1.9^74^, with a maximum distance of 50 kb, followed by binning into 1-kb distance windows. LD decay curves were generated for each region and genome-wide.

Key drug resistance loci including *pfkelch13, pfcrt, pfmdr1, pfdhfr, pfdhps*, and *pfpx1* were genotyped from the high-confidence variant call set and amino acid substitutions were called by local haplotype assembly using PathWeaver^75^, which enabled detection of multiallelic codons that routine biallelic SNP filtering would have removed. Specific mutations assessed included *pfkelch13* propeller-domain mutations (C469Y, A675V, P441L, R561H, R255K, T348I, L258M, C469F, A578S, P441A plus any novel mutations), *pfcrt* codons 72–76, 93, and 97, *pfmdr1* mutations N86Y, Y184F, S1034C, N1042D, and D1246Y, *pfdhfr* mutations N51I, C59R, S108N, and I164L, *pfdhps* mutations A437G, K540E, A581G, and A613S/T, *pfpx1* mutations L1222P, C1484Y, M1701I, and D1705N and any novel mutations, *pfgch1* mutation R230K, and *pfaat1* mutations S258L, F313S, and F442L. Resistance allele frequencies and haplotype distributions were summarised across regions and overlaid onto spatial and IBD network visualisations to assess the geographic distribution and lineage context of specific resistance profiles.

### Population structure analyses

To contextualize the geographic structure observed in Ugandan samples, publicly available *P. falciparum* WGS data from surrounding countries were incorporated (**Supplementary Table 2**). Public datasets were selected to provide regional context while prioritising recent collections, monoclonal infections, high-coverage samples, and sample sizes comparable to the regional subsets in the Ugandan dataset. The comparative dataset included samples from the DRC (n = 59, 2022), Eastern Kenya (n = 32, 2019–2020), Western Kenya (n = 20, 2015, n =10 2018), Western Tanzania (n = 33, 2022), Eastern Tanzania (n = 30, 2019), Ethiopia (n = 16, 2013–2015), Sudan (n = 6, 2015– 2017), and previous Uganda samples (n = 74, 2006-2022). Comparative WGS data were harmonised with the Ugandan dataset by restricting all samples to shared high-confidence biallelic SNPs, applying consistent missingness and quality filters, and excluding low-coverage or highly polyclonal samples.

Uniform Manifold Approximation and Projection (UMAP)^28,29^ was performed using 30 principal components derived from an LD-pruned genotype matrix, allowing major axes of genetic variation to be summarised before projection into two-dimensional space using the R package uwot^29^ and parameters of 15 minimum neighbors, min distance 0.1 and a metric of euclidean. UMAP analyses were performed separately for the Uganda-only dataset and for the combined Uganda plus neighbouring-country dataset; samples were coloured by region and country. Ancestry and admixture patterns were inferred using the Landscape and Ecological Association studies (LEA) R package^76^, employing the sparse non-negative matrix factorisation (sNMF) framework. The LD-pruned SNP dataset was converted into the genotype format required by LEA. The number of ancestral populations (*K*) was evaluated across a broad range of values (1-35), with cross-validation (cross-entropy) error used to identify the most informative *K*; each *K* was run across replicate analyses to evaluate stability. Individual ancestry coefficients were visualised as stacked bar plots ordered by geographic region. Genetic differentiation between regions was quantified using pairwise Jost’s D^34,35^ across 10-kb windows with 5-kb steps, providing a complementary measure of allelic differentiation that is particularly useful in highly diverse populations where *F*_ST_ can be constrained by high within-population diversity. Genome-wide mean values were summarised as symmetric matrices and visualised as heatmaps.

### Identity-by-descent analysis, relatedness networks, and genome-wide selection scan

Pairwise IBD was estimated using hmmibd-rs v0.1.5^27^, a parallelised implementation of the hmmIBD algorithm^77^, which extends the first-order hidden Markov model (HMM) framework for IBD detection between haploid genomes. IBD segments containing fewer than 15 SNPs or shorter than 15 kb were excluded, as short segments are likely to represent distant population-level sharing or false positives arising from residual linkage disequilibrium rather than recent common ancestry^14^

For each pair of isolates, the genome-wide pairwise IBD fraction was calculated as the total length of inferred IBD segments divided by the total genome length assayed. IBD-positive pairs were defined as those with a genome-wide pairwise IBD fraction ≥ 1/32 (5 generations to the most recent common ancestor), a threshold that identifies pairs sharing substantially more of their genome IBD than expected by chance and that has been widely used in *P. falciparum* IBD analyses^14,22^. To visualise patterns of parasite relatedness, we constructed sample-level and site-level IBD networks using the igraph R package v2.3.2^78^. Genome-wide IBD networks between isolates (Figure 1a) were constructed by drawing an edge between each pair of isolates sharing more than 1/32 (0.03125) of the accessible genome IBD, with edge width scaled to the shared fraction. Isolates with no pair above this cutoff are shown unconnected. In the site-level network, nodes represent sentinel sites and edges are weighted by the mean pairwise IBD fraction between sites. Networks between isolates were created at the genome wide level by fraction of the chromosome in IBD between samples and at the locus/gene level by connecting samples if they had an IBD block that completely covered the locus or gene of interest. Haplotype genotype plots with samples as rows and columns as variants centered around specific genes of interest were created using custom R code now packaged in an R package https://github.com/nickjhathaway/plasgenomicsutilsR. Extended haplotype homozygosity graphs were also created using this package by wrapping R package rehh^32^. In short, these graphs are created by picking a specific SNP of interest,grouping isolates into reference or alternative and going out in both directions for this SNP plotting the fractions of isolates in each group continued to sharing SNPs at each adjacent SNPs to the original. The iHS for each SNP was also calculated by wrapping rehh and in short is the natural log of the area under the curve of the alternative allele divided by the reference allele curve area. This package was wrapped to handle multiallelic SNPs, iHS scores were always calculated by alternative to reference only, so at sites with multiple alternatives the IHS score is relative to the reference allele and not to the other alternative.

A genome-wide selection scan was performed to identify genomic regions with excess pairwise IBD sharing, indicative of recent positive selection, using the normalized composite statistic as previously defined^14^. Briefly, a binary IBD-status matrix was constructed with rows corresponding to SNPs and columns corresponding to isolate pairs, where each entry indicates whether a pair shares IBD at that SNP. For each column (pair), the column mean was subtracted from all rows to account for the overall relatedness between the pair, ensuring that highly related pairs do not inflate signals at neutral loci. Next, for each row (SNP), the row mean was subtracted and the result divided by the square root of *p*_i_ (1 − *p*_i_), where *p*_i_ is the population allele frequency of SNP *i*, adjusting for differences in allele frequency that affect IBD detection power. Row sums were calculated and divided by the square root of the number of pairs to produce per-SNP summary statistics. These statistics were then normalised genome-wide by binning SNPs into 100 equally sized allele-frequency bins and subtracting the bin mean and dividing by the bin standard deviation, yielding z-scores. The z-scores were squared to produce chi-squared-distributed test statistics with 1 degree of freedom, from which p-values were derived. A −log _10_ transformation was applied, and significance was assessed after Bonferroni correction for multiple testing across all SNPs. Region-specific scans were performed by restricting the analysis to isolate pairs within each geographic region. The amount of pairwise sharing within a region was also calculated for each SNP.

## Code availability

Packages/code used to analyse and create figures for the manuscript can be found at https://github.com/nickjhathaway/plasgenomicsutils (version: v0.4.0) and https://github.com/nickjhathaway/plasgenomicsutilsR (version: v0.5.0). Analyses done for this manuscript can be found at https://doi.org/10.5281/zenodo.22926056.

## Data availability

Whole-genome sequencing data generated in this study have been deposited at PRJNA1429107. Publicly available WGS data from the DRC, Kenya, Ethiopia, Sudan, and Tanzania were obtained from NCBI SRA (**Supplementary Table 2**).

## Acknowledgements

We would like to acknowledge all the IMMRSE-U participants for their involvement in the study and all study staff who helped to complete the study. We would like to acknowledge the Greenhouse laboratory at UCSF and the Chan Zuckerberg Biohub for performing WGS.

## Funding support

This work was supported by grants from the Bill & Melinda Gates Foundation (INV-035751, INV-037316, INV-081860, INV-069975). SK was funded by the Makerere University Data Science Research Training Program to Strengthen Evidence Based Health Innovation, Intervention and Policy (MakDARTA) under the Fogarty International Center of the National Institutes of Health (NIH) under Award Number U2RTW012116 as a PhD fellow. GD was supported by PRISM NIH grant U19AI089674. JB was supported by NIH-NIAID K23AI166009. BG was supported by NIH-NIAID K24AI144048. MDC was supported by NIH grant R01 AI173557.

## Disclaimer

The funders had no role in study design, data collection and analysis, decision to publish, or preparation of the manuscript.

## Author contributions

SK, JB, BG, and IS conceived and designed the study. FVA, MD, and JB performed the laboratory experiments. SK and NH conducted the bioinformatic and statistical analyses. SK, NH, JB, BG, IS, and MM contributed to the interpretation of the results. IS, ST, JM, SLN, CB, GM, JNN, JB, BG, DPK, and MRK supervised the study. SK, NH, and JB drafted the manuscript. MM, TK, VA, FS, BN, ST, IW, JN, JM, EW, CMS, DPK, JNN, DJ, GM, CB, SLN, MRK, BBA, IS, AAD, FVA, GD, PR, MDC, BG, JB, and NH provided critical intellectual feedback and contributed to refining the analyses and manuscript. All authors reviewed and approved the final version of the manuscript.

## Competing interests

The authors declare no competing interests.

## Notes

### Competing Interest Statement

The authors have declared no competing interest.

