## Supplemental data for "Whole-genome characterization of *Plasmodium falciparum* population structure and drug resistance across Uganda and neighboring countries"

[+]Co-last authors

Nicholas Hathaway

Jessica Briggs

**Keywords:** *Plasmodium falciparum*; Uganda; whole-genome sequencing; population structure; genetic diversity; identity-by-descent; Selective sweeps; Artemisinin resistance

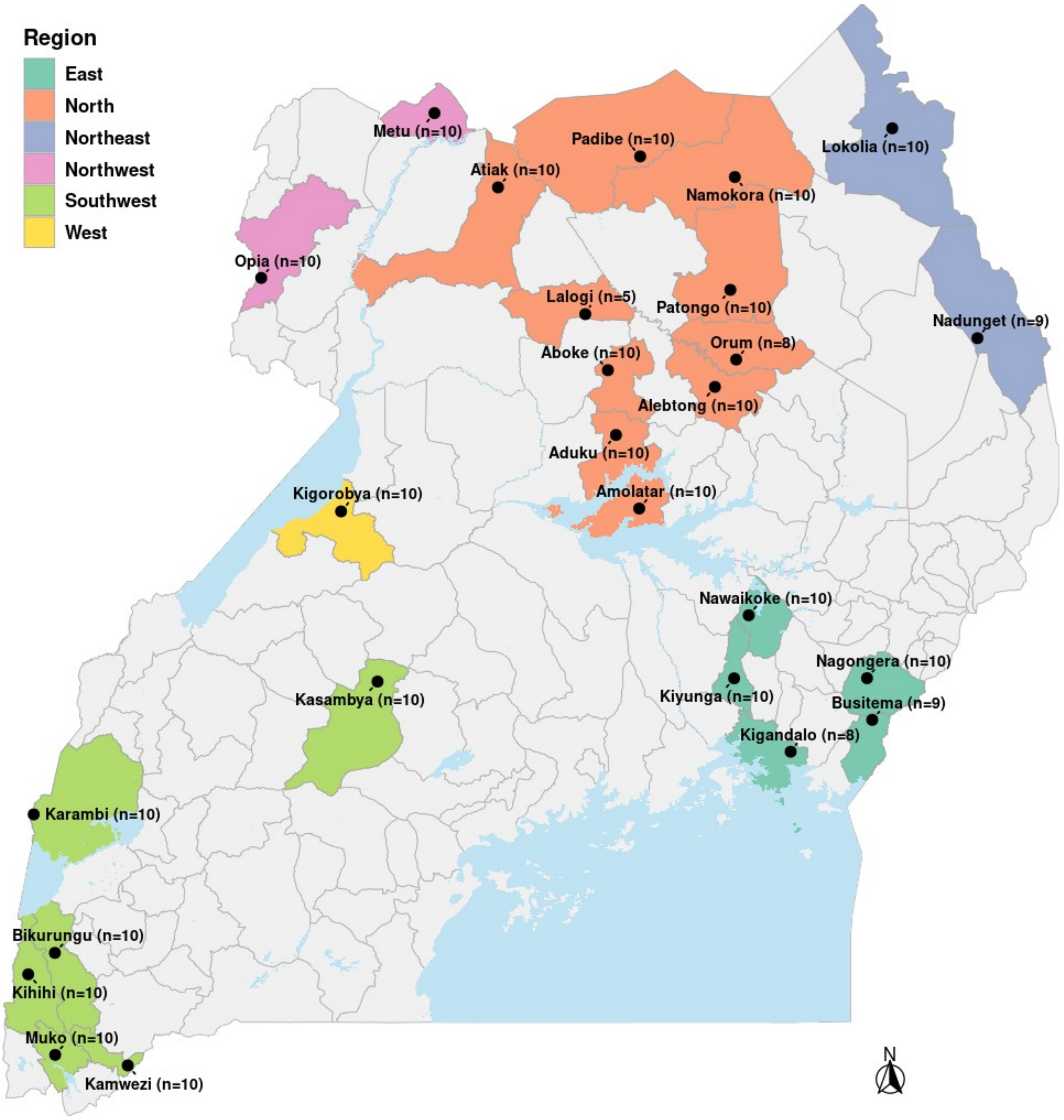


**Supplementary Figure 1. Map of sentinel sites by region with sample counts.** Geographic distribution of the 26 sentinel surveillance sites across Uganda's six epidemiological regions (Southwest, East, Northeast, North, Northwest, and West). Regions are represented by colors. Annotation in brackets at the end of each site name indicates the number of *P. falciparum* whole-genome sequences successfully generated from each site.

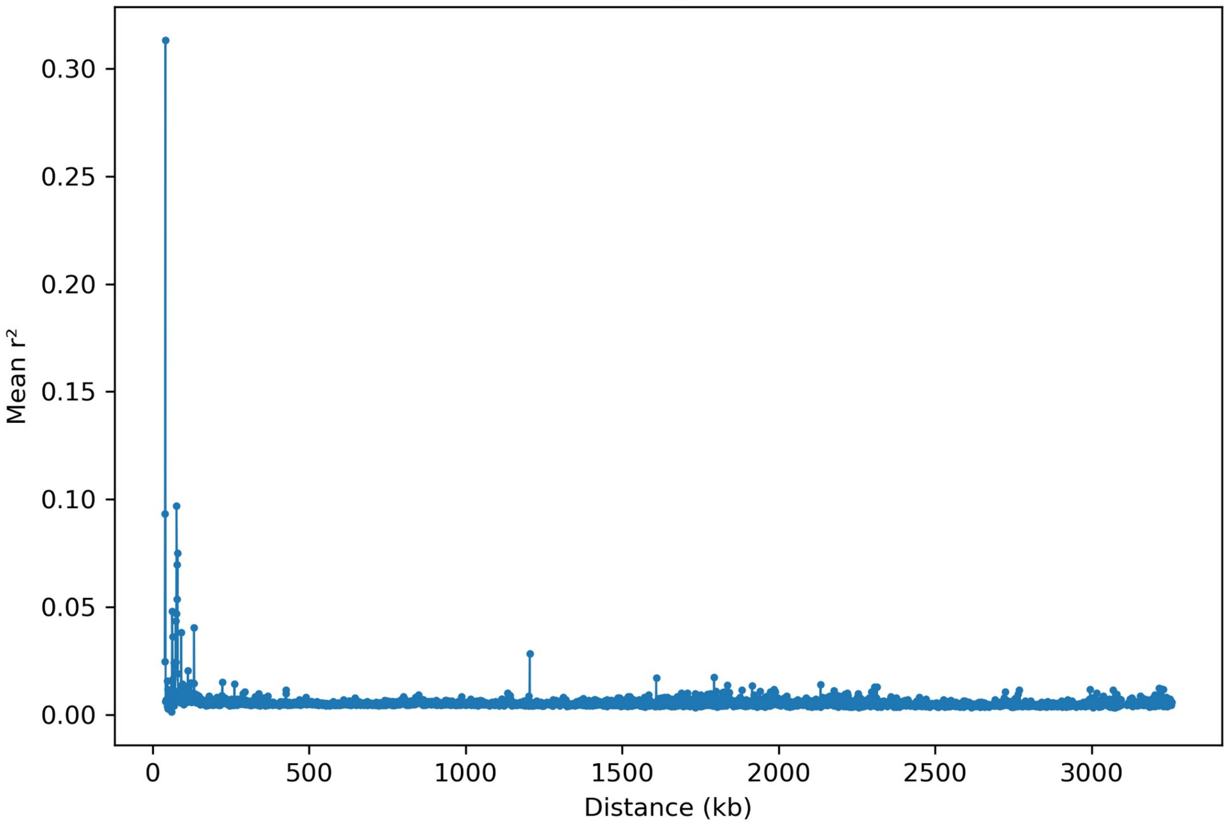


**Supplementary Figure 2. Genome-wide LD decay curve (all samples).** The curve illustrates the rate at which statistical association between alleles decays with physical distance, providing a baseline measure of recombination and population structure across the full Ugandan dataset.

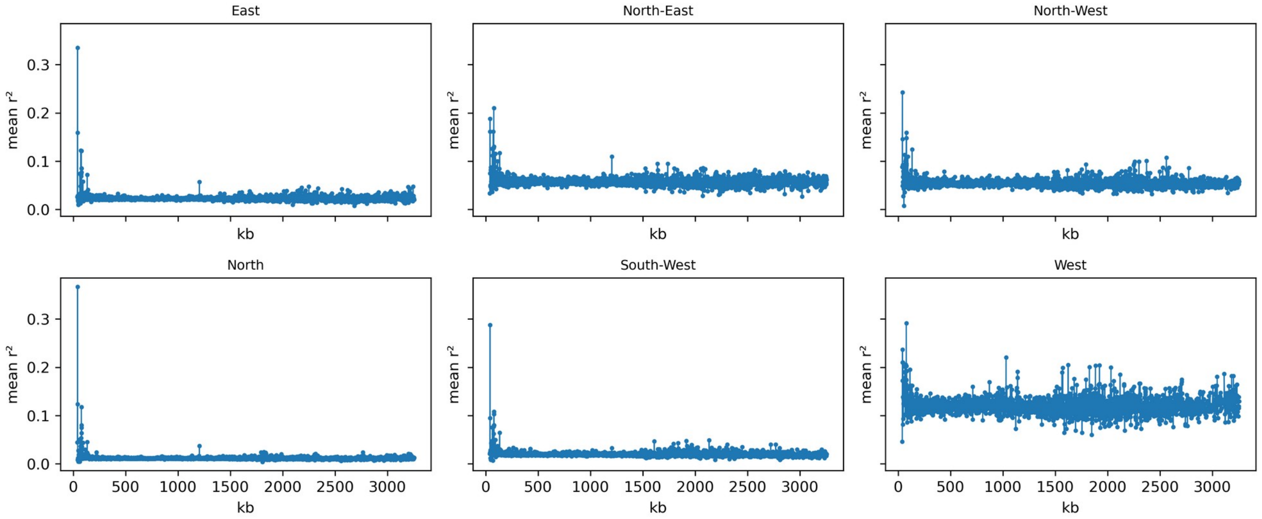


**Supplementary Figure 3. LD decay curves faceted by region.** Region-specific LD decay profiles ($r^2$ vs. distance in kb) are shown separately for each of the six geographic regions. Differences in the shape or rate of decay between regions may reflect variation in local effective population sizes, transmission intensity, and extent of inbreeding.

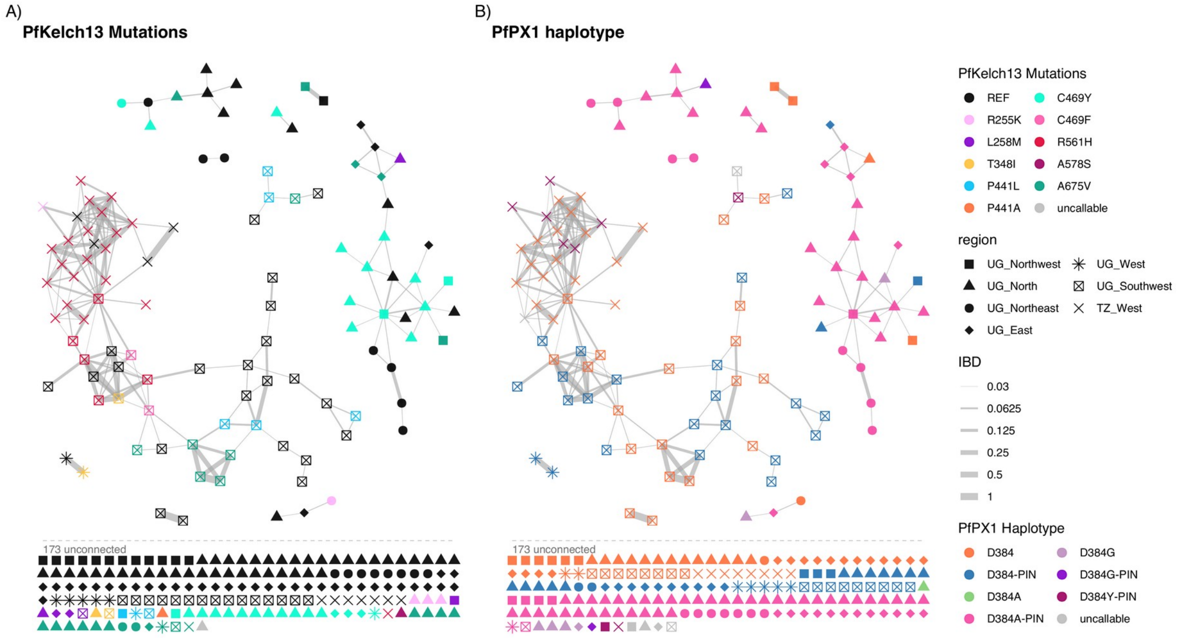


**Supplementary Figure 4. Sample-level IBD network coloured by PfKelch13 genotype, PfPX1 haplotype and region.** Each node represents an individual *P. falciparum* isolate; edges connect pairs sharing IBD >= 0.03125 (1/32). Node colour indicates the PfKelch13 genotype (wild-type vs. mutant in A), and in b) the PfPX1 haplotype (D384 variant with or without the PIN haplotype), and node shape indicates the sample's geographic region, enabling assessment of whether resistance variants cluster within or spread across regional transmission networks. The majority of the connections between the samples in the Northern regions are between isolates containing the 384A-PIN variant, are under a recent selection sweep which likely contributes to the amount of sharing being seen in those regions rather than sharing due to geographical constraint.

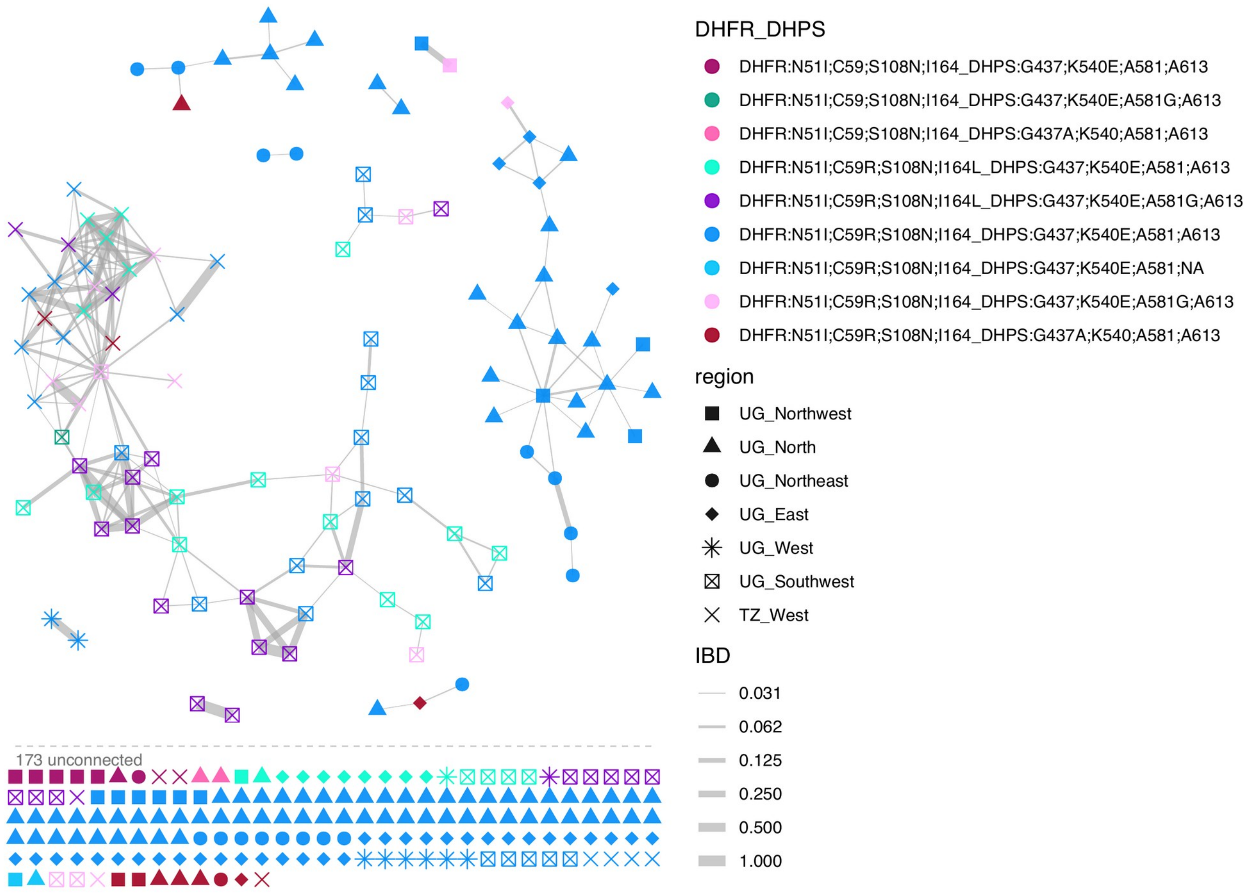


**Supplementary Figure 5. Sample-level IBD network coloured by PfDHPS/PFDHFR and region.** The nodes are coloured by the combined PfDHPS/PfDHFR sulfadoxine-pyrimethamine resistance haplotype. The network highlights the regional distribution of antifolate triple- and quintuple-mutant haplotypes and their sharing within IBD-connected clusters.

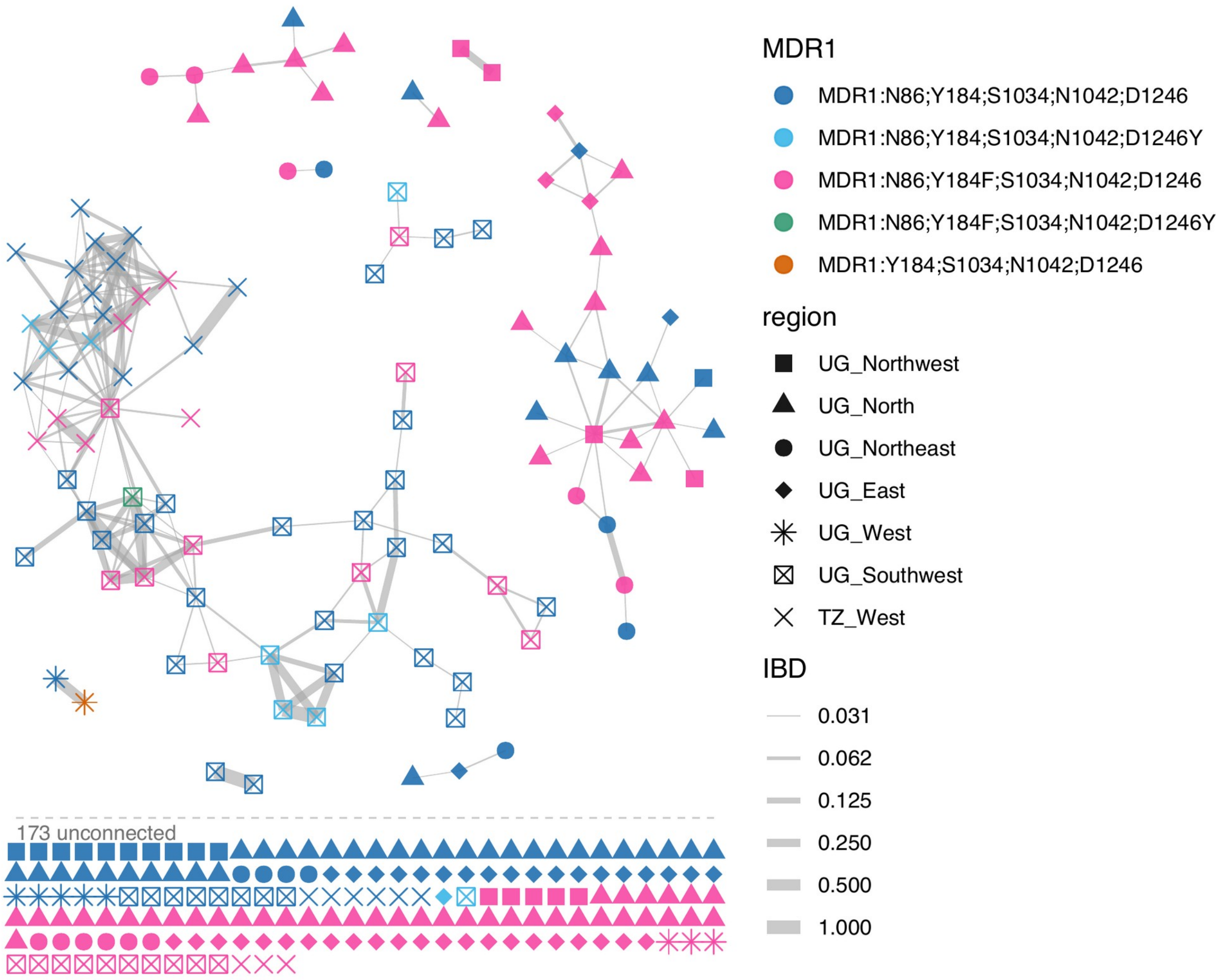


**Supplementary Figure 6. Sample-level IBD network coloured by PfMDR1 haplotype and region.** The nodes are coloured by PfMDR1 haplotype (N86Y, Y184F, D1246Y variants). The network illustrates how lumefantrine- and mefloquine-resistance associated alleles are distributed among related parasite lineages across Uganda's regions.

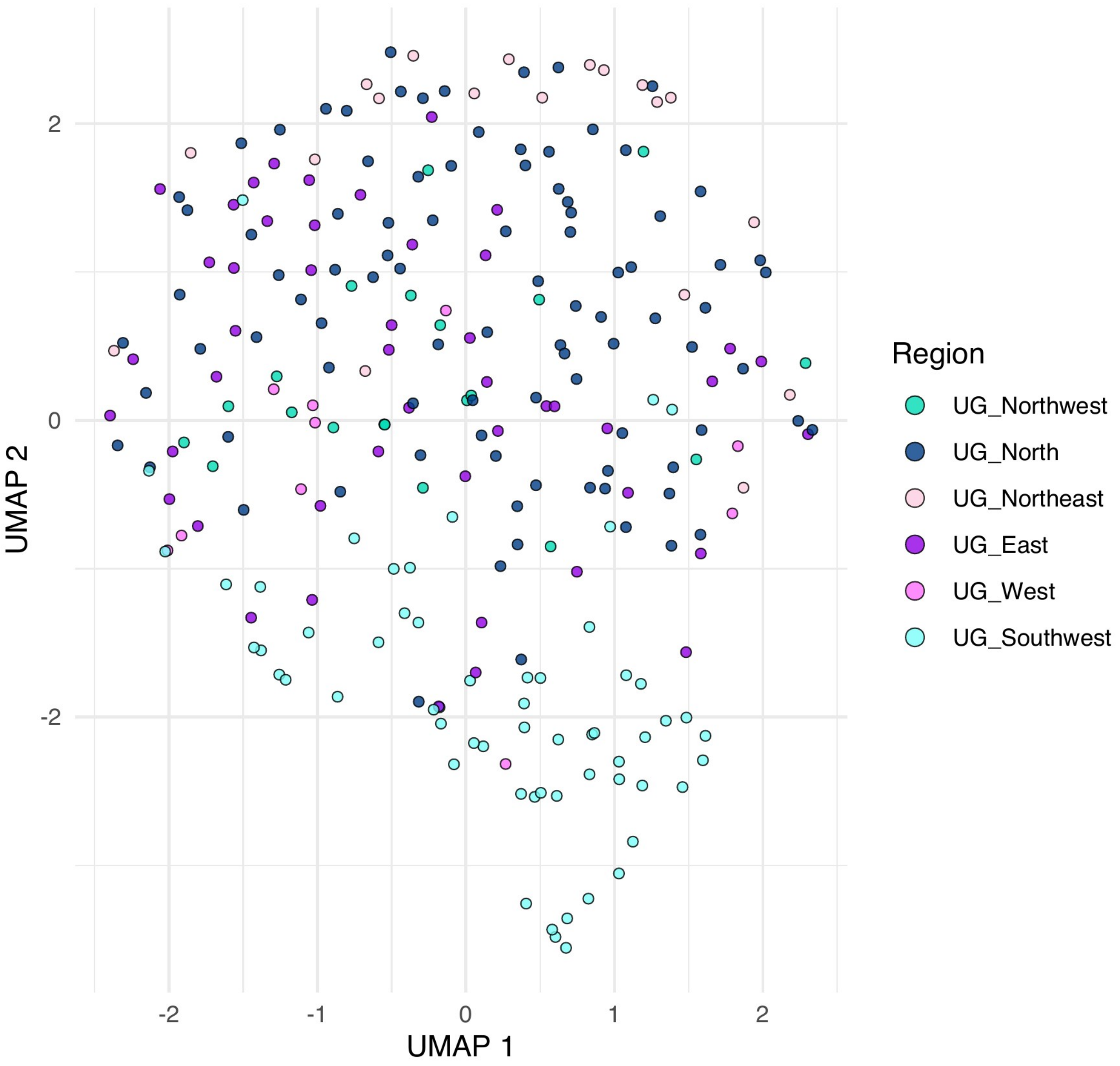


**Supplementary Figure 7. UMAP projection of Uganda-only samples coloured by region.** Uniform Manifold Approximation and Projection (UMAP) of genome-wide SNP variation restricted to the 249 Ugandan 2023 samples. Each point represents one sample, coloured by its region of collection. Clustering patterns reflect the degree of genetic differentiation among regions and identify any within-country population substructure. The only visible structure appears to be the Southwest samples which majority cluster together at the bottom of the larger cluster.

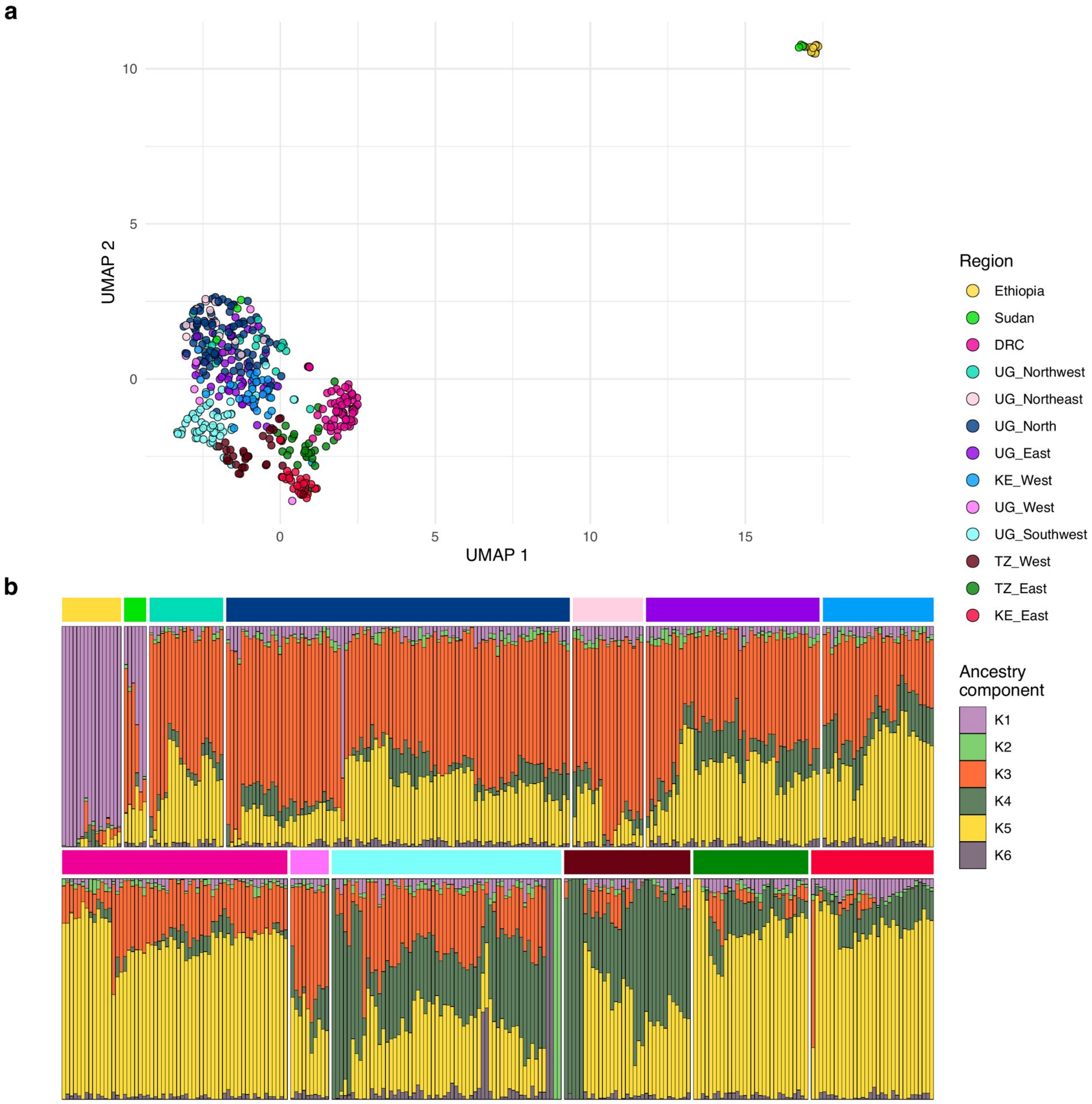


**Supplementary Figure 8. Population structure of *Plasmodium falciparum* samples from Uganda and regional comparator populations. (A)** UMAP of the samples from Uganda and comparator populations from Ethiopia, Sudan, Democratic Republic of the Congo (DRC), Kenya, and Tanzania coloured by region/population, showing genetic relationships and clustering patterns across Uganda and neighboring countries. **(B)** LEA sNMF admixture plot at K = 6 showing ancestry proportions for same samples. Each vertical bar represents one sample, and colours indicate inferred ancestry components. Three of the Sudanese parasites cluster with Uganda and one Ugandan parasite cluster with the Ethiopia/Sudan cluster, this parasite also is dominated by the K1 ancestry seen in the admixture for those countries, consistent with either with limited transmission in those regions with Northern Uganda or imported cases from travelers or refugees.

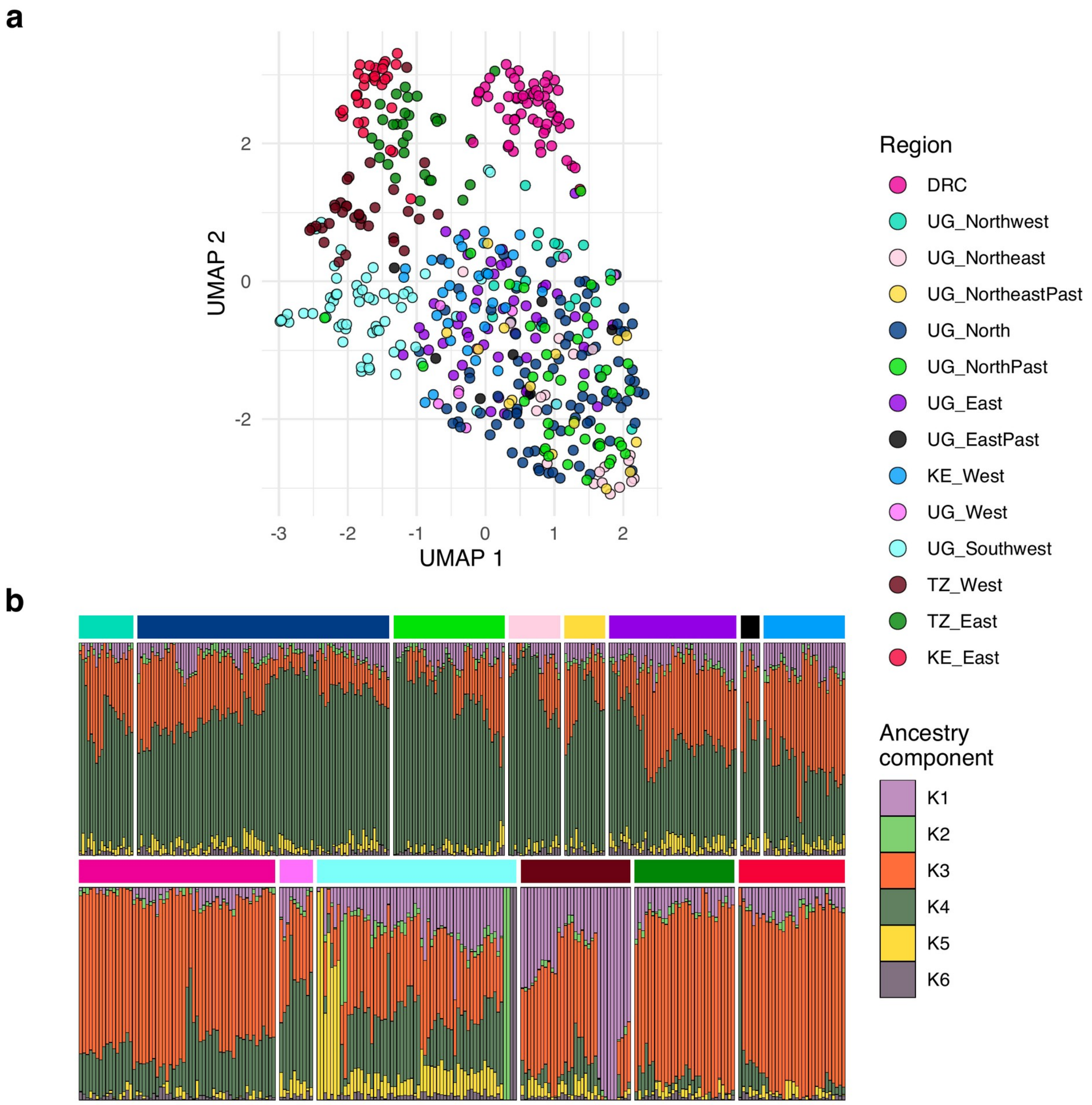


**Supplementary Figure 9. Population structure of contemporary and past *Plasmodium falciparum* samples from Uganda in the context of regional comparator populations. a)** UMAP of the samples coloured by region/group, illustrating genetic relationships between contemporary and past Ugandan parasites and their clustering relative to comparator populations from Democratic Republic of the Congo (DRC), Tanzania, and Kenya. Past data are only available for regions of Northeast, North and East and each of the past parasites have very similar ancestry components with their contemporary populations consistent with the parasite population not drastically changing over time. **b)** LEA sNMF admixture plot at K = 6 showing ancestry proportions for individual contemporary Ugandan samples, past Ugandan samples from selected regions, and comparator populations. Each vertical bar represents one sample, and colours indicate inferred ancestry components.

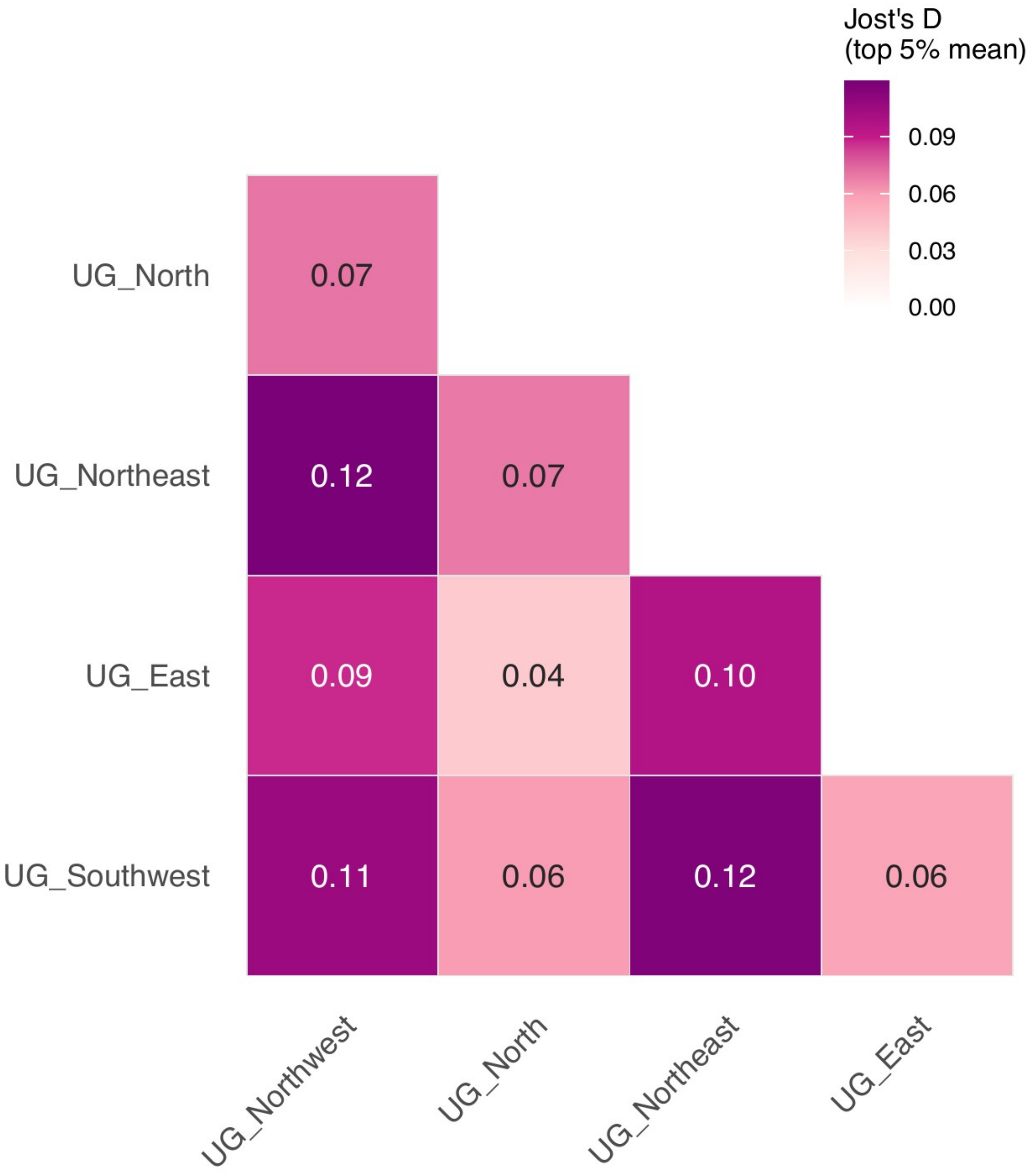


**Supplementary Figure 10. Pairwise Jost's D heatmap by region.** Colour-coded matrix of the mean pairwise top 5% of Jost's D differentiation statistics calculated between all region pairs. Higher values (darker colour) indicate greater allelic differentiation, providing a complementary measure of population structure to IBD-based analyses.

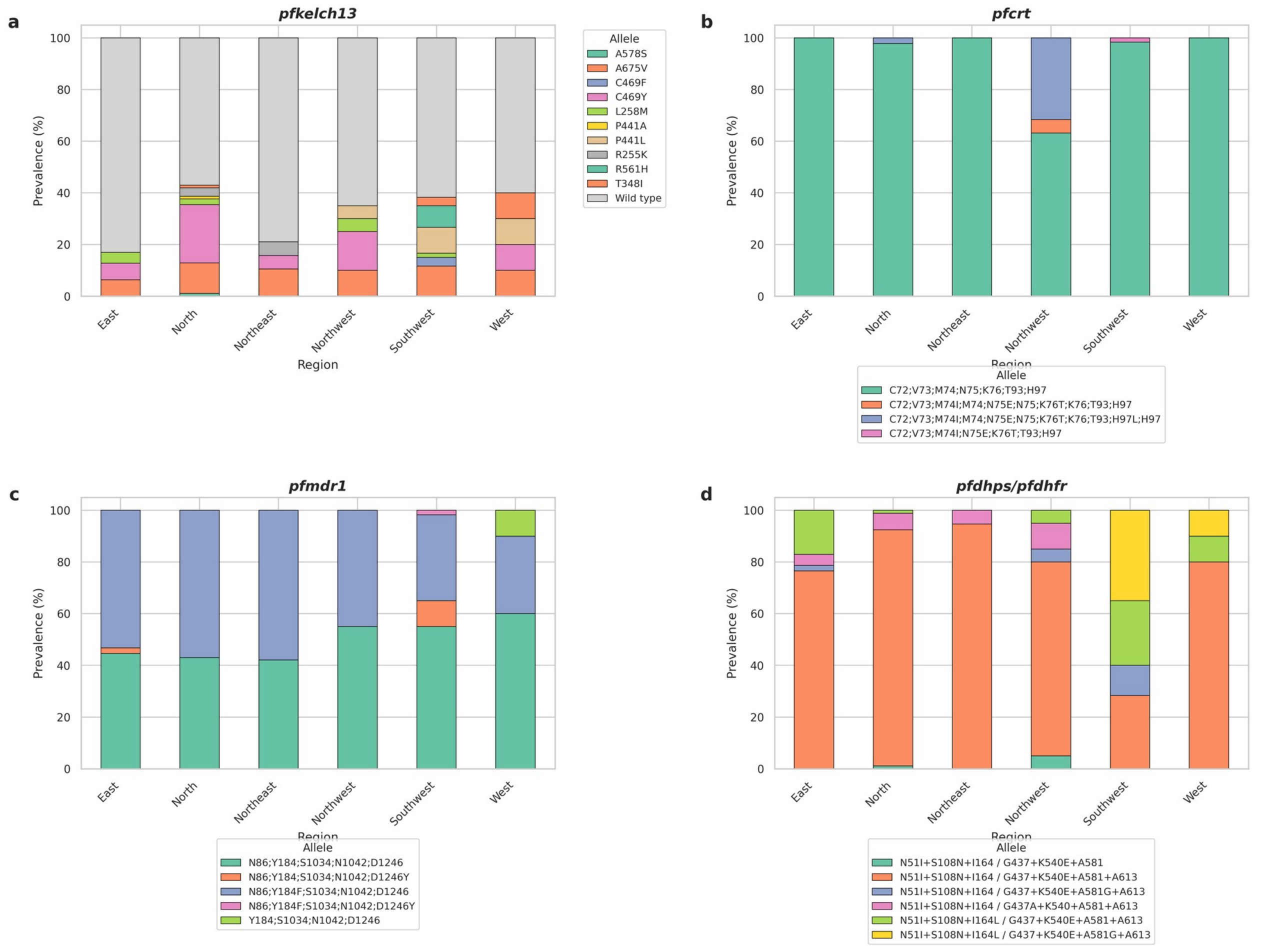


**Supplementary Figure 11. Prevalence of drug resistance markers across PfKelch13, PfCRT, PfMDR1, and PfDHFR/PfDHPS by region.** Bar charts summarizing the frequency of key antimalarial resistance mutations in each region. Markers include artemisinin partial-resistance candidates in PfKelch13, chloroquine-resistance alleles in PfCRT, multidrug-resistance alleles in PfMDR1, and antifolate-resistance haplotypes in PfDHFR and PfDHPS.

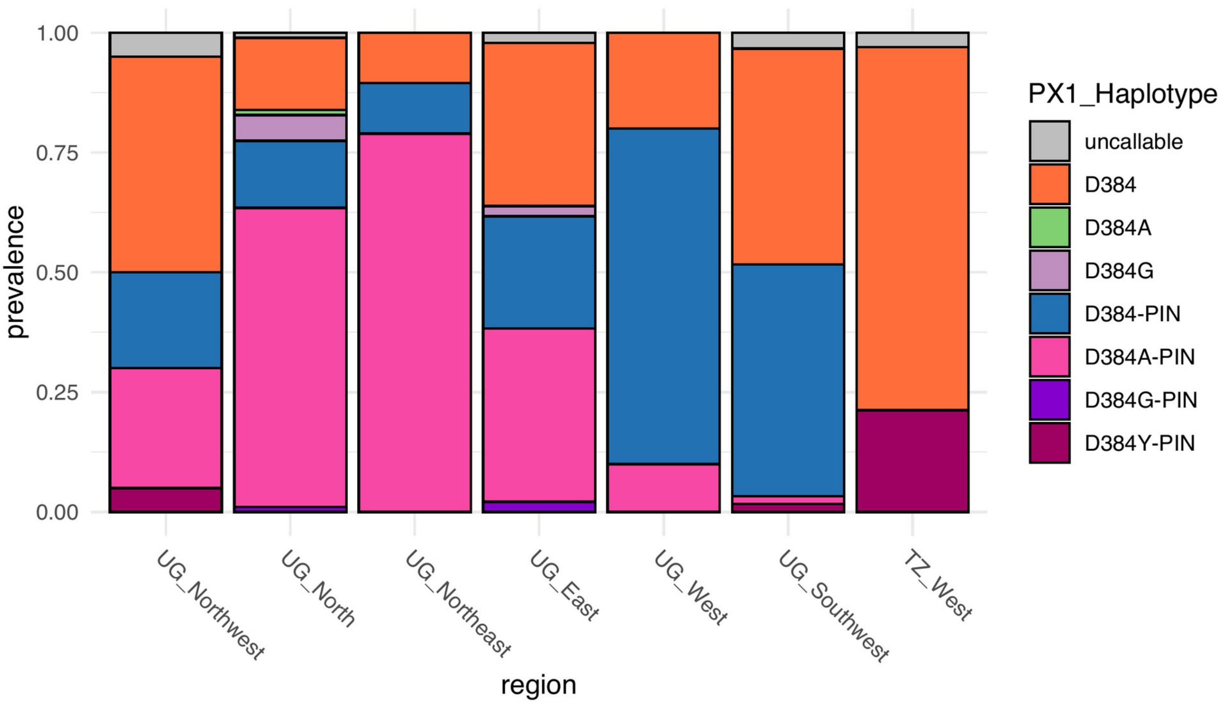


**Supplementary Figure 12. PfPX1 Haplotype Prevalence.** The prevalence of the PfPX1 D384A/G/Y, L1222P, M1701I, and D1705N haplotype across each region. The haplotypes are sorted by the presence or absence PIN. The total prevalence the PIN haplotype is the equivalent of they-axis height of the blue bar for the Ugandan samples and the dark maroon bar for the Tanzania West (TZ_West) samples.

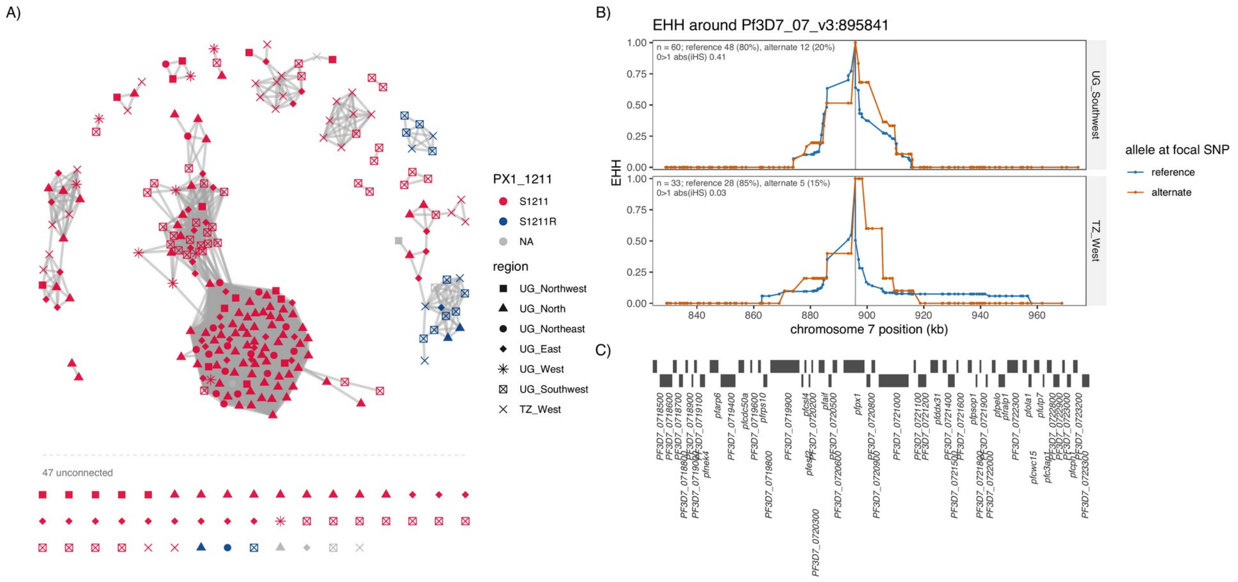


**Supplementary Figure 13. PfPX1 S1211R mutation network and EHH. (A)** The network between Uganda and Tanzania West samples. Connections are shown if samples have an IBD block that completely covers the PfPX 1211 corresponding genomic locus, colored by whether or not they contain S1211R and the shapes of the nodes are the region they are from. The S1211R mutation is found only in non-PIN *pfpx1* isolates. Almost all the S1211R samples have an IBD block with another sample. (**B)** Is the EHH for Uganda Southwest and Tanzania West. There is not a large extended haplotype around this mutant which would be consistent with the mutation not being selected for and therefore not sweeping but instead simply drifting. This variant was uncommon among African isolates (3.0% prevalence in Burkina Faso, 2.7% in Nigeria, 1.5% Tanzania, and <1% elsewhere) in the MalariaGEN Pf8 dataset but reached 33% prevalence in Indonesia and 12% in Papua New Guinea.

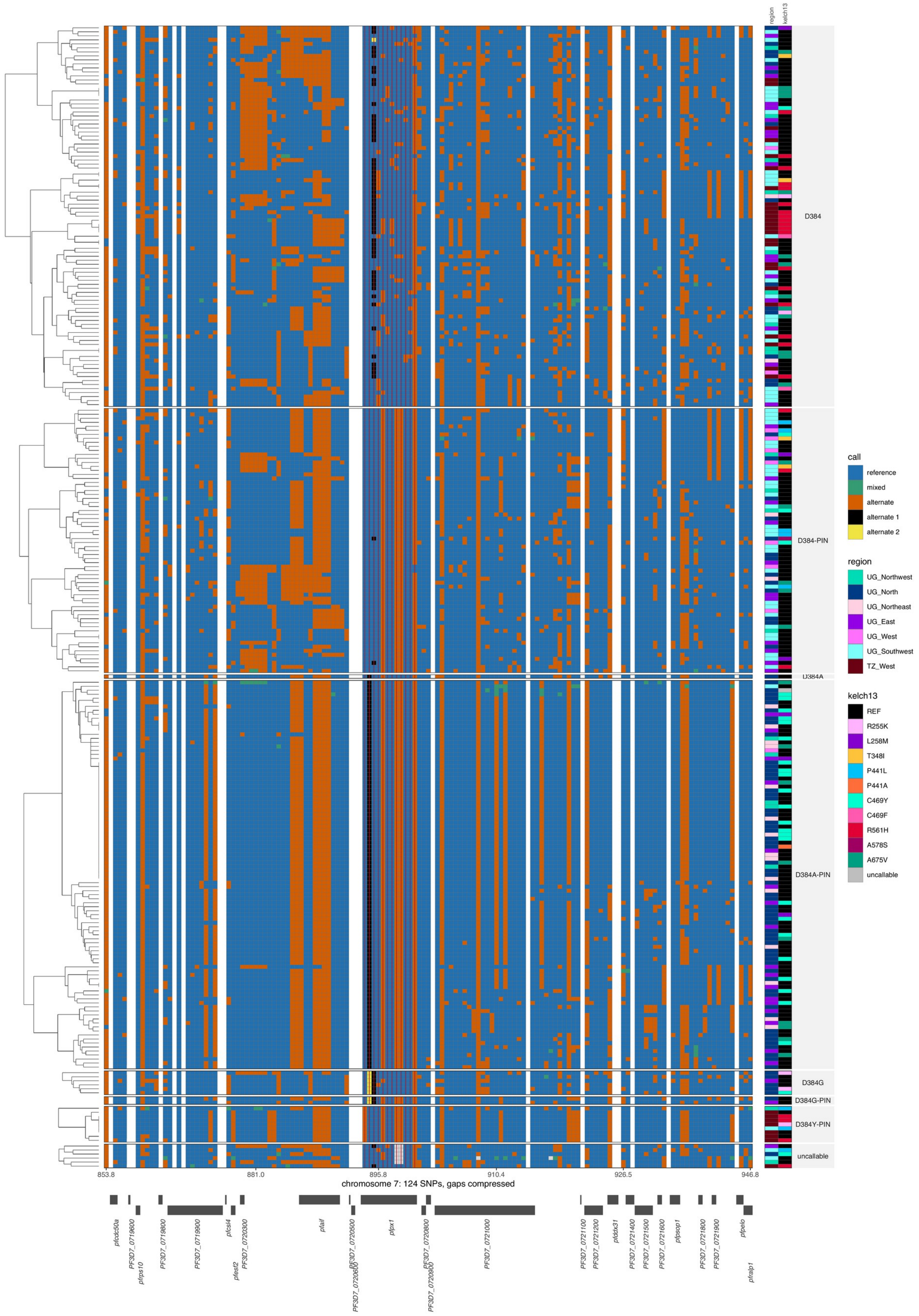

**Supplementary Figure 14. Local genotypes around *pfpx1.*** Each column is a called polymorphic single nucleotide position centered around *pfpx1,* extending left and right ~50kb. Each row is a sample clustered based on the variant calls and split by PfPX1 haplotype. Columns to the right annotate the regions the isolates come from and what PfKelch13 mutations they carry. D384A-PIN demonstrates a wider shared haplotype than D384-PIN. D384A-PIN also carries the majority of the PfKelch13 C469Y mutations. Like D384A-PIN, D384Y-PIN has a wide haplotype, but is predominately found in the west of Tanzania. D384G is found with PIN for 2 out of the 8 total D384G isolates and was only detected in the North and East regions.

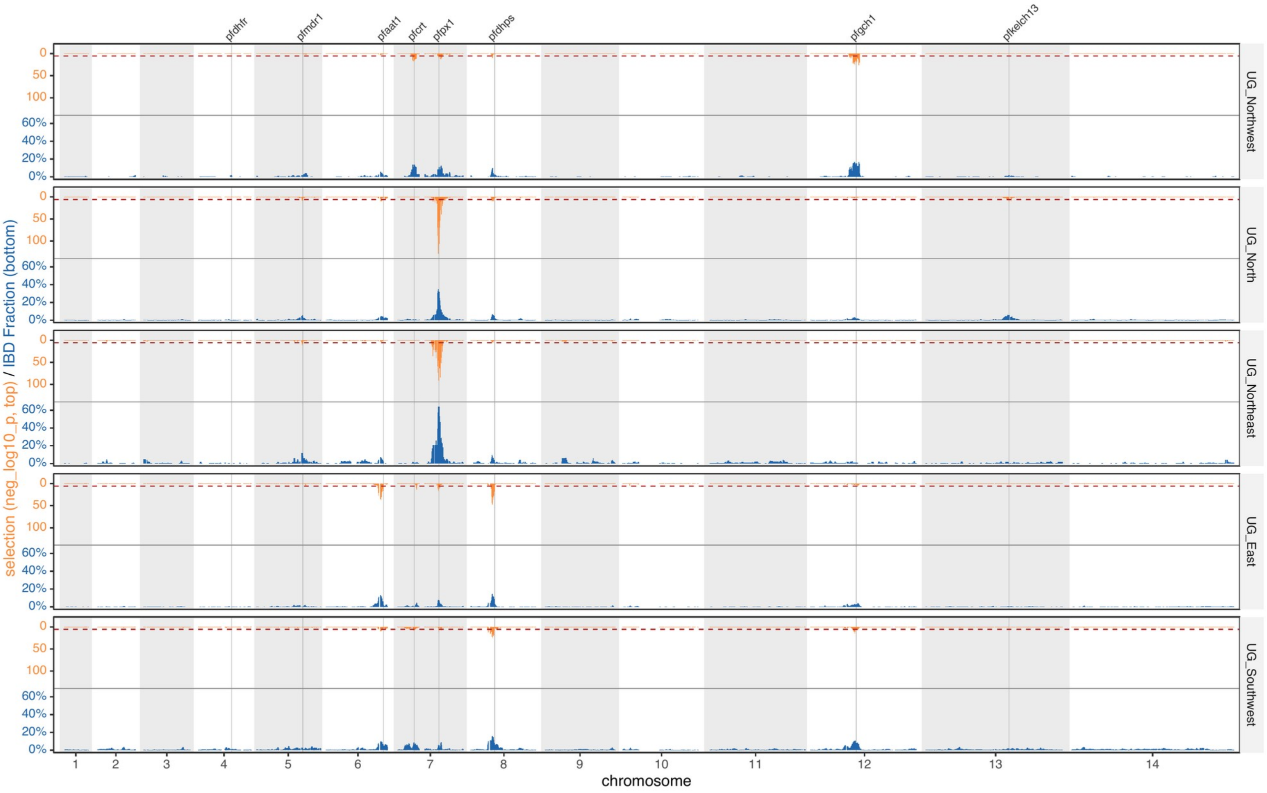


**Supplementary Figure 15. Genome-wide IBD-based selection/pairwise sharing scans IBD by region.** Above the central horizontal axis (orange), the IBD based selection statistic XiR,s SNPs with the red line marking where above SNPs were significant based on Bonferroni. Below the central horizontal axis (blue), the proportion of isolate pairs sharing IBD at each SNP position within each region. Candidate selection peaks corresponding to known drug resistance loci are labeled.

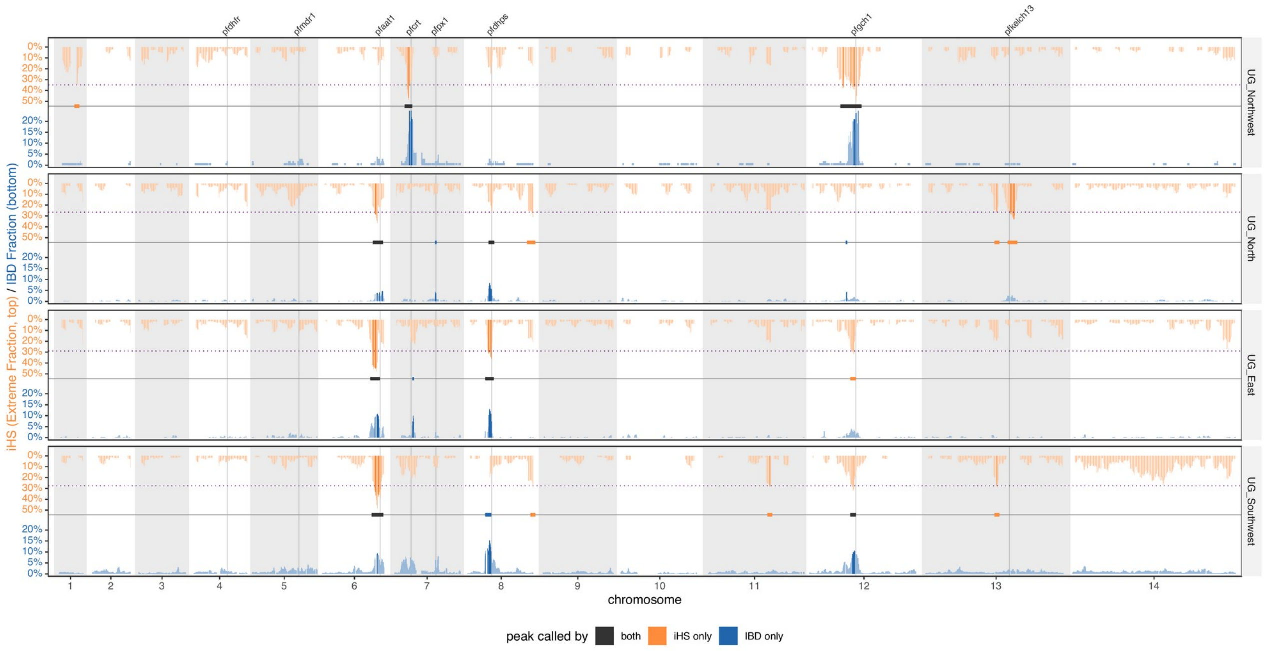


**Supplementary Figure 16. Haplotype-based selection with Integrated Haplotype Score (iHS) mirrored against IBD sharing, by region all chromosomes.** One panel per region. Above the axis (orange), the fraction of SNPs with |iHS| > 2 in 100-kb windows stepped every 10 kb; below (blue), the fraction of sample pairs sharing IBD at each SNP. Per-SNP iHS is dominated by sampling variance whereas a sweep elevates a contiguous run of SNPs, so selection is summarised as the proportion of extreme SNPs per window rather than SNP by SNP (Voight et al. 2006). iHS was computed in R package rehh, unpolarized so only |iHS| is interpreted, and standardised within 10 quantile bands of minor-allele frequency. Drug-resistance loci are marked. A purple dotted line marks the height of the top 1% genome wide of iHS peaks. The dominant signal is at *pfpx1* (chromosome 7), extreme in both tracks (iHS 67%, IBD 34% in UG_North; IBD 62% in UG_Northeast). Two further loci are elevated in both tracks in UG_Northwest: *pfgch1* (chromosome 12; iHS 42%, IBD 15%) and *pfcrt* (chromosome 7; iHS 35%, IBD 14%), the latter also moderately in UG_Southwest (iHS 20%, IBD 9%). Three loci are elevated in iHS with little corresponding IBD sharing: pfkelch13 (chromosome 13, 44% in UG_North), *pfaat1* (chromosome 6, 43% in UG_Southwest) and *pfdhps* (chromosome 8, 43% in UG_East). *pfdhfr* shows no signal in either track.

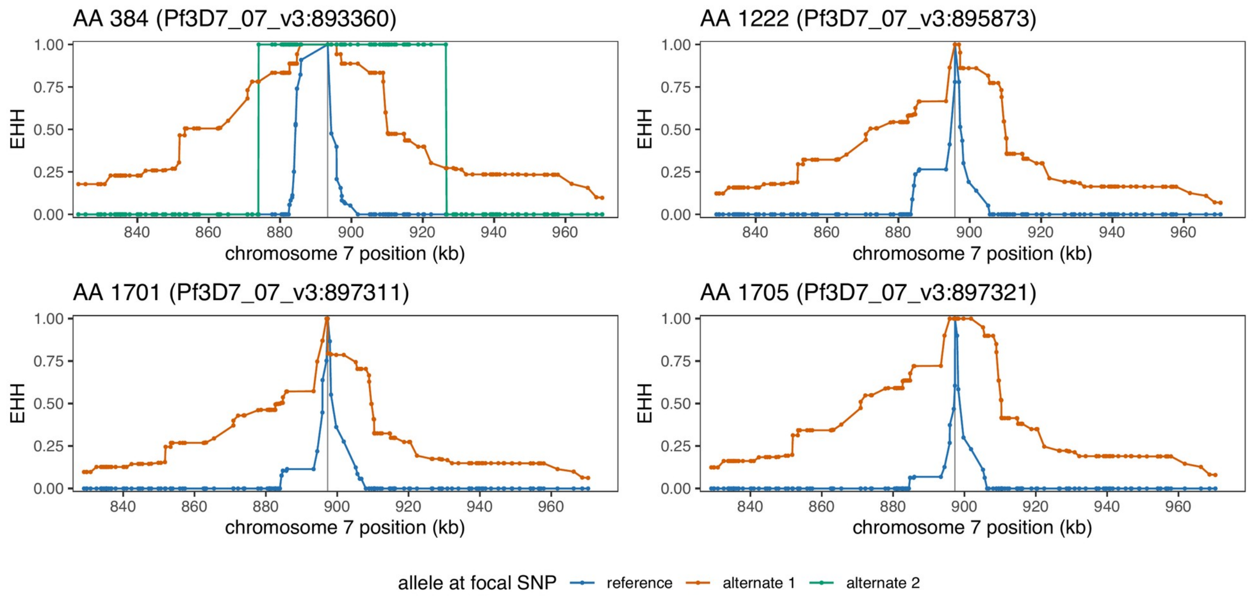


**Supplementary Figure 17. EHH plots of PfPX1 variants in 2020-2021 Ugandan Isolates.** Extended haplotype plots for Uganda 2020-2021 isolates of the 4 important residues seen in the sweeping D384A-PIN haplotype (384 top left, 1222 top right, 1701 bottom left, 1705 bottom right) created by wrapping R package rehh. Blue lines are the reference, and orange is the first alternative. Positions 1222, 1701, and 1705 are biallelic SNPS and only have 1 alternative, while 384 is a multiallelic SNP with a 2nd alternative allele (green line), for 384 the orange line is the allele coding for D384A, the green line is the alternative coding for D384G, the non-PIN associated 384 amino acid. The iHS score is based on the ratio of the area under the allele of interest to the reference allele. The longer the alternative allele's haplotype and/or the shorter the reference haplotype, the greater the iHS. The iHS scores from highest to lowest, D1705N 4.78 p=6.88e-06; D384A 3.84 p=4.92e-04; L1222P 3.72 p=7.91e-04; M1701I 3.55 p=0.00157; D384G 1.4 p=0.65.

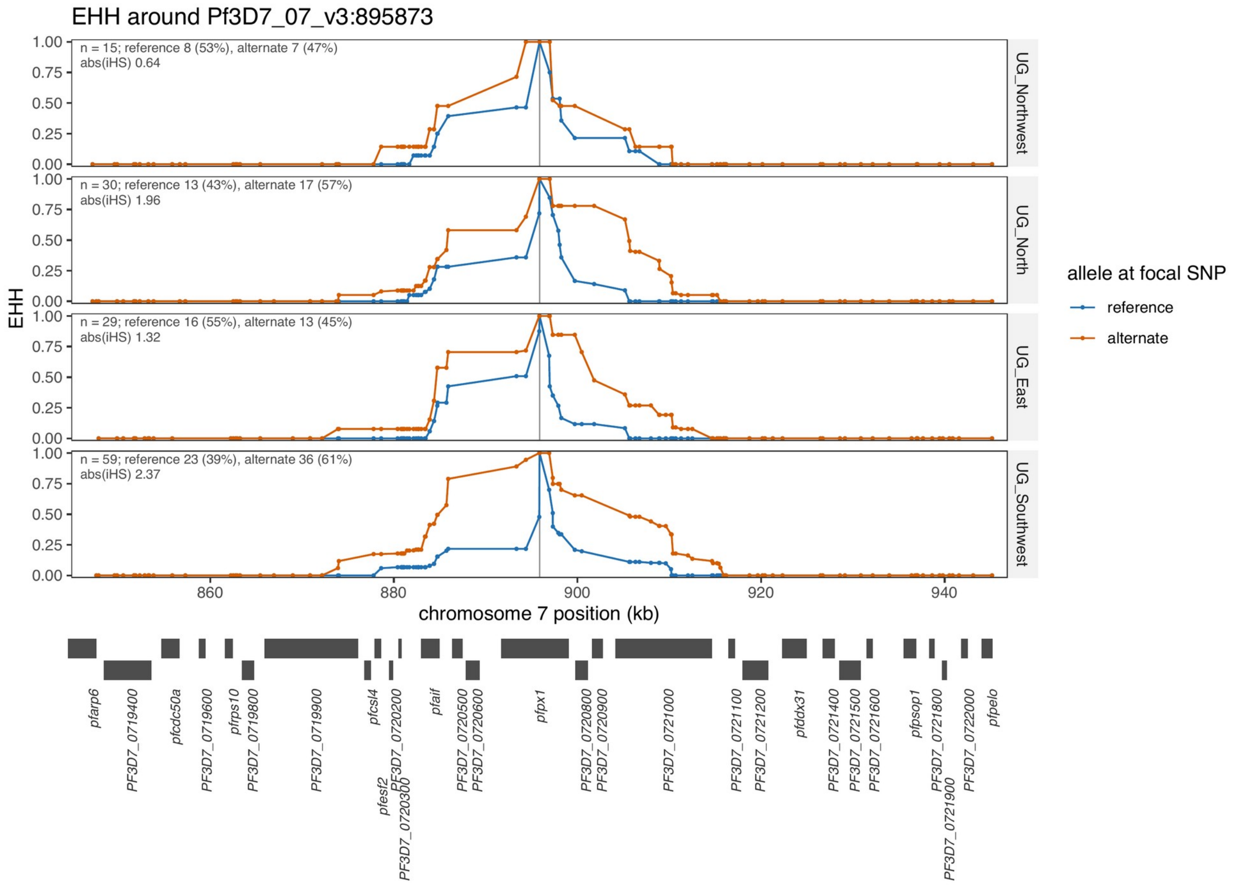


**Supplementary Figure 18. EHH plots of PfPX1 variants with D384A-PIN haplotypes removed.** After removing D384A-PIN haplotypes from the population and recalculating iHS and replotting EHH for 1222P, a very similar extended haplotype with similar mild iHS scores can be seen. There is also a similar prevalence across regions ranging from 45-60%. All of which would be consistent with a similar D384-PIN background from with D384A-PIN then arose.

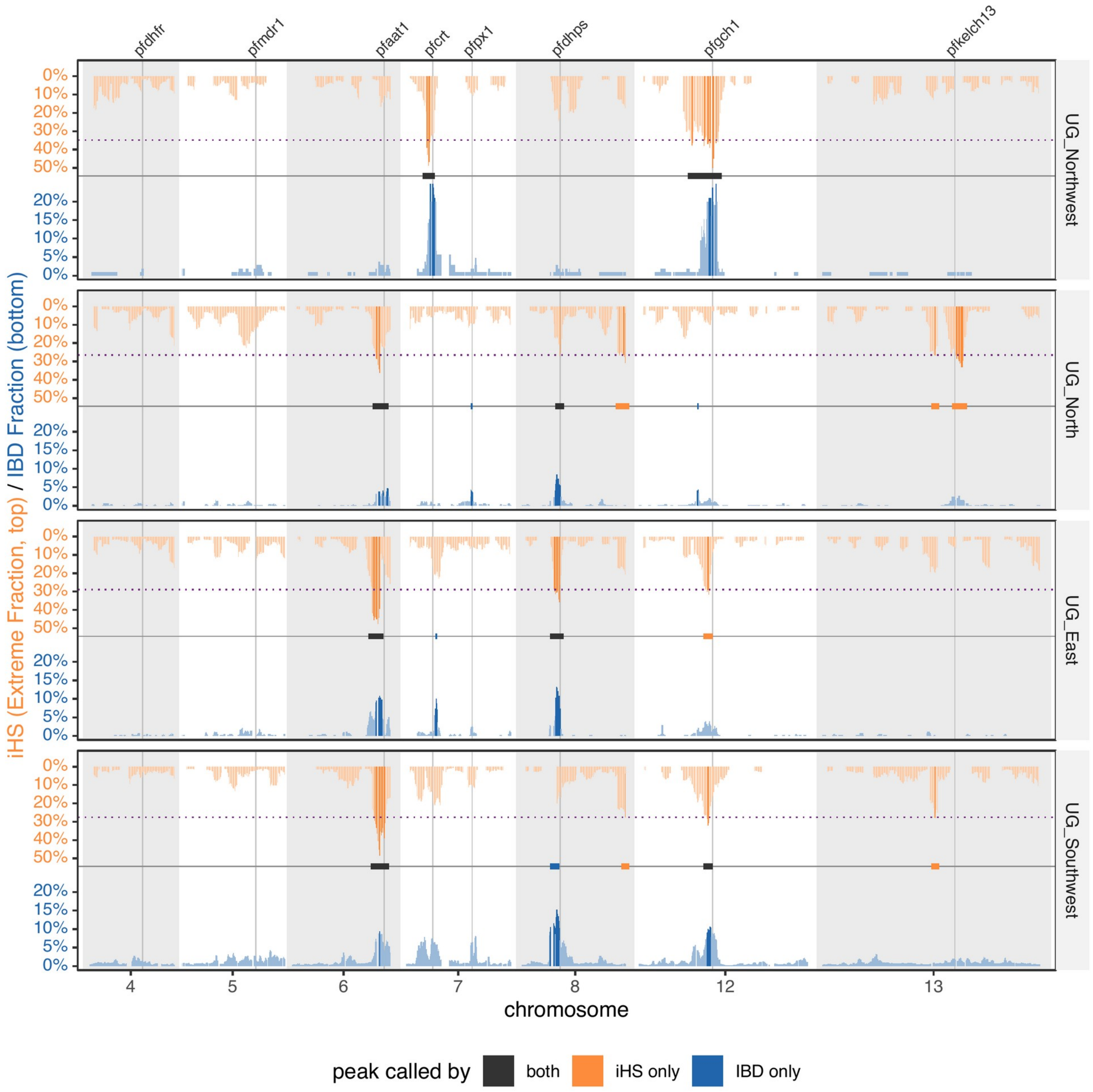


**Supplementary Figure 19. Haplotype-based selection with Integrated Haplotype Score (iHS) mirrored against IBD sharing with D384A-PIN removed.** After removing D384A-PIN from the population and recalculating IBD and iHS there is almost no significant signal detected at *pfpx1* loci despite a prevalence of 45-60% D384-PIN left in the remaining population.

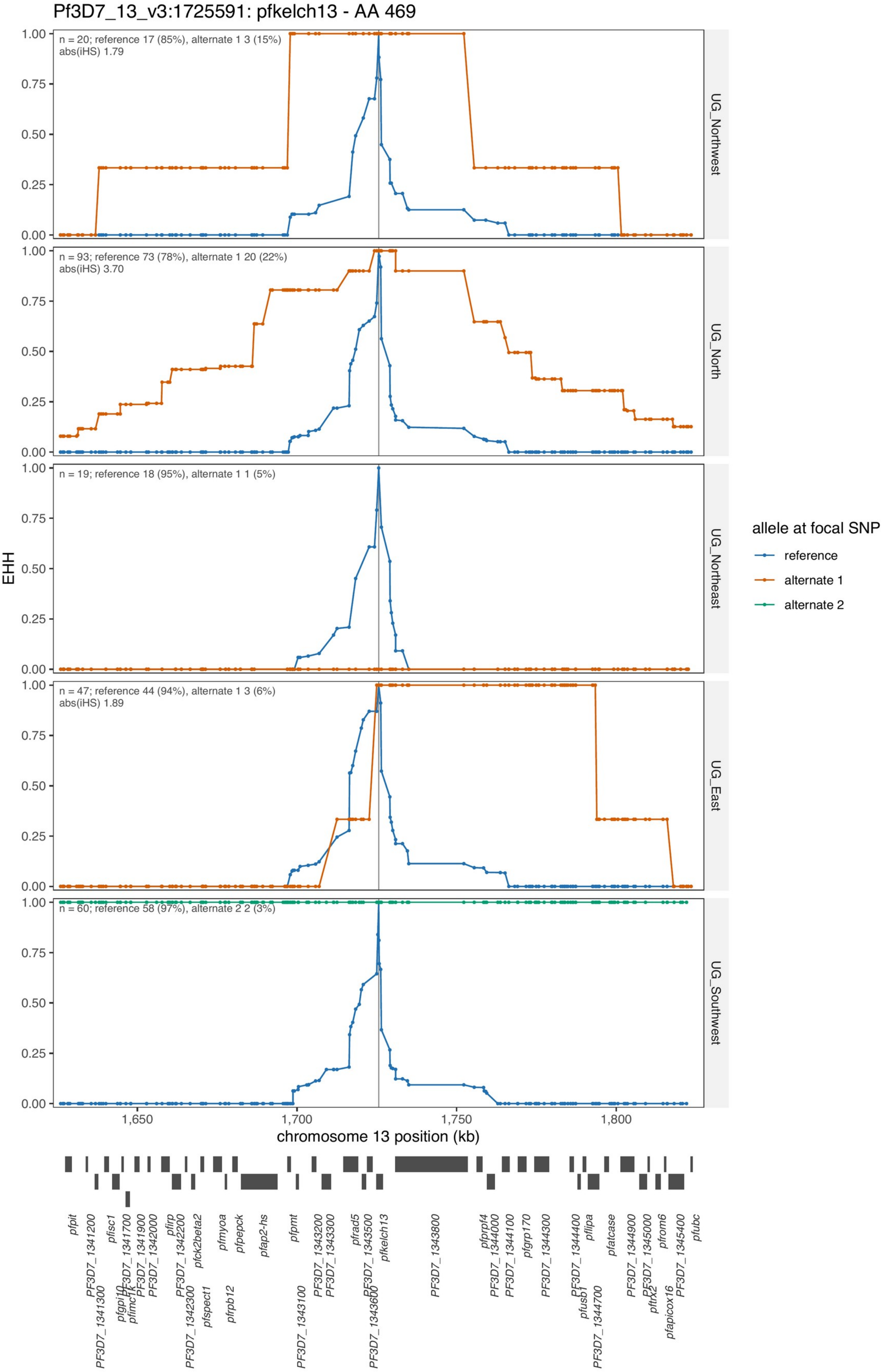


**Supplementary Figure 20. EHH for PfKelch13 C469Y/F.** Long haplotypes seen around C469Y (alternative 1) in the North (|iHS| 3.70) and Northwest (|iHS| 1.79). C469F (alternative 2) only seen in two isolates and share the same haplotype.

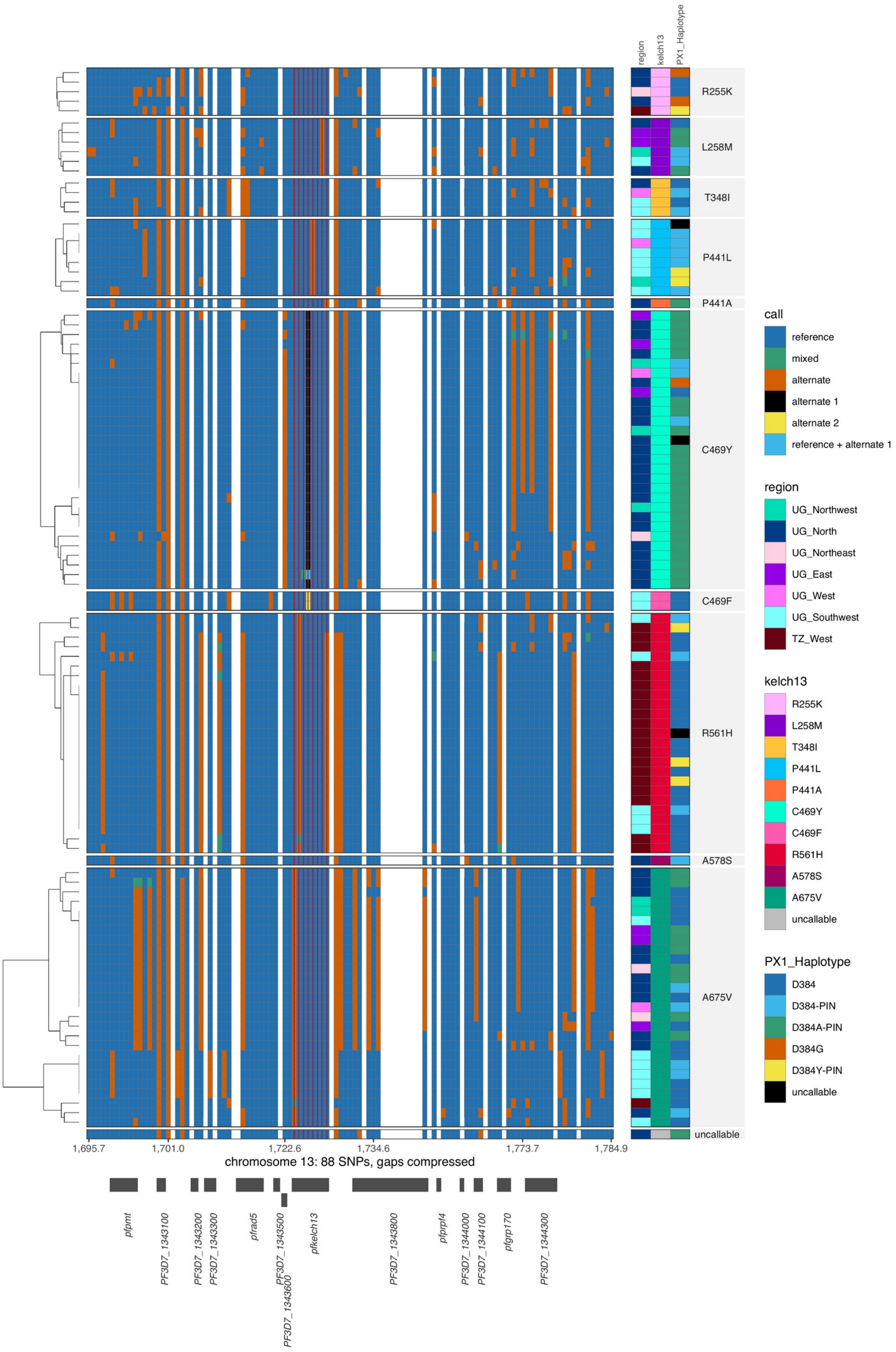


**Supplementary Figure 21. *pfkelch13* genotypes.** Each column is a variant SNP called position centered around *pfkelch13* extended left and right 100kb total. For every 2kb between SNPs that lack a SNP, a white column is placed to achieve relative genomic distance. Each row is a sample clustered based on the variant calls and split by their *pfkelch13* mutants (isolates containing reference left out). Distinctive haplotypes are found for several variants including A675V and R561H. R561H is found within Uganda Southwest and Tanzania West with at least two haplotypes.

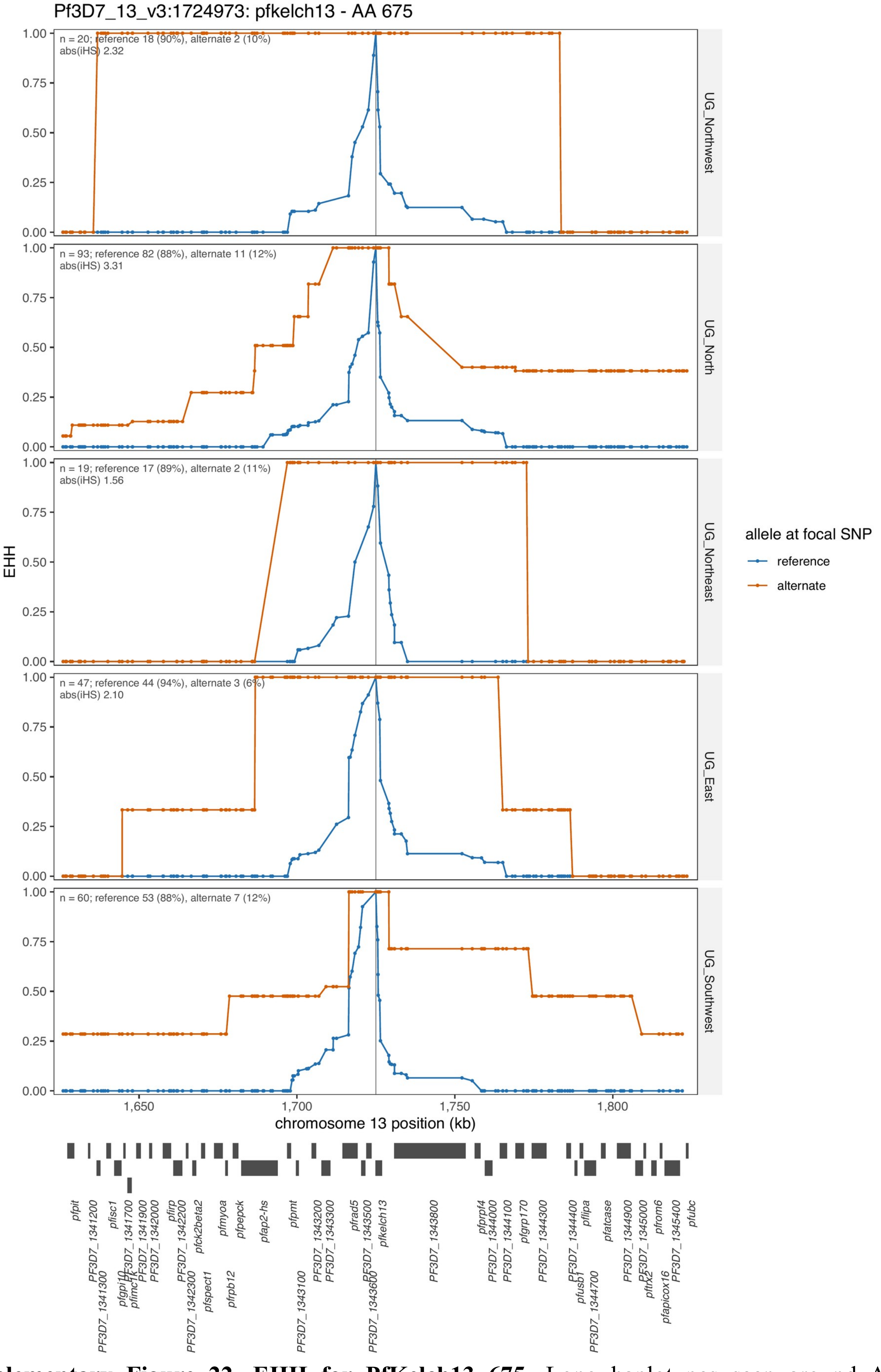


**Supplementary Figure 22. EHH for PfKelch13 675.** Long haplotypes seen around A675V (alternative 1) in the Northern/East region and Northwest as well for the haplotype in the Southwest.

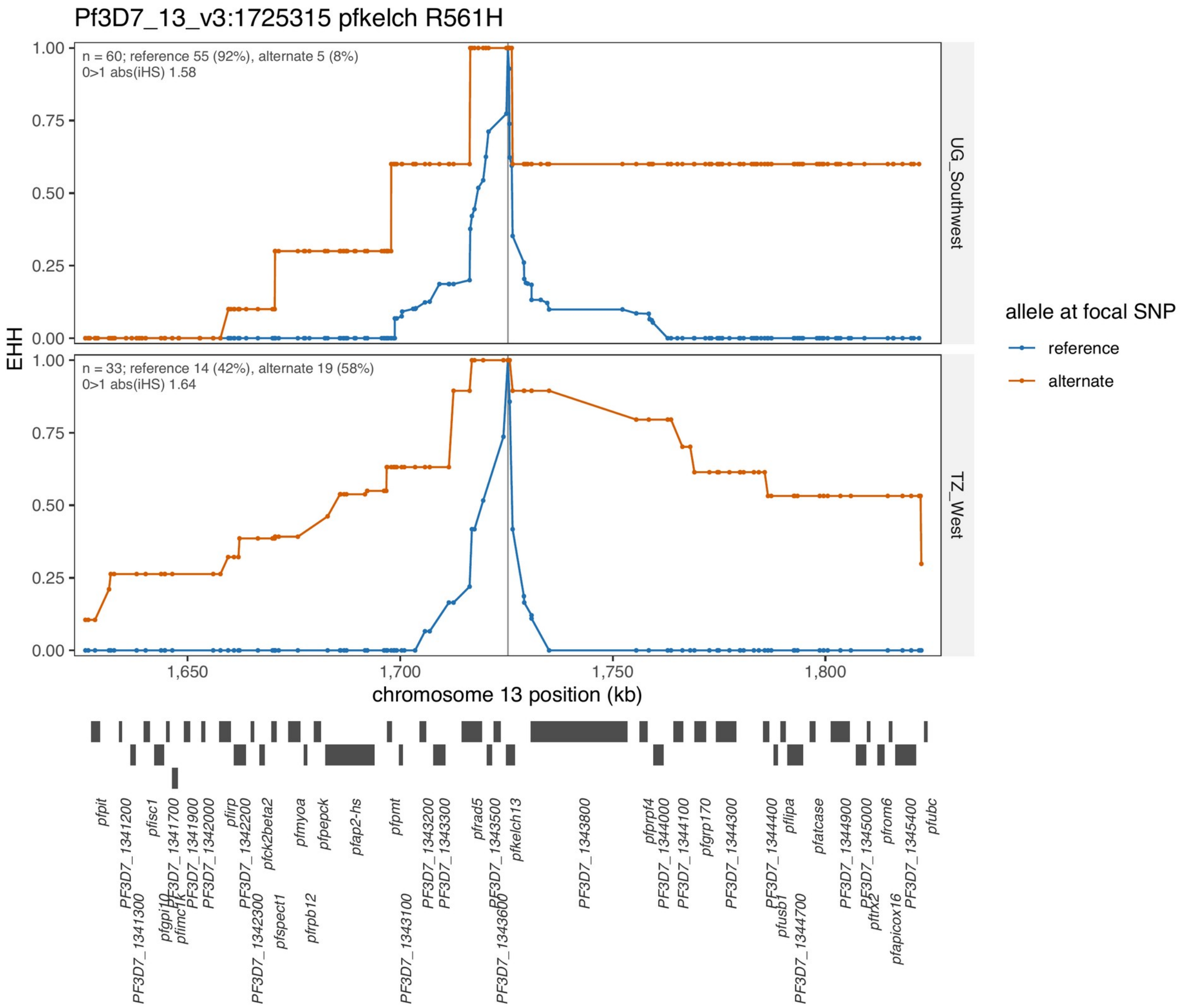


**Supplementary Figure 23. EHH for PfKelch13 R561H.** The EHH for the locus encoding for PfKelch13 R561H in Uganda Southwest and Tanzania West.

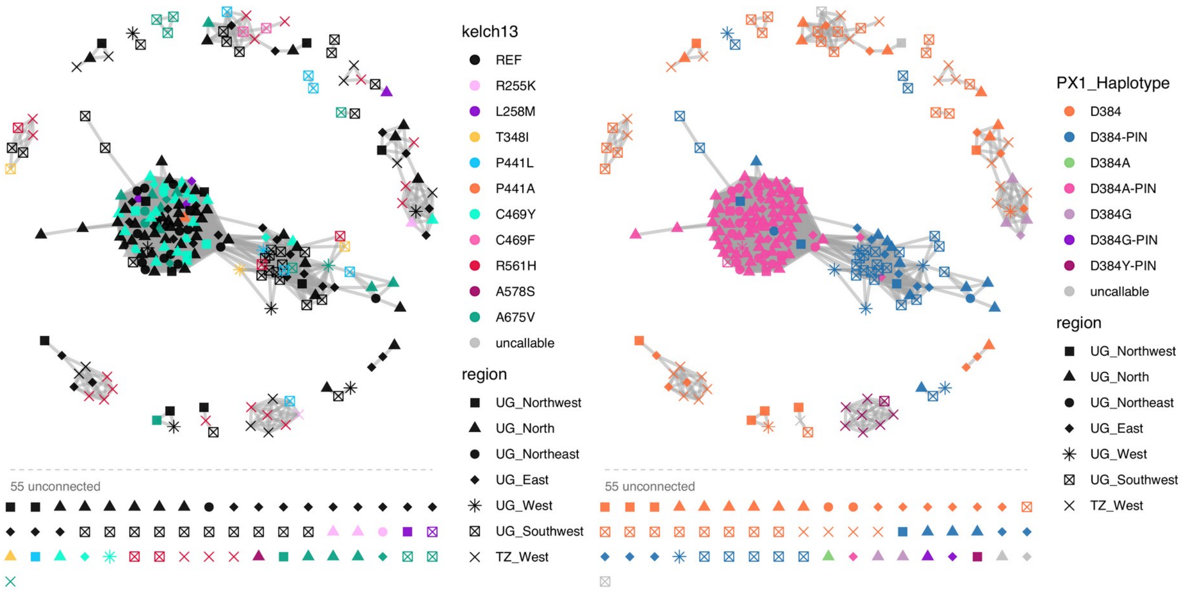


**Supplementary Figure 24. PfPX1 networks colored by PfKelch13 mutations and by PX1 Haplotype.** Association between the D384A-PIN vs D384-PIN vs everything else for PfKelch13 markers of artemisinin partial resistance (C469F, and A675V) was assessed countrywide by Fisher's exact test. Within the northern pool (North/Northwest/Northeast/East), where the D384A-PIN haplotype is essentially confined (81/83 isolates), C469Y was enriched on the D384A-PIN background relative to all other callable haplotypes (28.4% vs 6.3%; OR 5.8, 95% CI 1.8–24.4; BH-adjusted p = 0.004) and showed the same trend relative to D384-PIN specifically (28.4% vs 7.7%; OR 4.7, 95% CI 1.0–44.3; adjusted p = 0.067); A675V was unassociated with any PfPX1 variation. Although not statistically significant, 7 of the 8 P441L isolates identified countrywide carried the PIN haplotype; the small number of P441L observations limits the power of this comparison.

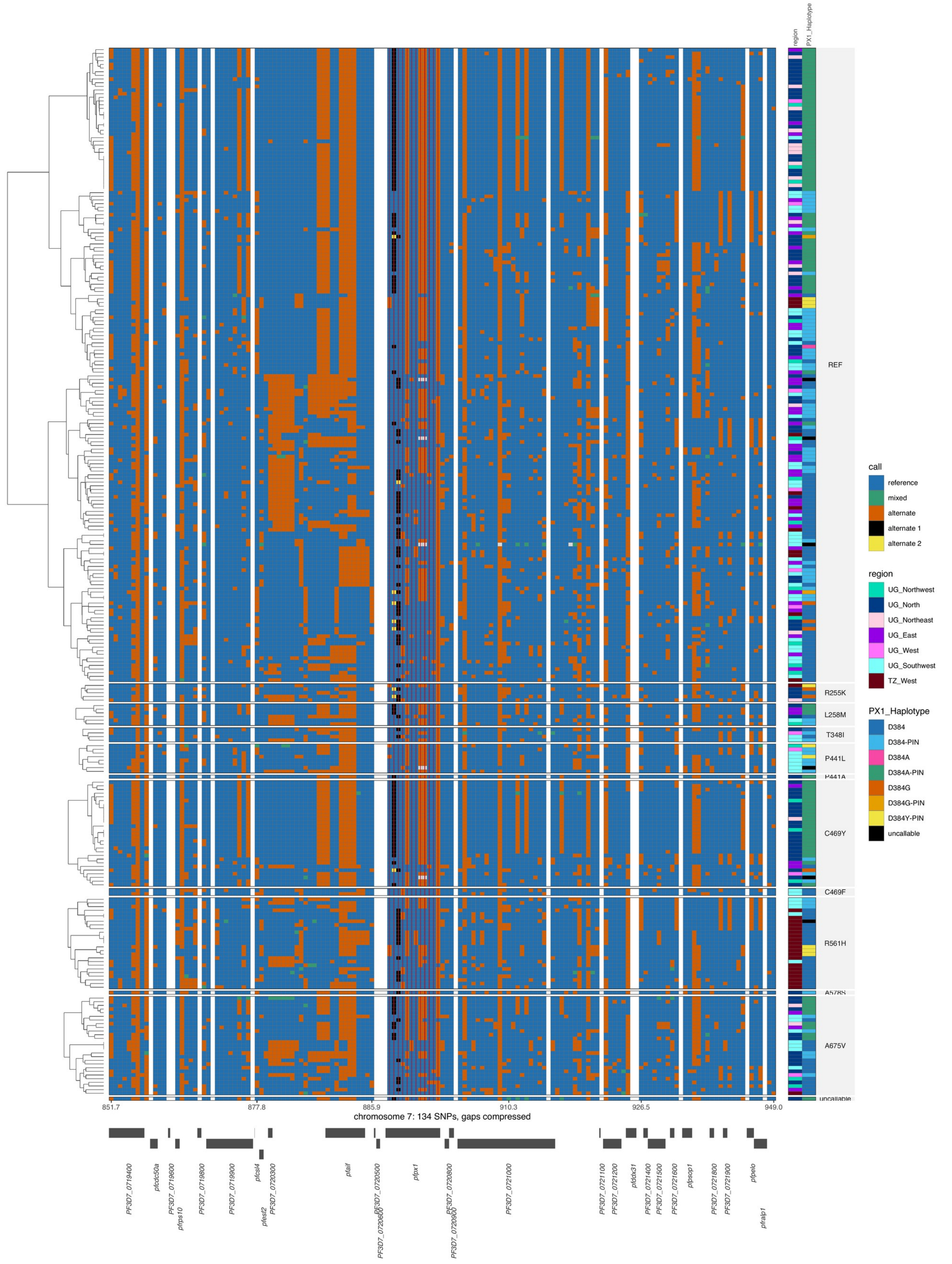


**Supplementary Figure 25.** ***pfpx1* genotypes sorted by PfKelch13 mutations.** The genotypes surrounding the *pfpx1* location split by PfKelch13 mutation. C469Y has almost all 384A-PIN haplotype. Seven out of 8 of the P441L have the PIN haplotype as well though the haplotypes for this mutation do not seem as wide as C469Y.

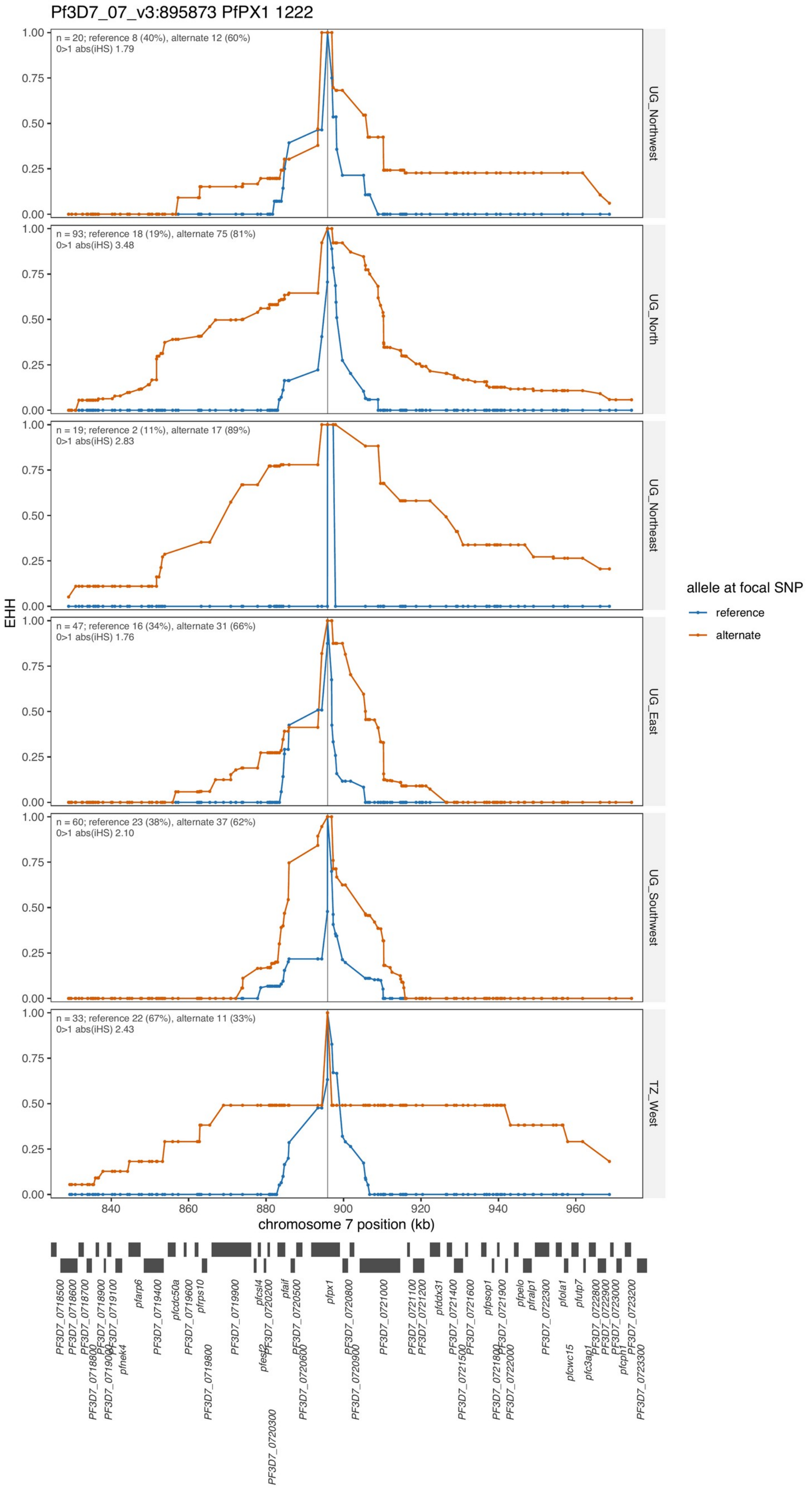


**Supplementary Figure 26. Extended haplotype around *pfpx1* L1222P.** Plotting the extended haplotype around the genomic locus that encodes for PfPX1 L1222P (variant P of the PIN haplotype). The haplotype is much wider in the North and Northeast where D384A-PIN is concentrated.

**Supplementary Tables**

**Supplementary Table 1.** Summary of genome-wide genetic diversity metrics (Nucleotide diversity, expected heterozygosity, and haplotype diversity) across six Ugandan regions.

| Region | *n* | Nucleotide diversity ($\pi \times 10-4$) | Expected heterozygosity (*He*) | Haplotype diversity |
|---|---|---|---|---|
| East | 47 | 2.34 | 0.149 | 0.966 |
| North | 93 | 2.36 | 0.152 | 0.991 |
| Northeast | 19 | 2.30 | 0.145 | 0.850 |
| Northwest | 20 | 2.30 | 0.143 | 0.854 |
| Southwest | 60 | 2.39 | 0.154 | 0.973 |
| West | 10 | 2.27 | 0.136 | 0.714 |

**Nucleotide diversity ($\pi$)**: average number of pairwise nucleotide differences per site among sampled genomes; **Expected heterozygosity (*He*)**: probability that two randomly selected alleles at a locus are different; **Haplotype diversity**: probability that two randomly selected haplotypes are different.

**Supplementary Table 2. Metadata for comparator and historical Plasmodium falciparum samples included in the study**

| Sample | District/Zone | Region | Country | African region | Collection year | Sequence accession |
|---|---|---|---|---|---|---|
| PZ0001-CW | Apac | Uganda_North-Past | Uganda | East Africa | 2010 | ERR063571 |
| PZ0004-CW | Apac | Uganda_North-Past | Uganda | East Africa | 2010 | ERR063574 |
| PZ0007-C | Apac | Uganda_North-Past | Uganda | East Africa | 2010 | ERR063577 |
| PZ0008-CW | Apac | Uganda_North-Past | Uganda | East Africa | 2010 | ERR063578 |
| PZ0010-CW | Apac | Uganda_North-Past | Uganda | East Africa | 2010 | ERR063580 |
| PZ0015-C | Apac | Uganda_North-Past | Uganda | East Africa | 2010 | ERR216476 |
| Pf003_HCMHWCCXY_L2 | Mfangano | Kenya_West | Kenya | East Africa | 2015 | ERR6908269 |
| Pf004_HFKVJCCXY_L6 | Mfangano | Kenya_West | Kenya | East Africa | 2015 | ERR6908270 |
| Pf005_HFKVJCCXY_L6 | Mfangano | Kenya_West | Kenya | East Africa | 2015 | ERR6908271 |
| Pf006_HCMHWCCXY_L2 | Mfangano | Kenya_West | Kenya | East Africa | 2015 | ERR6908272 |
| Pf007_HFKVJCCXY_L6 | Mfangano | Kenya_West | Kenya | East Africa | 2015 | ERR6908273 |
| Pf011_HCMHWCCXY_L2 | Mfangano | Kenya_West | Kenya | East Africa | 2015 | ERR6908274 |
| Pf012_HKCCLCCXY_L3 | Suba | Kenya_West | Kenya | East Africa | 2015 | ERR6908275 |
| Pf013_HK7MKCCXY_L3 | Suba | Kenya_West | Kenya | East Africa | 2015 | ERR6908276 |
| Pf015_HK7LVCCXY_L3 | Suba | Kenya_West | Kenya | East Africa | 2015 | ERR6908277 |
| Pf017_HKCCLCCXY_L3 | Suba | Kenya_West | Kenya | East Africa | 2015 | ERR6908279 |
| Pf018_HK7MKCCXY_L3 | Suba | Kenya_West | Kenya | East Africa | 2015 | ERR6908280 |
| Pf019_HK7MKCCXY_L3 | Suba | Kenya_West | Kenya | East Africa | 2015 | ERR6908281 |
| Pf020_HK7MKCCXY_L3 | Suba | Kenya_West | Kenya | East Africa | 2015 | ERR6908282 |
| Pf021_HKCCLCCXY_L3 | Mfangano | Kenya_West | Kenya | East Africa | 2015 | ERR6908283 |
| Pf023_HK7MKCCXY_L3 | Mfangano | Kenya_West | Kenya | East Africa | 2015 | ERR6908284 |
| Pf024_HKCCLCCXY_L3 | Mfangano | Kenya_West | Kenya | East Africa | 2015 | ERR6908285 |
| Pf025_HKCCLCCXY_L3 | Mfangano | Kenya_West | Kenya | East Africa | 2015 | ERR6908286 |
| Pf026_HK7MKCCXY_L3 | Mfangano | Kenya_West | Kenya | East Africa | 2015 | ERR6908287 |
| Pf028_HKCCLCCXY_L3 | Mfangano | Kenya_West | Kenya | East Africa | 2015 | ERR6908288 |
| Pf029_HKCCLCCXY_L3 | Mfangano | Kenya_West | Kenya | East Africa | 2015 | ERR6908289 |
| QS0056-C | West Arsi Zone | — | Ethiopia | North East Africa | 2013 | ERR1035493 |
| QS0087-C | West Arsi Zone | — | Ethiopia | North East Africa | 2013 | ERR1035520 |
| QS0104-C | West Arsi Zone | — | Ethiopia | North East Africa | 2013 | ERR1035536 |
| QS0109-C | West Arsi Zone | — | Ethiopia | North East Africa | 2013 | ERR1045266 |
| QS0110-C | West Arsi Zone | — | Ethiopia | North East Africa | 2013 | ERR1045267 |
| QS0127-C | West Arsi Zone | — | Ethiopia | North East Africa | 2013 | ERR1045281 |

| Sample | District/Zone | Region | Country | African region | Collection year | Sequence accession |
|---|---|---|---|---|---|---|
| QS0129-C | West Arsi Zone | — | Ethiopia | North East Africa | 2013 | ERR1045283 |
| QS0132-C | West Arsi Zone | — | Ethiopia | North East Africa | 2013 | ERR1045286 |
| QS0135-C | West Arsi Zone | — | Ethiopia | North East Africa | 2013 | ERR1045288 |
| QS0144-C | West Arsi Zone | — | Ethiopia | North East Africa | 2013 | ERR1045295 |
| QS0154-C | Shewa Robit Town Health Centre | — | Ethiopia | North East Africa | 2015 | ERR1106575 |
| QS0159-C | Shewa Robit Town Health Centre | — | Ethiopia | North East Africa | 2015 | ERR1106579 |
| QS0162-C | Shewa Robit Town Health Centre | — | Ethiopia | North East Africa | 2015 | ERR1106581 |
| QS0163-C | Shewa Robit Town Health Centre | — | Ethiopia | North East Africa | 2015 | ERR1106582 |
| QS0169-C | Shewa Robit Town Health Centre | — | Ethiopia | North East Africa | 2015 | ERR1106587 |
| QS0170-C | Shewa Robit Town Health Centre | — | Ethiopia | North East Africa | 2015 | ERR1106590 |
| SPT00898 | Kassala | — | Sudan | North East Africa | 2015 | ERR2299571 |
| SPT00901 | Kassala | — | Sudan | North East Africa | 2015 | ERR2299574 |
| SPT00902 | Kassala | — | Sudan | North East Africa | 2015 | ERR2299575 |
| SPT38368 | Khartoum | — | Sudan | North East Africa | 2017 | ERR3609483 |
| SPT38388 | Khartoum | — | Sudan | North East Africa | 2017 | ERR3609499 |
| SPT38540 | Khartoum | — | Sudan | North East Africa | 2016 | ERR3609275 |
| SPT46076 | Pwani | Tanzania_East | Tanzania | East Africa | 2019 | ERR4021329 |
| SPT46094 | Pwani | Tanzania_East | Tanzania | East Africa | 2019 | ERR4021335 |
| SPT46112 | Pwani | Tanzania_East | Tanzania | East Africa | 2019 | ERR4021297 |
| SPT46113 | Pwani | Tanzania_East | Tanzania | East Africa | 2019 | ERR4021288 |
| SPT46135 | Pwani | Tanzania_East | Tanzania | East Africa | 2019 | ERR4021091 |
| SPT46152 | Pwani | Tanzania_East | Tanzania | East Africa | 2019 | ERR4021344 |

| Sample | District/Zone | Region | Country | African region | Collection year | Sequence accession |
|---|---|---|---|---|---|---|
| SPT46162 | Pwani | Tanzania_East | Tanzania | East Africa | 2019 | ERR4021280 |
| SPT46163 | Pwani | Tanzania_East | Tanzania | East Africa | 2019 | ERR4021333 |
| SPT46226 | Pwani | Tanzania_East | Tanzania | East Africa | 2019 | ERR4020838 |
| SPT46235 | Pwani | Tanzania_East | Tanzania | East Africa | 2019 | ERR4021267 |
| SPT46238 | Pwani | Tanzania_East | Tanzania | East Africa | 2019 | ERR4021236 |
| SPT46265 | Pwani | Tanzania_East | Tanzania | East Africa | 2019 | ERR4020970 |
| SPT46267 | Pwani | Tanzania_East | Tanzania | East Africa | 2019 | ERR4021314 |
| SPT46283 | Pwani | Tanzania_East | Tanzania | East Africa | 2019 | ERR4021046 |
| SPT46300 | Pwani | Tanzania_East | Tanzania | East Africa | 2019 | ERR4021320 |
| SPT46307 | Pwani | Tanzania_East | Tanzania | East Africa | 2019 | ERR4021000 |
| SPT46319 | Pwani | Tanzania_East | Tanzania | East Africa | 2019 | ERR4021013 |
| SPT46323 | Pwani | Tanzania_East | Tanzania | East Africa | 2019 | ERR4020989 |
| SPT46324 | Pwani | Tanzania_East | Tanzania | East Africa | 2019 | ERR4020983 |
| SPT46335 | Pwani | Tanzania_East | Tanzania | East Africa | 2019 | ERR4021050 |
| SPT46350 | Pwani | Tanzania_East | Tanzania | East Africa | 2019 | ERR4021052 |
| SPT46352 | Pwani | Tanzania_East | Tanzania | East Africa | 2019 | ERR4021319 |
| SPT46421 | Pwani | Tanzania_East | Tanzania | East Africa | 2019 | ERR4021070 |
| SPT46457 | Pwani | Tanzania_East | Tanzania | East Africa | 2019 | ERR4288959 |
| SPT46474 | Pwani | Tanzania_East | Tanzania | East Africa | 2019 | ERR4021142 |
| SPT46477 | Pwani | Tanzania_East | Tanzania | East Africa | 2019 | ERR4021149 |
| SPT46482 | Pwani | Tanzania_East | Tanzania | East Africa | 2019 | ERR4021140 |
| SPT46503 | Pwani | Tanzania_East | Tanzania | East Africa | 2019 | ERR4021192 |
| SPT46521 | Pwani | Tanzania_East | Tanzania | East Africa | 2019 | ERR4021201 |
| SPT46555 | Pwani | Tanzania_East | Tanzania | East Africa | 2019 | ERR4288983 |
| SPT53928 | Kilifi | Kenya_East | Kenya | East Africa | 2019 | ERR5839394 |
| SPT53932 | Kilifi | Kenya_East | Kenya | East Africa | 2019 | ERR5839347 |
| SPT53933 | Kilifi | Kenya_East | Kenya | East Africa | 2019 | ERR5839378 |
| SPT53936 | Kilifi | Kenya_East | Kenya | East Africa | 2019 | ERR5839401 |
| SPT53951 | Kilifi | Kenya_East | Kenya | East Africa | 2019 | ERR5839369 |
| SPT53952 | Kilifi | Kenya_East | Kenya | East Africa | 2019 | ERR5839348 |
| SPT53960 | Kilifi | Kenya_East | Kenya | East Africa | 2019 | ERR5839523 |
| SPT66893 | Kilifi | Kenya_East | Kenya | East Africa | 2019 | ERR9515544 |
| SPT66895 | Kilifi | Kenya_East | Kenya | East Africa | 2019 | ERR9515546 |
| SPT66898 | Kilifi | Kenya_East | Kenya | East Africa | 2019 | ERR9515547 |
| SPT66903 | Kilifi | Kenya_East | Kenya | East Africa | 2019 | ERR9515613 |
| SPT66920 | Kilifi | Kenya_East | Kenya | East Africa | 2019 | ERR9515549 |
| SPT66928 | Kilifi | Kenya_East | Kenya | East Africa | 2019 | ERR9515551 |
| SPT66939 | Kilifi | Kenya_East | Kenya | East Africa | 2019 | ERR9515614 |
| SPT66966 | Kilifi | Kenya_East | Kenya | East Africa | 2019 | ERR9515629 |

| Sample | District/Zone | Region | Country | African region | Collection year | Sequence accession |
|---|---|---|---|---|---|---|
| SPT66986 | Kilifi | Kenya_East | Kenya | East Africa | 2019 | ERR9515570 |
| SPT66987 | Kilifi | Kenya_East | Kenya | East Africa | 2019 | ERR9515571 |
| SPT67008 | Kilifi | Kenya_East | Kenya | East Africa | 2020 | ERR9515583 |
| SPT67010 | Kilifi | Kenya_East | Kenya | East Africa | 2020 | ERR9515584 |
| SPT67011 | Kilifi | Kenya_East | Kenya | East Africa | 2020 | ERR9515585 |
| SPT67012 | Kilifi | Kenya_East | Kenya | East Africa | 2020 | ERR9515586 |
| SPT67023 | Kilifi | Kenya_East | Kenya | East Africa | 2019 | ERR9515541 |
| SPT67058 | Kilifi | Kenya_East | Kenya | East Africa | 2019 | ERR9515719 |
| SPT67068 | Kilifi | Kenya_East | Kenya | East Africa | 2019 | ERR9515723 |
| SPT67073 | Kilifi | Kenya_East | Kenya | East Africa | 2019 | ERR9515727 |
| SPT67093 | Kilifi | Kenya_East | Kenya | East Africa | 2019 | ERR9515652 |
| SPT67104 | Kilifi | Kenya_East | Kenya | East Africa | 2019 | ERR9515658 |
| SPT67128 | Kilifi | Kenya_East | Kenya | East Africa | 2019 | ERR9515677 |
| SPT67134 | Kilifi | Kenya_East | Kenya | East Africa | 2019 | ERR9515683 |
| SPT67164 | Kilifi | Kenya_East | Kenya | East Africa | 2019 | ERR9515733 |
| SPT67170 | Kilifi | Kenya_East | Kenya | East Africa | 2019 | ERR9515738 |
| SPT67182 | Kilifi | Kenya_East | Kenya | East Africa | 2019 | ERR9515747 |
| SPT72025 | Kinshasa | — | DRC | Central Africa | 2022 | ERR10685567 |
| SPT72087 | Kinshasa | — | DRC | Central Africa | 2022 | ERR10685613 |
| SPT72089 | Kinshasa | — | DRC | Central Africa | 2022 | ERR10685561 |
| SPT72125 | Kinshasa | — | DRC | Central Africa | 2022 | ERR10685534 |
| SPT72191 | Kinshasa | — | DRC | Central Africa | 2022 | ERR10685509 |
| SPT72247 | Kinshasa | — | DRC | Central Africa | 2022 | ERR10685393 |
| SPT72252 | Kinshasa | — | DRC | Central Africa | 2022 | ERR10685543 |
| SPT72259 | Kinshasa | — | DRC | Central Africa | 2022 | ERR10685389 |
| SPT72271 | Kinshasa | — | DRC | Central Africa | 2022 | ERR10685408 |
| SPT72285 | Kinshasa | — | DRC | Central Africa | 2022 | ERR10685383 |
| SPT72286 | Kinshasa | — | DRC | Central Africa | 2022 | ERR10685379 |
| SPT72309 | Kinshasa | — | DRC | Central Africa | 2022 | ERR10763977 |

| Sample | District/Zone | Region | Country | African region | Collection year | Sequence accession |
|---|---|---|---|---|---|---|
| SPT72311 | Kinshasa | — | DRC | Central Africa | 2022 | ERR10763968 |
| SPT72328 | Kinshasa | — | DRC | Central Africa | 2022 | ERR10763976 |
| SPT72337 | Kinshasa | — | DRC | Central Africa | 2022 | ERR10763954 |
| SPT72340 | Kinshasa | — | DRC | Central Africa | 2022 | ERR10763926 |
| SPT72361 | Kinshasa | — | DRC | Central Africa | 2022 | ERR10763961 |
| SPT72367 | Kinshasa | — | DRC | Central Africa | 2022 | ERR10763960 |
| SPT72373 | Kinshasa | — | DRC | Central Africa | 2022 | ERR10763930 |
| SPT72389 | Kinshasa | — | DRC | Central Africa | 2022 | ERR10763985 |
| SPT72408 | Kinshasa | — | DRC | Central Africa | 2022 | ERR10764010 |
| SPT72418 | Kinshasa | — | DRC | Central Africa | 2022 | ERR10764002 |
| SPT72421 | Kinshasa | — | DRC | Central Africa | 2022 | ERR10764007 |
| SPT72427 | Kinshasa | — | DRC | Central Africa | 2022 | ERR10763979 |
| SPT87699 | Kinshasa | — | DRC | Central Africa | 2022 | ERR11840555 |
| SPT87702 | Kinshasa | — | DRC | Central Africa | 2022 | ERR11840552 |
| SPT87729 | Kinshasa | — | DRC | Central Africa | 2022 | ERR11840541 |
| SPT87737 | Kinshasa | — | DRC | Central Africa | 2022 | ERR11840534 |
| SPT87745 | Kinshasa | — | DRC | Central Africa | 2022 | ERR11840571 |
| SPT87977 | Kinshasa | — | DRC | Central Africa | 2022 | ERR11840472 |
| SPT87979 | Kinshasa | — | DRC | Central Africa | 2022 | ERR11840470 |
| SPT87986 | Kinshasa | — | DRC | Central Africa | 2022 | ERR11840480 |
| SPT87995 | Kinshasa | — | DRC | Central Africa | 2022 | ERR11840502 |
| SPT88010 | Kinshasa | — | DRC | Central Africa | 2022 | ERR11840492 |

| Sample | District/Zone | Region | Country | African region | Collection year | Sequence accession |
|---|---|---|---|---|---|---|
| SPT88018 | Kinshasa | — | DRC | Central Africa | 2022 | ERR11840425 |
| SPT88048 | Kinshasa | — | DRC | Central Africa | 2022 | ERR11840439 |
| SPT88145 | Kinshasa | — | DRC | Central Africa | 2022 | ERR11840798 |
| SPT88148 | Kinshasa | — | DRC | Central Africa | 2022 | ERR11840801 |
| SPT88181 | Kinshasa | — | DRC | Central Africa | 2022 | ERR11840843 |
| SPT88273 | Kinshasa | — | DRC | Central Africa | 2022 | ERR11840835 |
| SPT88310 | Kinshasa | — | DRC | Central Africa | 2022 | ERR11840901 |
| SPT88323 | Kinshasa | — | DRC | Central Africa | 2022 | ERR11840904 |
| SPT88327 | Kinshasa | — | DRC | Central Africa | 2022 | ERR11840870 |
| SPT88348 | Kinshasa | — | DRC | Central Africa | 2022 | ERR11840882 |
| SPT88393 | Kinshasa | — | DRC | Central Africa | 2022 | ERR11840920 |
| SPT88398 | Kinshasa | — | DRC | Central Africa | 2022 | ERR11840876 |
| SPT88452 | Kinshasa | — | DRC | Central Africa | 2022 | ERR11840952 |
| SPT88454 | Kinshasa | — | DRC | Central Africa | 2022 | ERR11840951 |
| SPT88514 | Kinshasa | — | DRC | Central Africa | 2022 | ERR11871804 |
| SPT88563 | Kinshasa | — | DRC | Central Africa | 2022 | ERR11871853 |
| SPT88578 | Kinshasa | — | DRC | Central Africa | 2022 | ERR11871830 |
| SPT88693 | Kinshasa | — | DRC | Central Africa | 2022 | ERR11871721 |
| SPT88715 | Kinshasa | — | DRC | Central Africa | 2022 | ERR11871752 |
| SPT88722 | Kinshasa | — | DRC | Central Africa | 2022 | ERR11871759 |
| SPT88723 | Kinshasa | — | DRC | Central Africa | 2022 | ERR11871760 |
| SPT88727 | Kinshasa | — | DRC | Central Africa | 2022 | ERR11871775 |

| Sample | District/Zone | Region | Country | African region | Collection year | Sequence accession |
|---|---|---|---|---|---|---|
| SPT88750 | Kinshasa | — | DRC | Central Africa | 2022 | ERR11871781 |
| SPT88755 | Kinshasa | — | DRC | Central Africa | 2022 | ERR11871786 |
| SPT88760 | Kinshasa | — | DRC | Central Africa | 2022 | ERR11871771 |
| T0907AD0 | Karagwe | Tanzania_West | Tanzania | East Africa | 2022 | SRR29397016 |
| T0909BD0 | Karagwe | Tanzania_West | Tanzania | East Africa | 2022 | SRR29397011 |
| T0911AD0 | Karagwe | Tanzania_West | Tanzania | East Africa | 2022 | SRR29397007 |
| T0913AD0 | Karagwe | Tanzania_West | Tanzania | East Africa | 2022 | SRR29397003 |
| T0918BD0 | Karagwe | Tanzania_West | Tanzania | East Africa | 2022 | SRR29396972 |
| T0919AD0 | Karagwe | Tanzania_West | Tanzania | East Africa | 2022 | SRR29396971 |
| T0919BD0 | Karagwe | Tanzania_West | Tanzania | East Africa | 2022 | SRR29396970 |
| T0922AD0 | Karagwe | Tanzania_West | Tanzania | East Africa | 2022 | SRR29396964 |
| T0922BD0 | Karagwe | Tanzania_West | Tanzania | East Africa | 2022 | SRR29396963 |
| T0923AD0 | Karagwe | Tanzania_West | Tanzania | East Africa | 2022 | SRR29396962 |
| T0925BD0 | Karagwe | Tanzania_West | Tanzania | East Africa | 2022 | SRR29396998 |
| T0926AD0 | Karagwe | Tanzania_West | Tanzania | East Africa | 2022 | SRR29396997 |
| T0926BD0 | Karagwe | Tanzania_West | Tanzania | East Africa | 2022 | SRR29396995 |
| T0928AD0 | Karagwe | Tanzania_West | Tanzania | East Africa | 2022 | SRR29396992 |
| T0930AD0 | Karagwe | Tanzania_West | Tanzania | East Africa | 2022 | SRR29396990 |
| T0932AD0 | Karagwe | Tanzania_West | Tanzania | East Africa | 2022 | SRR29396987 |
| T0937BD0 | Karagwe | Tanzania_West | Tanzania | East Africa | 2022 | SRR29396982 |
| T0938AD0 | Karagwe | Tanzania_West | Tanzania | East Africa | 2022 | SRR29396961 |
| T0943BD0 | Karagwe | Tanzania_West | Tanzania | East Africa | 2022 | SRR29396958 |
| T0944AD0 | Karagwe | Tanzania_West | Tanzania | East Africa | 2022 | SRR29396957 |
| T0945AD0 | Karagwe | Tanzania_West | Tanzania | East Africa | 2022 | SRR29396956 |
| T0946BD0 | Karagwe | Tanzania_West | Tanzania | East Africa | 2022 | SRR29396955 |
| T0949BD0 | Karagwe | Tanzania_West | Tanzania | East Africa | 2022 | SRR29396953 |
| T0955BD0 | Karagwe | Tanzania_West | Tanzania | East Africa | 2022 | SRR29396951 |
| T0956BD0 | Karagwe | Tanzania_West | Tanzania | East Africa | 2022 | SRR29396950 |
| T0961AD0 | Karagwe | Tanzania_West | Tanzania | East Africa | 2022 | SRR29396948 |
| T0962AD0 | Karagwe | Tanzania_West | Tanzania | East Africa | 2022 | SRR29396947 |
| T0967BD0 | Karagwe | Tanzania_West | Tanzania | East Africa | 2022 | SRR29396944 |
| T0968BD0 | Karagwe | Tanzania_West | Tanzania | East Africa | 2022 | SRR29396942 |
| T0973AD0 | Karagwe | Tanzania_West | Tanzania | East Africa | 2022 | SRR29396941 |
| T0980AD0 | Karagwe | Tanzania_West | Tanzania | East Africa | 2022 | SRR29396939 |
| T0983BD0 | Karagwe | Tanzania_West | Tanzania | East Africa | 2022 | SRR29396936 |
| T0987AD0 | Karagwe | Tanzania_West | Tanzania | East Africa | 2022 | SRR29396934 |
| UGK_707_3-SRX146743 | Kampala | Uganda_East-Past | Uganda | East Africa | — | SRR641960 |

| Sample | District/Zone | Region | Country | African region | Collection year | Sequence accession |
|---|---|---|---|---|---|---|
| Uganda-AG-04-11 | Agago | Uganda_North-Past | Uganda | East Africa | 2019 | SRR34783618 |
| Uganda-AG-04-13 | Agago | Uganda_North-Past | Uganda | East Africa | 2019 | SRR34783538 |
| Uganda-AG-05-15 | Agago | Uganda_North-Past | Uganda | East Africa | 2020 | SRR34783511 |
| Uganda-AG-05-22 | Agago | Uganda_North-Past | Uganda | East Africa | 2020 | SRR34783500 |
| Uganda-AG-05-41 | Agago | Uganda_North-Past | Uganda | East Africa | 2020 | SRR34783595 |
| Uganda-AG-05-47 | Agago | Uganda_North-Past | Uganda | East Africa | 2020 | SRR34783584 |
| Uganda-AG-05-49 | Agago | Uganda_North-Past | Uganda | East Africa | 2020 | SRR34783562 |
| Uganda-AG-05-56 | Agago | Uganda_North-Past | Uganda | East Africa | 2020 | SRR34783553 |
| Uganda-AG-05-59 | Agago | Uganda_North-Past | Uganda | East Africa | 2020 | SRR34783552 |
| Uganda-AG-05-81 | Agago | Uganda_North-Past | Uganda | East Africa | 2020 | SRR34783548 |
| Uganda-BUS-025 | Mbale | Uganda_East-Past | Uganda | East Africa | 2021 | SRR34783542 |
| Uganda-BUS-041 | Mbale | Uganda_East-Past | Uganda | East Africa | 2021 | SRR34783540 |
| Uganda-KBG-04-05 | Kaabong | Uganda_North-East-Past | Uganda | East Africa | 2019 | SRR34783536 |
| Uganda-KBG-05-65 | Kaabong | Uganda_North-East-Past | Uganda | East Africa | 2020 | SRR34783482 |
| Uganda-KBG-05-73 | Kaabong | Uganda_North-East-Past | Uganda | East Africa | 2020 | SRR34783480 |
| Uganda-KBG-05-84 | Kaabong | Uganda_North-East-Past | Uganda | East Africa | 2020 | SRR34783478 |
| Uganda-KO-05-14 | Kole | Uganda_North-Past | Uganda | East Africa | 2020 | SRR34783477 |
| Uganda-KO-05-75 | Kole | Uganda_North-Past | Uganda | East Africa | 2020 | SRR34783471 |
| Uganda-KTK-04-27 | Katakwi | Uganda_North-East-Past | Uganda | East Africa | 2019 | SRR34783466 |
| Uganda-KTK-04-45 | Katakwi | Uganda_North-East-Past | Uganda | East Africa | 2019 | SRR34783464 |
| Uganda-KTK-05-20 | Katakwi | Uganda_North-East-Past | Uganda | East Africa | 2020 | SRR34783462 |
| Uganda-KTK-05-21 | Katakwi | Uganda_North-East-Past | Uganda | East Africa | 2020 | SRR34783535 |
| Uganda-KTK-05-22 | Katakwi | Uganda_North-East-Past | Uganda | East Africa | 2020 | SRR34783534 |
| Uganda-KTK-05-45 | Katakwi | Uganda_North-East-Past | Uganda | East Africa | 2020 | SRR34783531 |
| Uganda-KTK-05-55 | Katakwi | Uganda_North-East-Past | Uganda | East Africa | 2020 | SRR34783529 |
| Uganda-KTK-05-56 | Katakwi | Uganda_North-East-Past | Uganda | East Africa | 2020 | SRR34783528 |
| Uganda-KTK-05-77 | Katakwi | Uganda_North-East-Past | Uganda | East Africa | 2020 | SRR34783526 |
| Uganda-KTK-05-79 | Katakwi | Uganda_North-East-Past | Uganda | East Africa | 2020 | SRR34783525 |

| Sample | District/Zone | Region | Country | African region | Collection year | Sequence accession |
|---|---|---|---|---|---|---|
| Uganda-KTK-05-97 | Katakwi | Uganda_North-East-Past | Uganda | East Africa | 2020 | SRR34783524 |
| Uganda-LA-02-47 | Lamwo | Uganda_North-Past | Uganda | East Africa | 2017 | SRR34783516 |
| Uganda-LA-05-10 | Lamwo | Uganda_North-Past | Uganda | East Africa | 2020 | SRR34783510 |
| Uganda-LA-05-17 | Lamwo | Uganda_North-Past | Uganda | East Africa | 2020 | SRR34783509 |
| Uganda-LA-05-25 | Lamwo | Uganda_North-Past | Uganda | East Africa | 2020 | SRR34783507 |
| Uganda-LA-05-30 | Lamwo | Uganda_North-Past | Uganda | East Africa | 2020 | SRR34783506 |
| Uganda-LA-05-39 | Lamwo | Uganda_North-Past | Uganda | East Africa | 2020 | SRR34783504 |
| Uganda-LA-05-52 | Lamwo | Uganda_North-Past | Uganda | East Africa | 2020 | SRR34783502 |
| Uganda-LA-05-59 | Lamwo | Uganda_North-Past | Uganda | East Africa | 2020 | SRR34783497 |
| Uganda-LA-05-66 | Lamwo | Uganda_North-Past | Uganda | East Africa | 2020 | SRR34783496 |
| Uganda-LA-05-75 | Lamwo | Uganda_North-Past | Uganda | East Africa | 2020 | SRR34783493 |
| Uganda-LA-05-79 | Lamwo | Uganda_North-Past | Uganda | East Africa | 2020 | SRR34783492 |
| Uganda-LA-05-89 | Lamwo | Uganda_North-Past | Uganda | East Africa | 2020 | SRR34783489 |
| Uganda-PAT-005-P2-D7 | Agago | Uganda_North-Past | Uganda | East Africa | 2021 | SRR34783609 |
| Uganda-PAT-011-P2-A3 | Agago | Uganda_North-Past | Uganda | East Africa | 2021 | SRR34783603 |
| Uganda-PAT-015-P2-H4 | Agago | Uganda_North-Past | Uganda | East Africa | 2021 | SRR34783600 |
| Uganda-PAT-018 | Agago | Uganda_North-Past | Uganda | East Africa | 2021 | SRR34783597 |
| Uganda-PAT-019-P2-F6 | Agago | Uganda_North-Past | Uganda | East Africa | 2021 | SRR34783596 |
| Uganda-PAT-020-P2-H6 | Agago | Uganda_North-Past | Uganda | East Africa | 2021 | SRR34783594 |
| Uganda-PAT-024-P2-B10 | Agago | Uganda_North-Past | Uganda | East Africa | 2021 | SRR34783592 |
| Uganda-PAT-039 | Agago | Uganda_North-Past | Uganda | East Africa | 2021 | SRR34783585 |
| Uganda-PAT-040 | Agago | Uganda_North-Past | Uganda | East Africa | 2021 | SRR34783583 |
| Uganda-PAT-043 | Agago | Uganda_North-Past | Uganda | East Africa | 2021 | SRR34783582 |
| Uganda-PAT-045 | Agago | Uganda_North-Past | Uganda | East Africa | 2021 | SRR34783581 |
| Uganda-PAT-046 | Agago | Uganda_North-Past | Uganda | East Africa | 2021 | SRR34783580 |
| Uganda-PAT-048 | Agago | Uganda_North-Past | Uganda | East Africa | 2021 | SRR34783578 |
| Uganda-PAT-052 | Agago | Uganda_North-Past | Uganda | East Africa | 2021 | SRR34783577 |
| Uganda-PAT-071 | Agago | Uganda_North-Past | Uganda | East Africa | 2022 | SRR34783570 |
| Uganda-PBC-261-12Feb21 | Tororo | Uganda_East-Past | Uganda | East Africa | 2021 | SRR34783564 |
| Uganda-TDH-173-P2-B11 | Tororo | Uganda_East-Past | Uganda | East Africa | 2021 | SRR34783558 |
| Uganda-TDH-174 | Tororo | Uganda_East-Past | Uganda | East Africa | 2021 | SRR34783557 |
| Uganda-TDH-190-P2-H2 | Tororo | Uganda_East-Past | Uganda | East Africa | 2021 | SRR34783556 |
| Uganda-TDH-191-P2-G8 | Tororo | Uganda_East-Past | Uganda | East Africa | 2021 | SRR34783555 |
| Uganda_Kampala1-SRX006181 | Kampala | Uganda_East-Past | Uganda | East Africa | 2006 | SRR340211, SRR340216 |
| Uganda_Kampala2-SRX006665 | Kampala | Uganda_East-Past | Uganda | East Africa | 2007 | SRR340214, SRR340220 |
| Uganda_Kampala3-SRX006734 | Kampala | Uganda_East-Past | Uganda | East Africa | 2006 | SRR340224, SRR340225 |

| Sample | District/Zone | Region | Country | African region | Collec tion year | Sequence accession |
|---|---|---|---|---|---|---|
| Uganda_Kampala4-SRX006178 | Kampala | Uganda_East-Past | Uganda | East Africa | 2006 | SRR340208, SRR340221 |
| Uganda_Kampala7-SRX007630 | Kampala | Uganda_East-Past | Uganda | East Africa | 2007 | SRR340213, SRR340218 |
| Uganda_Kampala9-SRX006735 | Kampala | Uganda_East-Past | Uganda | East Africa | 2007 | SRR340215, SRR340226 |
| CW_002_lib666283 | Bungoma | Kenya_West | Kenya | East Africa | 2018 | ERR15534969 |
| CW_003_lib666284 | Bungoma | Kenya_West | Kenya | East Africa | 2018 | ERR15534970 |
| CW_004_lib666285 | Bungoma | Kenya_West | Kenya | East Africa | 2018 | ERR15534971 |
| CW_006_lib666287 | Bungoma | Kenya_West | Kenya | East Africa | 2018 | ERR15534972 |
| CW_024_lib666305 | Bungoma | Kenya_West | Kenya | East Africa | 2018 | ERR15534989 |
| CW_030_lib666311 | Bungoma | Kenya_West | Kenya | East Africa | 2018 | ERR15534995 |
| CW_034_lib666315 | Bungoma | Kenya_West | Kenya | East Africa | 2018 | ERR15534999 |
| CW_042_lib666323 | Bungoma | Kenya_West | Kenya | East Africa | 2018 | ERR15535007 |
| CW_045_lib666326 | Bungoma | Kenya_West | Kenya | East Africa | 2018 | ERR15535010 |
| CW_046_lib666327 | Bungoma | Kenya_West | Kenya | East Africa | 2018 | ERR15535011 |

**Note:** Metadata for external comparator samples and historical Ugandan samples used in the population-genetic analyses. Sample denotes the study sample identifier; District/Zone and region provide the available sub-national location or analytical grouping; African region and continent give broad geographic classification; collection year gives the year of sampling; and sequence accession is the public archive accession. An em dash indicates information that was not available in the source metadata. Groups ending in "-Past" denote historical Ugandan sample sets.

**Supplementary Table 3. Country-wide distribution of alternative amino-acid alleles in pfpx1 (PF3D7_0720700) in Uganda**

| Gene | Gene ID | Reference AA | AA position | Alternative AA | Samples with allele, n | PIN-positive samples, n | Samples assessed at position, n | Allele prevalence | PIN-positive fraction | Variant ID |
|---|---|---|---|---|---|---|---|---|---|---|
| PX1 | PF3D7_0720700 | D | 384 | A | 98 | 97 | 247 | 0.3968 | 0.9898 | D384A |
| PX1 | PF3D7_0720700 | D | 384 | G | 8 | 2 | 247 | 0.0324 | 0.2500 | D384G |
| PX1 | PF3D7_0720700 | D | 384 | Y | 2 | 2 | 247 | 0.0081 | 1.0000 | D384Y |
| PX1 | PF3D7_0720700 | Y | 1006 | C | 4 | 1 | 244 | 0.0164 | 0.2500 | Y1006C |
| PX1 | PF3D7_0720700 | S | 1211 | R | 16 | 0 | 244 | 0.0656 | 0.0000 | S1211R |
| PX1 | PF3D7_0720700 | L | 1222 | P | 177 | 166 | 244 | 0.7254 | 0.9379 | L1222P |
| PX1 | PF3D7_0720700 | C | 1484 | F | 2 | 0 | 236 | 0.0085 | 0.0000 | C1484F |
| PX1 | PF3D7_0720700 | C | 1484 | Y | 4 | 4 | 236 | 0.0169 | 1.0000 | C1484Y |
| PX1 | PF3D7_0720700 | D | 1502 | E | 3 | 0 | 236 | 0.0127 | 0.0000 | D1502E |
| PX1 | PF3D7_0720700 | L | 1570 | V | 7 | 0 | 236 | 0.0297 | 0.0000 | L1570V |
| PX1 | PF3D7_0720700 | E | 1591 | K | 4 | 0 | 236 | 0.0169 | 0.0000 | E1591K |
| PX1 | PF3D7_0720700 | S | 1673 | N | 211 | 160 | 236 | 0.8941 | 0.7583 | S1673N |
| PX1 | PF3D7_0720700 | V | 1686 | M | 2 | 0 | 236 | 0.0085 | 0.0000 | V1686M |
| PX1 | PF3D7_0720700 | M | 1701 | I | 166 | 160 | 236 | 0.7034 | 0.9639 | M1701I |
| PX1 | PF3D7_0720700 | D | 1705 | N | 166 | 160 | 236 | 0.7034 | 0.9639 | D1705N |
| PX1 | PF3D7_0720700 | I | 1707 | M | 3 | 0 | 236 | 0.0127 | 0.0000 | I1707M |
| PX1 | PF3D7_0720700 | D | 1711 | N | 36 | 0 | 236 | 0.1525 | 0.0000 | D1711N |
| PX1 | PF3D7_0720700 | D | 1889 | G | 4 | 2 | 239 | 0.0167 | 0.5000 | D1889G |
| PX1 | PF3D7_0720700 | D | 1889 | N | 3 | 3 | 239 | 0.0126 | 1.0000 | D1889N |
| PX1 | PF3D7_0720700 | G | 1891 | K | 2 | 2 | 238 | 0.0084 | 1.0000 | G1891K |
| PX1 | PF3D7_0720700 | T | 1903 | A | 5 | 0 | 236 | 0.0212 | 0.0000 | T1903A |
| PX1 | PF3D7_0720700 | R | 1951 | I | 5 | 0 | 236 | 0.0212 | 0.0000 | R1951I |
| PX1 | PF3D7_0720700 | E | 2160 | K | 3 | 0 | 249 | 0.0120 | 0.0000 | E2160K |

**Note:** Country-wide counts and proportions of non-synonymous alternative amino-acid alleles detected in PfPX11. Reference AA and AA position identify the reference residue and its protein position; Alternative AA is the observed replacement residue; Samples with allele is the number of samples carrying the alternative allele; PIN-positive samples is the number of allele-carrying samples classified as PIN-positive in the source analysis; Samples assessed at position is the denominator with callable data at that position; Allele prevalence = Samples with allele / Samples assessed at position; PIN-positive fraction = PIN-positive samples / Samples with allele; and Variant ID combines the reference residue, amino-acid position, and alternative residue. AA, amino acid; n, number of samples.
